# Structural basis of sulfide production in dissimilatory sulfur metabolism

**DOI:** 10.64898/2026.08.25.746951

**Authors:** Max Dongsheng Yin, Raquel M. Bernardino, Ana C. C. Barbosa, José A. Brito, Andreia I. Pimenta, Sonja Welsch, Filipa L. Sousa, Inês A. C. Pereira, Bonnie J. Murphy

## Abstract

The membrane-bound DsrMK(JOP) complex is central to dissimilatory sulfur metabolism, including in sulfate-reducing microbes (SRM), a group that plays important roles in shaping planetary and human health. Despite this global importance, the mechanism of sulfide production and its links to energy conservation remain unclear. Here, we present high-resolution cryo-EM structures of DsrMKJOP from *Archaeoglobus fulgidus*, alone, with menadiol and with the sulfur-carrying substrate DsrC-trisulfide, complemented by physiological and biochemical studies. The results clarify how SRM control the reactivity of sulfur to selectively achieve sulfide production. While DsrC-trisulfide is highly stable in isolation, interaction with the DsrK subunit facilitates its hydrolytic activation, triggering a conformational change. This brings a perthiosulfenate sulfur intermediate into the catalytic pocket of DsrK for reduction at a single non-cubane [4Fe-4S] cluster, likely supported by a conserved non-ligating cysteine. DsrM harbors a structural quinone-binding site, but seems not to catalyze menaquinol oxidation, although this likely occurs in DsrMK complexes from different sulfur-metabolizing organisms. In DsrMKJOP, trisulfide reduction by DsrK is linked to quinol oxidation at DsrP, releasing protons to the periplasm to generate a proton-motive force.

## Introduction

The microbial conversion of inorganic sulfur compounds is thought to be among the earliest processes sustaining life on Earth^1^, with sulfite reduction likely present already in the last universal common ancestor ^2,3^. The metabolic pathway that derives energy from sulfite reduction to sulfide, known as dissimilatory sulfite reduction (DSR), is also central to the metabolism of other oxidized sulfur compounds including sulfate, thiosulfate, and organosulfonates as terminal electron acceptors for anaerobic respiration. These respiratory processes play crucial roles in driving the global sulfur and carbon biogeochemical cycles ^4,5^, including through major impacts on methane production and uptake^6^, and therefore on global climate. Microbial sulfide production in the gut via DSR is also important in human and animal health, where it is associated with inflammatory bowel diseases and colorectal cancer ^7–9^.

Although metabolic pathways converting sulfur compounds are widespread and diverse, a few central complexes are present in all organisms performing DSR. In spite of their central role, a molecular understanding of the reaction mechanisms of some of the complexes is still missing. The dissimilatory sulfite reductase DsrAB, a key enzyme in sulfur metabolism, reduces sulfite to zero-valent sulfur (S^0^) and binds it to a protein substrate, DsrC, as a trisulfide bridge between two strictly conserved cysteine residues near the C-terminus of DsrC ^10^. This S^0^ substrate is subsequently reduced to sulfide (H_2_S) by the membrane-bound DsrMKJOP complex via menaquinol oxidation ^11^. Sulfite reduction is coupled to the generation of a proton motive force (pmf) ^12,13^, and as one of only two conserved transmembrane redox complexes in SRM ^14^, DsrMKJOP is thought to mediate this coupling ^5,11^. Such coupling likely requires tight control over the reduction step. Consistent with this, the DsrC-trisulfide exhibits remarkable resistance to chemical reductants, including dithiothreitol and the strong reductant sodium borohydride, in the absence of DsrMKJOP ^11^, suggesting that its reduction is controlled by specific protein– protein interactions. Despite the important role of DsrMKJOP in sulfur metabolism, its structure is unknown and key aspects of its function remain unresolved.

A major open question concerns the architecture and modular organization of the complex. DsrMKJOP likely evolved from a simpler ancestral core, DsrMK, with the additional subunits DsrJOP acquired later ^15^. DsrJOP proteins are missing in several DSR-utilizing organisms ^5,15^, and their functional contributions are unclear. Both membrane subunits DsrM and DsrP belong to separate quinone-binding protein families ^16^ involved in energy conservation ^17,18^, but it is unknown why two quinone-binding subunits are required. The periplasmic subunits DsrJ and DsrO — predicted to bind hemes *c* and [4Fe-4S] clusters, respectively — also lack functional assignment. Fundamentally, it remains to be clarified how electrons flow through DsrMKJOP and how this process is connected to pmf generation.

Another unresolved question is how sulfur chemistry is precisely controlled during DsrC-trisulfide reduction. Sulfur exhibits highly flexible chemistry, so that the reactivity of sulfur-based intermediates must be closely controlled to allow coupling to energy conservation. This control is exemplified by the fact that DsrC-trisulfide is so inert to reduction in isolation, but can be reduced at DsrMKJOP by the relatively mild electron donor menaquinol ^11^, suggesting that protein-protein interactions are important for activating the DsrC-trisulfide for reduction. The reaction is believed to be catalyzed by the cytoplasmic subunit DsrK, given its homology to HdrBC and HdrD, the catalytic subunits of methanogenic heterodisulfide reductases ^14,19^, but how DsrMKJOP enables reduction of DsrC-trisulfide is unclear. Hdr enzymes use two unique non-cubane [4Fe-4S] clusters to catalyze the homolytic two-electron reduction of heterodisulfide ^20,21^, whereas DsrK contains a binding motif for only one such cluster ^14,19^ and must deliver four electrons to reduce the DsrC-trisulfide, yielding sulfide and DsrC in its dithiol form ^11^. Thus, the proposed Hdr mechanism does not provide a clear template for the reaction catalyzed by DsrK. So far, no structure of an Hdr-related protein with a single non-cubane cluster has been reported.

To address these questions, we determined the first high-resolution structures of DsrMKJOP, using cryogenic electron microscopy (cryo-EM). We characterize the quinol binding sites in DsrM and DsrP, and provide evidence for electron transfer between the distant DsrP site and DsrK. We also capture states of the DsrMKJOP in complex with its substrate, DsrC-trisulfide, that reveal how this is activated for reduction. Our data suggest that hydrolytic activation of the DsrC-trisulfide precedes sulfur reduction. Overall, these findings provide key mechanistic insights into sulfur chemistry during DSR, a pathway with strong relevance to environmental and human health.

### Architecture of DsrMKJOP

We purified DsrMKJOP solubilized in n-dodecyl-β-D-maltoside from *Archaeoglobus fulgidus* VC-16 (*A. fulgidus*) (fig. S1) and confirmed its menadiol:DsrC-trisulfide oxidoreductase activity ^11^. Cryo-EM structures determined by single-particle analysis reached 2.0 Å resolution and revealed a dimer of DsrMKJOP heteropentamers (Fig. 1A). In addition to an “as-isolated” sample, we also prepared samples of DsrMKJOP stably reduced with menadiol (“reduced” sample), incubated with its substrate DsrC-trisulfide (“DsrC-trisulfide” sample), and reduced with menadiol followed by reaction with DsrC-trisulfide (“turnover” sample) (figs. S2-9, tables S1-3; see methods).

**Fig. 1.**
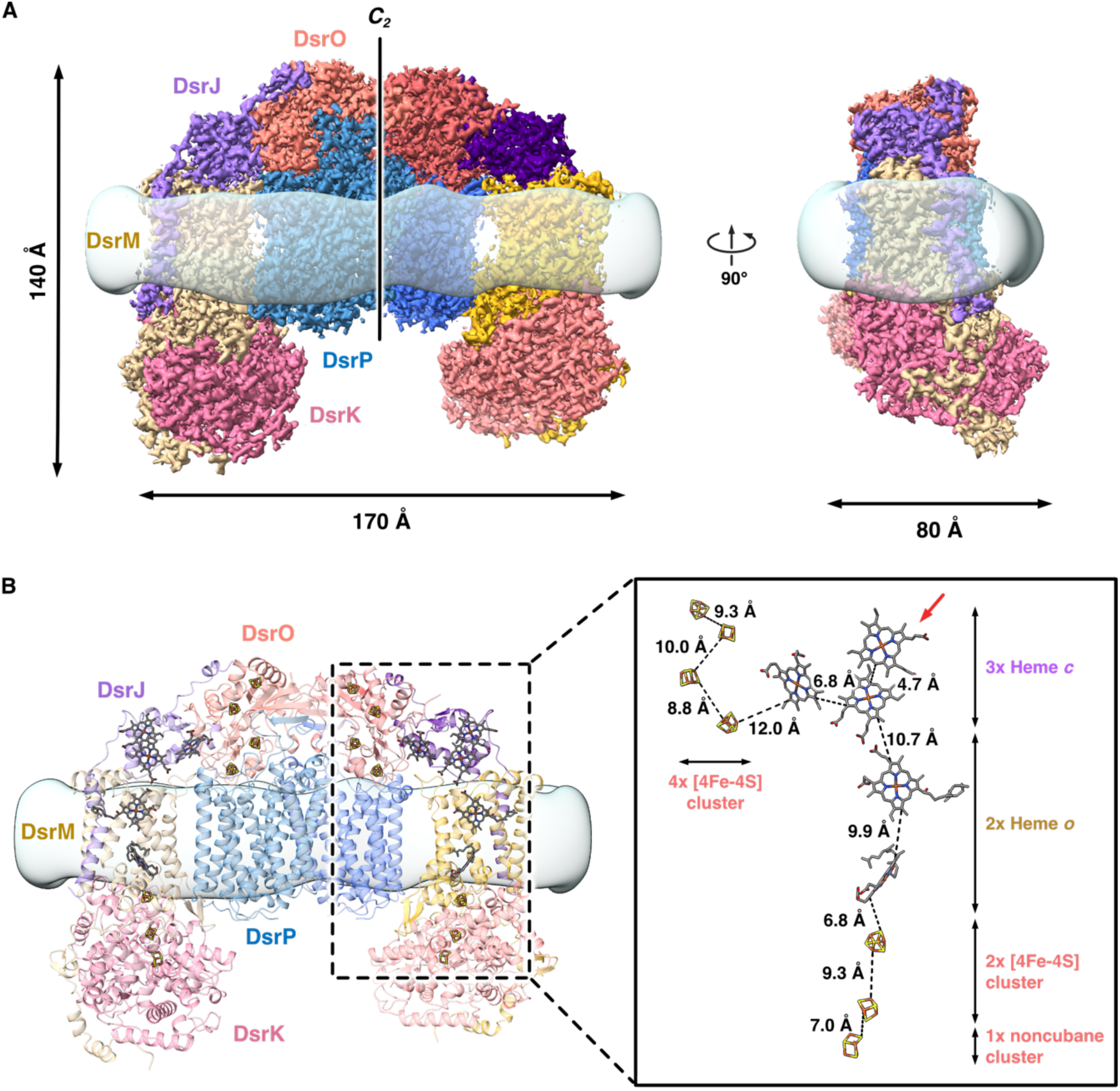
Overall structure of DsrMKJOP. **(A)** Cryo-EM map of the “as-isolated” sample contoured at 1 σ. **(B)** The refined model of the complex, shown as ribbons. To better visualize micelle density, a transparent overlay was applied in (A) and (B), showing the micelle volume Gaussian-filtered to 6 Å and contoured at 1.3 σ. Inset shows cofactors positioned for efficient electron transfer, with edge-to-edge distances indicated by dashed lines. The unusual Cys/His-coordinated heme in DsrJ is marked by a red arrow. Cofactors are shown as sticks and colored as follows: carbon - gray; oxygen - red; nitrogen - blue; sulfur - yellow; and iron - orange.

All redox cofactors are well resolved and positioned for efficient electron transfer (Fig. 1B). Membrane-embedded DsrM contains two hemes *o* that were predicted to be hemes *b* ^14^. The DsrMK subcomplex, strictly conserved in all organisms carrying out DSR, is homologous to the membrane-bound heterodisulfide reductase HdrED of methanogens ^22–24^. Our structure confirms a close association of DsrM with DsrK, which binds two canonical [4Fe-4S] clusters and one non-cubane cluster ^14,19^. DsrM also contacts the triheme cytochrome *c* DsrJ on its periplasmic face and, to a lesser extent, the iron-sulfur protein DsrO and the membrane-bound DsrP. Two hemes *c* in DsrJ exhibit bis-His coordination, while the third shows an unusual Cys/His coordination ^14^, which has been observed for catalytic hemes in other systems, including for thiosulfate oxidation in TsdA/SoxAX ^25,26^, but also for electron transfer hemes. Previously DsrMKJOP was shown not to catalyze thiosulfate or sulfite oxidation ^11^. We tested whether substitution of the axial cysteine ligand affected growth in a genetically tractable SRM (tables S4-5; see methods). These point mutations did not affect sulfate or sulfite respiration, suggesting that DsrJ does not play a catalytic role, at least during growth on these sulfur compounds. In contrast, *dsrJ* and *dsrJOP* deletions abolished growth on both sulfate and sulfite (fig. S10). DsrO binds four [4Fe-4S] clusters and contributes an axial histidine ligand (His148) to a DsrJ heme. DsrP lacks intrinsic cofactors and interacts primarily with DsrO. DsrP-DsrP and DsrO-DsrO contacts at the *C_2_* symmetry axis stabilize the decamer.

### DsrM binds a structural menaquinone but lacks features for quinol oxidation

The membrane-bound subunit DsrM shows density for a copurified menaquinone-7 (MK-7) between the two hemes *o* (Fig. 2A-C). MK-7 is located in a lipid-exposed hydrophobic cavity and surrounded by nonpolar residues from both DsrM and DsrJ (figs. S11 and S12A), and we could not identify a clear pathway for protons from this site to either side of the membrane. No proton-accepting/-donating groups directly coordinate the quinone, suggesting that MK-7 may play a structural role rather than serving as electron donor. This interpretation is supported by its copurification with the complex (no MK-7 was added at any step of the purification; see methods) and by its persistence in structures determined under menadiol-reduced and “turnover” conditions, suggesting very slow exchange kinetics. A DsrM homolog, the nitrate reductase subunit NarI, presents a similar lipid-exposed cavity that was previously proposed to bind a structural quinone, though none was observed structurally ^27^. A structure of NarI bound to its competitive inhibitor pentachlorophenol (PCP) revealed the likely site for quinol oxidation at a distinct location close to the heme on the periplasmic side of the protein ^27^ (fig. S12B). The equivalent site in DsrM cannot bind a quinol, as it is blocked by two conserved arginines (fig. S13) and a short helical segment of the transmembrane helix (TMH) 3, separated from the rest of TMH3 by a loop insertion (fig. S14). In NarI, this loop insertion is absent, so that TMH3 is continuous, and glycine residues replace the arginines seen in DsrM (figs. S13-14).

**Fig. 2.**
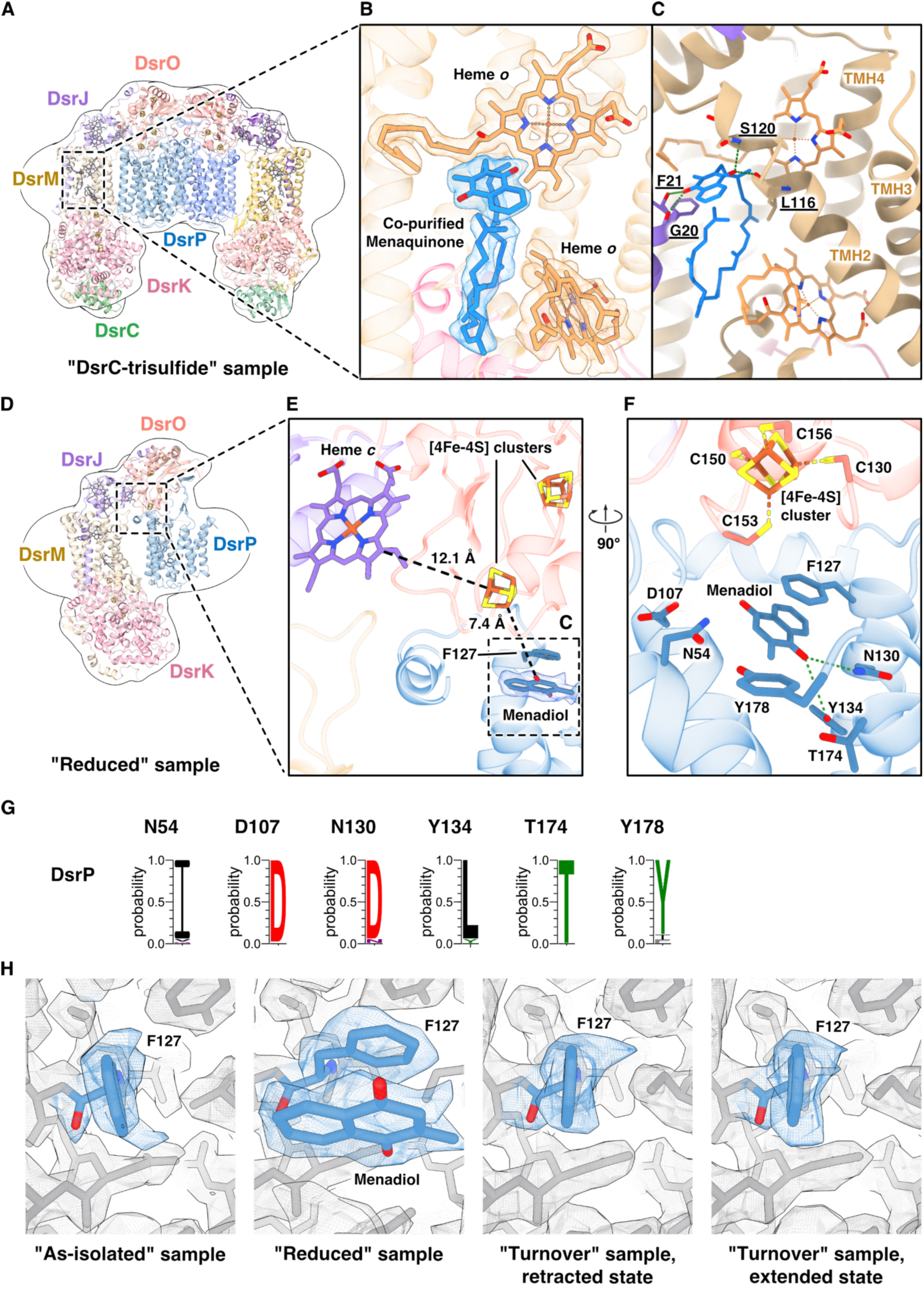
Menaquinone and menadiol binding by the membrane-bound DsrM and DsrP subunits. **(A-C)** MK-7 co-purifies with DsrMKJOP, bound in a cleft formed by DsrM and DsrJ. In the “DsrC-trisulfide” sample, where the MK-7 density is best resolved, the head group of MK-7 establishes four hydrogen bonds (green dashed lines in (C)), while its polyisoprenyl tail is surrounded by hydrophobic residues (fig. S11B). Residues involved in hydrogen bonding via their main-chain atoms are underlined (C). **(D-E)** Menadiol-incubated DsrMKJOP shows menadiol binding at DsrP, positioned to allow efficient electron transfer to one [4Fe-4S] cluster of DsrO, as indicated by edge-to-edge distances (black dashed lines) between cofactors in (E). **(F)** Phe127^DsrP^ and nearby residues likely involved in hydrogen bonding (green dashed lines) to menadiol are shown; most are conserved **(G)**. **(H)** Phe127^DsrP^ undergoes a conformational change to accommodate menadiol binding. After mixing the menadiol-incubated sample with DsrC-trisulfide, menadiol is absent and Phe127 reverts to the conformation seen in the as-isolated sample. Cryo-EM maps in (A, D) were Gaussian-filtered to 4Å and contoured at 5.7 and 8.1 σ, respectively, outlining the micelle. MK-7 and heme *o* densities in (B) are contoured at 5.96 σ; menadiol (E) at 11.71 σ. In (H), densities are contoured at 1.97, 15.4, 4.9, and 4.9 σ, respectively.

This comparison provides important insight into differences between DsrM proteins that are found as part of a DsrMKJOP complex (here termed DsrM-2) and DsrM proteins that are part of more ancient DsrMK complexes, lacking subunits J, O, and P (here termed DsrM-1) (see Supplementary Text). Sequence alignments comparing DsrM-1 and DsrM-2 show that within SRM, the features occluding the quinol oxidation site (arginine residues and loop insertion) are conserved only in DsrM-2, whereas DsrM-1 proteins share a similar sequence pattern with NarI (fig. S15). AlphaFold 3 (AF3) structural models support the sequence-based prediction that the quinol oxidation site is accessible in DsrMK but occluded in DsrMKJOP (figs. S16-17). It seems that the transition from DsrMK to DsrMKJOP involved both the blocking of the quinone-binding site and the creation of the DsrM-DsrJ interface, since both appear to rely on the helix-breaking loop in TMH3 (see Supplementary Text). Similar occluded quinol-site architecture is observed for other DsrM homologs, but with different sequence signatures. DsrM proteins from sulfur-oxidizing *Chlorobiota* and cable bacteria closely resemble the reductive-type DsrM-2 (fig. S18). In contrast, sulfur-oxidizing *Pseudomonadota* lack this loop interruption but encode a conserved proline that is predicted to induce a helix break in TMH3 that similarly blocks the quinol pocket (fig. S18). Thus, two separate sequence signatures within the DsrM-2 proteins are predicted to encode slightly different structural variants resulting in the same outcome: quinol-site occlusion. Notably, unlike other DsrM-2 systems, cable bacteria encode a DsrMKJ complex, whereas DsrOP homologues are predicted from genome organization to belong to a separate complex with a tetraheme cytochrome *c* ^28,29^ (named DsrO_h_P_h_Cyt), with unknown physiological function, which is also present in some organisms without dissimilatory sulfur metabolism ^29^. Structure predictions of the DsrMKJ and DsrO_h_P_h_Cyt subcomplexes gave architectures similar to those observed within subcomplexes of our DsrMKJOP assembly, but differences in DsrO, DsrP, and DsrJ would produce substantial clashes if arranged in a DsrMKJO_h_P_h_ structure, which could not be stably predicted. This structural analysis gives strong support to the suggestion that in cable bacteria DsrMKJ and DsrO_h_P_h_Cyt are two separate protein complexes, with distinct physiological functions ^29^ (fig. S19; Supplementary Text).

### DsrP harbors a redox-active quinol-binding site

DsrP belongs to a family of cofactor-free quinone-binding proteins related to the nitrite reductase subunit NrfD ^16^. Based on this homology, DsrP has been suggested to bind quinone/quinol ^14,18^, but our structure of “as-isolated” DsrMKJOP shows no evidence of copurified quinol in DsrP. To try to characterize the quinol-bound, reduced state, we incubated DsrMKJOP with menadiol at 60 °C for one hour, resulting in the reduction of the hemes and non-cubane cluster ^11^ (fig. S20), and plunge-froze the sample for structural analysis without further substrate addition. Cryo-EM data processing revealed dissociation of the decamer into DsrMKJOP pentamers, along with a population of other subcomplexes (fig. S5), possibly related to thermal treatment. In pentameric DsrMKJOP, menadiol is clearly resolved within a binding pocket of DsrP near the periplasmic side of the membrane (Fig. 2D-E), but is not observed in DsrM. The menadiol is surrounded by hydrogen-bonding residues (Fig. 2F-G), as well as hydrophobic residues from DsrP and DsrO (fig. S21). A similarly positioned quinol-binding pocket has been observed for two NrfD family members ^30,31^ and proposed in others ^32–35^ (figs. S22-23). The menadiol is located less than 8 Å from the nearest DsrO [4Fe-4S] cluster, a distance compatible with efficient electron transfer (Fig. 2E). Comparison with the “as-isolated” structure shows that menadiol binding is accompanied by a flip of the Phe127^DsrP^ phenyl ring, enabling π-stacking interaction with the ligand (Fig. 2H). Surface analysis of DsrP and DsrO reveals that the menadiol-binding pocket is connected — via the strictly conserved Asp107^DsrP^ — to a solvent-accessible cavity at the subunit interface (fig. S24). In the related ActC subunit of alternative complex III, this position is occupied by a conserved His (fig. S23) suggesting a possible functional conservation. The solvent-accessible cavity is lined with polar residues, among them the conserved Glu359^DsrP^ and Arg154^DsrO^, and likely permits water entry, suggesting a proton pathway toward the periplasm (fig. S24C-E).

In the “turnover” sample, the menadiol-reduced DsrMKJOP was mixed with the DsrC-trisulfide. In this sample, DsrMKJOP decamers were again observed, reversing the dissociation to pentamers observed in the menadiol heat-treated condition. Incubation with DsrC-trisulfide caused the menadiol density in DsrP to disappear, and Phe127^DsrP^ reverted to the position observed in the ‘as-isolated’ sample, where it partially overlaps with the quinone-binding site (Fig. 2H). Therefore, addition of DsrC-trisulfide substrate appears to have triggered menadiol oxidation and release from DsrP, suggesting that electrons were transferred from DsrP to DsrK.

### A unique catalytic site for trisulfide reduction in DsrK

DsrK is closely related to the catalytic subunits of methanogenic heterodisulfide reductases, HdrBC and HdrD ^14,19^ (Fig. 3A), which mediate heterodisulfide bond cleavage at a pair of non-cubane [4Fe-4S] clusters ^20–22,36^. In contrast, DsrK harbors a single non-cubane cluster at the position equivalent to HdrB’s proximal cluster, HB1 (Fig. 3A-B). One ligand of HB1, the “supernumerary” Cys234, is replaced in DsrK by Asp500 (Fig. 3B). All residues coordinating the distal cluster, HB2, within HdrB are replaced in DsrK, except for Cys329^DsrK^, which is strictly conserved among DsrK homologs (Fig. 3C) ^11^, suggesting a likely functional role. Our map shows additional density at Cys329, assigned as a cysteine persulfide (Fig. 3B), and this modification persists in both the menadiol-reduced and “turnover” states. In HdrB, the nearby His154 is proposed to mediate proton transfer during disulfide reduction ^20,37^, but this residue is not conserved in DsrK. Its role may instead be fulfilled by His331 (Fig. 3B), which is well conserved in DsrK-related proteins featuring a single non-cubane cluster (Fig. 3C) ^11^, but replaced by other residues in members of the Hdr family containing two non-cubane clusters ^11^. Altogether, these differences point to a distinct catalytic strategy in DsrK compared to the better-studied HdrBC.

**Fig. 3.**
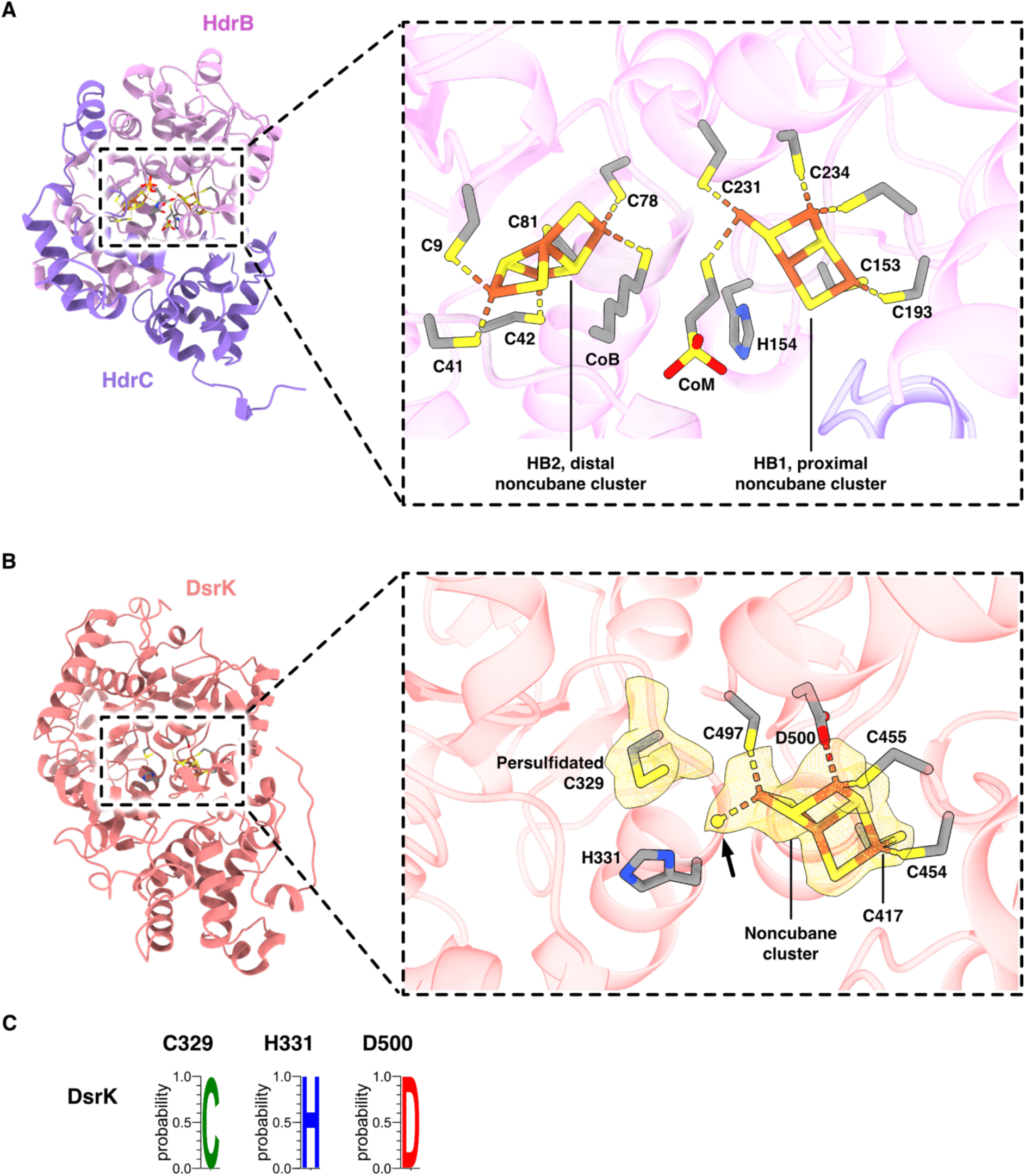
The catalytic subunit DsrK is homologous to the methanogenic HdrBC. **(A)** HdrBC from *Methanothermococcus thermolithotrophicus* (PDB ID 5ODR) ^20^ in cartoon representation, with HdrB colored in pink and HdrC in purple. Inset: a detailed view of non-cubane clusters involved in heterodisulfide reduction. Coenzyme B (CoB), modeled with a shortened tail for clarity, is covalently bound to the distal non-cubane cluster, whereas coenzyme M (CoM) is bound to the proximal one. **(B)** DsrK resolved in the current cryo-EM study (“as-isolated” sample), presented as coral cartoons. Inset: a detailed view of the single non-cubane cluster within DsrK. The cryo-EM densities, contoured at 1.7 σ and displayed as yellow surfaces, include an elongated density near the ‘special’ iron. This density, modeled as a sulfide, is highlighted with an arrow. **(C)** DsrK residues C329, H331, and D500 are strictly conserved among a diverse set of Dsr proteins.

A recent spectroscopic and computational study concluded that HB1 and HB2 in HdrB switch from a closed to a more open state upon substrate binding, during which the ‘special’ substrate-binding iron shifts to a nonbonding distance from the supernumerary cysteine, instead forming a bond with the substrate cysteine thiolate ^37^. To date, no structural data were available to confirm this suggestion. Our substrate-free structure shows the non-cubane cluster already in such an open conformation, with a distance of 4 Å between the Asp500 and the ‘special’ Fe. Consistent with the proposal from this study ^37^, we observe additional density at this iron due to a small ligand (Fig. 3B). Although the ligand’s identity is unknown, we tentatively assign it as a sulfide based on its fit to the density and the high sulfide availability in SRM. The presence of this ligand, together with the open cluster geometry, may explain why the non-cubane cluster of as-isolated DsrK exhibits an electron paramagnetic resonance (EPR) signal resembling that of substrate-bound Hdr ^14,19^. In the menadiol-reduced state, which a previous EPR study showed causes reduction of the non-cubane cluster^11^, we observe a very similar open conformation of the cluster, and persistence of the ligand at the ‘special Fe’.

### DsrK activates DsrC-trisulfide, allowing its C-terminus to approach the non-cubane cluster

Unlike HdrBC and HdrD, which act on small-molecule heterodisulfides, DsrK targets a DsrC-bound S^0^ atom carried as a trisulfide bridge ^11^. How this bulky protein-based substrate reaches the non-cubane cluster — located deep within a cavity of DsrK — is unknown. Combined with the trisulfide’s exceptional resistance to reductants, this raises key questions about the mechanism of trisulfide reduction by DsrMKJOP. To investigate this, DsrC-trisulfide was prepared as previously described ^10^ and verified by intact mass spectrometry (MS) and gel-shift assay (figs. S25-26). After preparation of the “reduced” DsrMKJOP sample by incubation with menadiol at 60 °C for one hour, we added DsrC-trisulfide at the same temperature, mixed and plunge-froze on EM grids within 30 seconds (“turnover” sample, Fig. 4A; see methods). From this sample, supervised classification yielded two structural states of DsrC bound to DsrMKJOP via the DsrK subunit (figs. S6, S27), with the interface mainly stabilized by electrostatic and hydrophobic interactions (fig. S28).

**Fig. 4.**
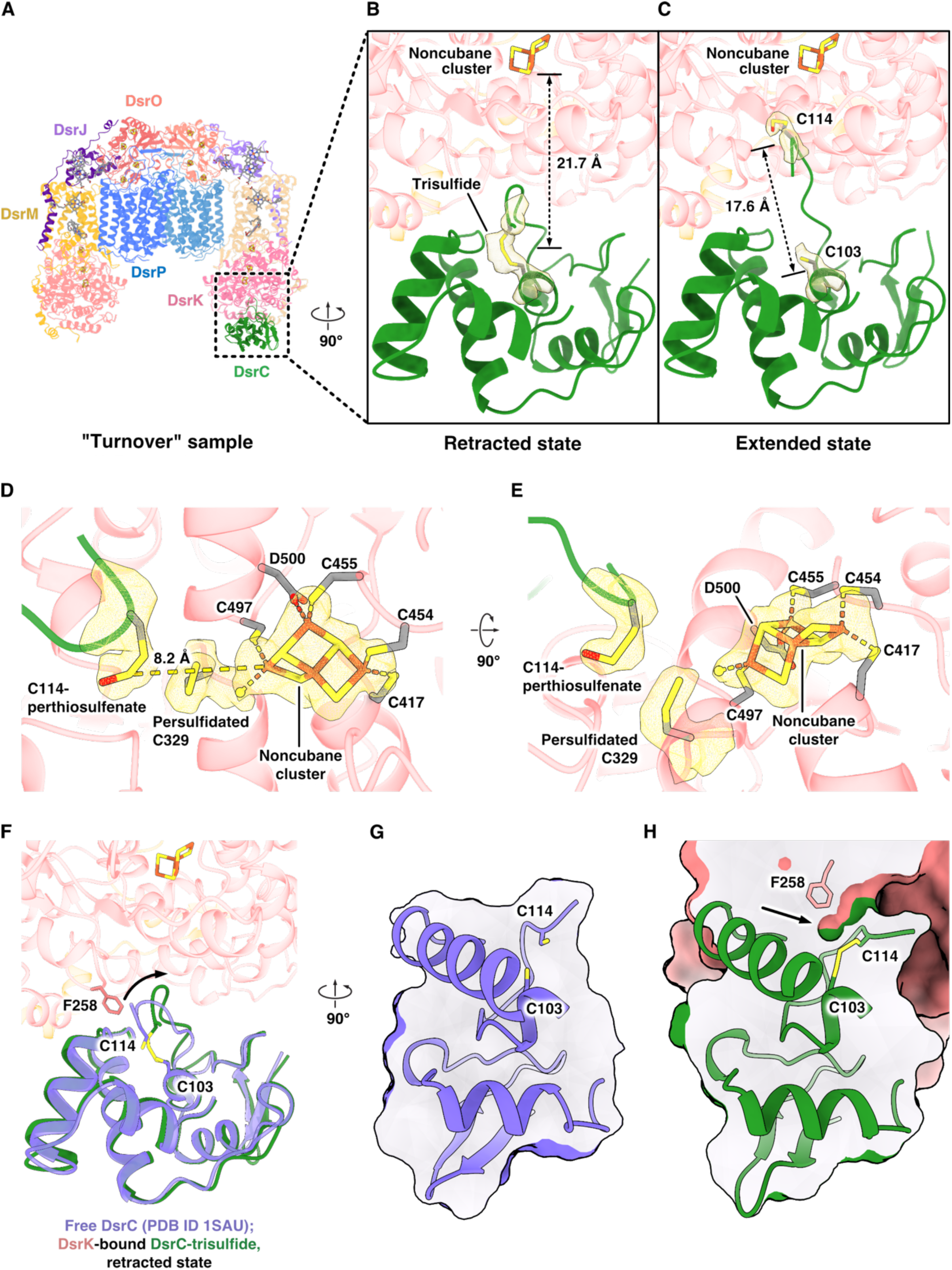
Hydrolysis of DsrC-trisulfide is the first step towards its reduction. **(A)** Two states were observed following the co-incubation of DsrMKJOP with menadiol and DsrC-trisulfide. The retracted state features an intact trisulfide **(B)**, while the extended state contains a hydrolyzed trisulfide **(C)**. The cryo-EM densities of the trisulfide, as well as Cys103^DsrC^ and Cys114^DsrC^, are shown as yellow meshes contoured at 7 σ. **(D-E)** Cys114^DsrC^ in the extended state, modified as a perthiosulfenate, is positioned over 8 Å from the non-cubane cluster within DsrK. The cryo-EM densities of the non-cubane cluster, along with Cys329^DsrK^ and Cys114^DsrC^, are shown as yellow meshes contoured at 6.28 σ. **(F-H)** Phe258^DsrK^ appears to trigger a local rearrangement of the DsrC C-terminal arm (curved arrow), forming a narrow crevice (straight arrow) that increases solvent exposure to promote trisulfide hydrolysis. Free DsrC (purple, PDB ID 1SAU) ^38^ and DsrC in complex with DsrMKJOP resolved by cryo-EM (green) are aligned according to DsrC. To visualize the crevice, solvent-excluded surfaces are generated by ChimeraX ^47^ in (G) and (H), and colored consistently with the left panel.

In one state, resolved to 2.6 Å (table S2), the C-terminal arm of DsrC is retracted, positioning the two strictly conserved C-terminal cysteines, Cys103 and Cys114 ^10^, close together (retracted state, Fig. 4B). Unlike existing structures of DsrC [*e.g.* Protein Data Bank (PDB) ID 1SAU] ^38^, our cryo-EM map reveals continuous density between Cys103 and Cys114 that fits well to the cysteine-bound trisulfide (Fig. 4B and fig. S27C), with local resolution in this region ranging from 2.7 to 2.9 Å. In this state, the trisulfide is 22 Å from the non-cubane cluster (Fig. 4B), the closest redox cofactor. This distance is too long to support rapid electron transfer for direct trisulfide reduction. In a second state, resolved to 2.5 Å (table S2), the trisulfide linkage is cleaved and the C-terminal arm extends toward the catalytic site of DsrK, separating Cys114 and Cys103 by over 17 Å (extended state, Fig. 4C). Because of the long distance to the catalytic site in the closed state, this conformational change must result from electroneutral opening of the trisulfide, which we propose as the initial activation step in DsrC-trisulfide reaction with DsrK. A plausible mechanism for this activation is hydrolysis, reversing the final step of DsrC-trisulfide formation by DsrAB, which involves condensation of a Cys114 perthiosulfenate (Cys-S–S–OH) with a Cys103 thiol ^10^. Supporting this model, our cryo-EM map of the extended state shows density at Cys114 that is consistent with a perthiosulfenate (Fig. 4C-E, fig. S27C), stabilized by hydrogen bonding with His210^DsrK^ and Glu292^DsrK^ (fig. S29).

The model we propose is well supported by previous findings: mutation of Cys114^DsrC^ abolishes growth entirely, whereas Cys103^DsrC^ mutation only slows growth ^10^. Reaction of DsrAB with a DsrC Cys103Ala variant yields a product whose mass is consistent with a Cys114-perthiosulfenate intermediate. Unlike DsrC-trisulfide, this intermediate is unstable ^10^. The ability of the Cys103Ala variant to support growth indicates that the Cys114-perthiosulfenate intermediate can still serve as a substrate for reduction by DsrMKJOP. Nevertheless, Cys103^DsrC^ is strictly required to form the trisulfide product, whose enhanced stability likely confers a selective advantage, explaining both the conservation of Cys103^DsrC^ and the slower growth of the Cys103Ala mutant.

Despite the substantial conformational shift, the DsrC Cys114-perthiosulfenate remains over 8 Å from the non-cubane cluster (Fig. 4D-E). The strictly conserved Cys329^DsrK^ lies between them (Fig. 4D-E), and the likely persulfidation observed here suggests it may serve as a sulfur relay, accepting sulfur from Cys114^DsrC^ and transferring it to the cluster.

We performed intact MS to monitor the mass shifts in DsrC-trisulfide upon interaction with DsrMKJOP. Reaction of reduced DsrC with DsrAB produced a +30 Da mass shift corresponding to the formation of DsrC-trisulfide (fig. S25), which remained stable in isolation over a period of four weeks at 4°C under anaerobic conditions (fig. S30), in agreement with previous studies ^11^. Upon incubation with DsrMKJOP, we observed re-formation of reduced DsrC (figs. S31-32). Additional peaks corresponding to the incorporation of one or two sulfur atoms beyond trisulfide also appeared, particularly in experiments using stoichiometric amounts of DsrMKJOP and DsrC (fig. S32). This indicates that DsrMKJOP can transfer additional sulfur atoms to DsrC, consistent with the extra sulfurs observed at the active site, which might be related to reported production of zero-valent sulfur by SRM ^39^. In some experiments, we also detected a minor peak with a mass shift (+16 Da) matching a perthiosulfenate species (fig. S32), which fits well with our proposed hydrolytic activation mechanism. This peak is not always observed, which could be explained by the transient nature of this intermediate, but further experiments are required to confirm its identity and role.

Our structural and MS evidence, together with previous mutagenesis data ^10^, support a mechanism in which DsrC-trisulfide is first hydrolyzed, allowing the DsrC C-terminal arm to approach the DsrK catalytic site. In the crystal structure of reduced DsrC ^38^, which also adopts a retracted conformation, the two conserved cysteines are sheltered from solvent within a highly hydrophobic environment (Fig. 4F-G, fig. S33). This is likely important not only to reduce the rate of hydrolysis, but also in rendering the direct reduction of the trisulfide in free DsrC unfavorable, since reduction would generate charged products in a strongly hydrophobic environment, which could explain the stability of DsrC-trisulfide to direct reduction. In comparison, our structure of DsrC trisulfide bound to DsrK in the retracted state shows increased solvent accessibility around these cysteine residues, due to insertion of conserved Phe258^DsrK^ (fig. S28E) beneath the C-terminal arm of DsrC, lifting it and exposing the underlying trisulfide (Fig. 4H). Thus, it would appear that the major effect of DsrC binding to DsrK is to increase local water access to the trisulfide through the insertion of Phe258^DsrK^, which likely explains the essential role of DsrK in promoting trisulfide hydrolysis, in contrast to the remarkable stability of DsrC-trisulfide in the isolated form ^11^. We asked whether any nearby residues are well-placed to act as acidic or basic catalysts to promote hydration of the trisulfide, but found no obvious candidates. In this context, it is worth considering that hydrolysis of a trisulfide is expected to occur more easily than the equivalent hydrolysis of a disulfide, given the relatively electrophilic character of the bridging sulfane sulfur.

## Discussion

Our data provide new insights into the mechanism of DsrMK(JOP)-catalyzed trisulfide reduction. We show that the redox cofactors within this large complex form a pathway for rapid electron transfer across the entire complex. While both membrane-integral subunits were predicted from homology to contain a quinol-oxidizing site, our structures suggest that the reduction of DsrC-trisulfide is coupled to menaquinol oxidation in DsrP. In contrast, DsrM binds a copurified MK-7 molecule that apparently plays a structural role, while a separate quinone-binding site identified in the homolog NarI is blocked in DsrM. This was unexpected, because some SRM express only a dimeric DsrMK complex, which is predicted to be the more ancient form of the complex (fig. S34) ^15^, with homology to methanogenic HdrED ^22–24^. Sequence comparison and structure prediction suggest an important difference between DsrM proteins present in reductive DsrMK, *i.e.*, DsrM-1, and those contained in later-emerged DsrMKJOP complexes, *i.e.*, DsrM-2. Similar to the homolog NarI, DsrM-1 appears to be able to bind and oxidize quinol at a site near the periplasmic face of the membrane, while the equivalent position is blocked for quinol binding in DsrM-2. Therefore, we hypothesize that, in DsrMK complexes, quinol is oxidized at DsrM, while in DsrMKJOP complexes, quinol is oxidized at DsrP, with sequential electron transfer through DsrO, DsrJ, DsrM, and finally DsrK. This is consistent with our mutagenesis results showing that deletion of *dsrJ* or *dsrJOP* abrogates growth by sulfite respiration (fig. S10). An exception is within the cable bacteria, where the DsrMKJ complex seems to have a blocked quinone-binding site, so an additional partner for electron exchange may be involved.

Our structures indicate that protons from menaquinol oxidation at DsrP are released to the periplasmic side of the membrane, while protons for DsrC reduction come from the cytoplasm, allowing the overall reaction to contribute to pmf generation. Early studies analysing the growth yield of *Nitratidesulfovibrio (Desulfovibrio) vulgaris* during growth on hydrogen (H_2_) with sulfite or thiosulfate concluded that approximately 3 – 3.5 moles of ATP are produced per mole of sulfite reduced to sulfide ^40,41^. Assuming an ATP synthase c-ring stoichiometry of around 10 (similar to other non-alkaliphilic bacteria), sulfite reduction to sulfide should move a minimum of 10 - 12 charges across the membrane. This reaction requires six electrons, four of which are supplied through the DsrMKJOP complex and two from an unknown electron donor to DsrAB (fig. S35). In *N. vulgaris*, assuming DsrAB reduction proceeds with electrons derived from a periplasmic hydrogenase, as for DsrMKJOP, the overall H_2_/sulfite redox reaction releases six protons into the periplasm and consumes six protons in the cytoplasm. In addition, the redox loop formed by quinone reductase complex (QrcABCD) and DsrMKJOP, involving proton uptake by QrcD ^42^ and release by DsrP, contributes four further protons to the pmf. Thus, the overall process could be associated with translocation of 10 H⁺ across the membrane, perhaps just sufficient to produce 3 ATP, under the stated assumptions (fig. S35). Nevertheless, the overall pmf generation for this reaction would be expected to be the same as when quinol is oxidized in DsrM of DsrMK complexes, given the periplasmic location of that binding site. What advantage do organisms gain by expressing a full DsrMKJOP complex? One obvious explanation would be that the DsrP may couple quinol oxidation with vectorial transport of one or more protons across the membrane, as previously suggested for the homologous subunits of polysulfide reductase ^30^, alternative complex III ^31,32,34,35^, and the quinone-independent reductive dehalogenase complex ^43^. However, a clear structural basis for vectorial proton transport or its coupling to quinol oxidation is not obvious from our data, so this needs to be tested experimentally. A previously suggested alternative was that the rare Cys/His-coordinated heme *c* of DsrJ could catalyse additional reactions with a periplasmic sulfur substrate ^16,44^. However, we find that mutation of the heme-coordinating cysteine does not impair the physiological function of DsrMKJOP in sulfate- or sulfite-dependent growth. Taken together with previous findings that neither thiosulfate nor sulfite can serve as an additional electron donor to DsrMKJOP ^11^, this hypothesis seems unlikely.

The established catalytic model for non-cubane clusters derives from the methanogenic Hdr enzymes, in which a pair of non-cubane clusters act in concert to cleave a disulfide bond, generating sulfur adducts at each cluster that are released upon cluster reduction. While this mechanism suits the Hdr system, it does not readily explain how a single non-cubane cluster, bound deeply within a narrow cavity, could mediate the four-electron reduction of DsrC-trisulfide to regenerate DsrC with two free cysteine thiols. Our results provide a new paradigm for understanding respiratory sulfide production in SRM (Fig. 5). The highly stable DsrC-trisulfide is activated only upon binding to DsrK, which increases the solvent accessibility of the trisulfide bond and promotes its hydrolytic activation, yielding a reduced Cys103 and a perthiosulfenate at Cys114. This hypothesis agrees well with our MS data (figs. S30, S32) and previous mutagenesis results ^10^. Hydrolysis of the trisulfide linkage allows the C-terminal arm of DsrC to swing into the “extended” conformation and insert into the DsrK catalytic site where the conserved Cys329 promotes transfer of the sulfenate moiety and release of reduced DsrC, with proton uptake from the cytoplasm. The perthiosulfenate group now at Cys329 is finally reduced at the non-cubane cluster with release of H_2_S. This mechanism helps to explain how the DsrC-trisulfide, which is surprisingly stable to reduction in isolation, can be efficiently reduced in the physiological context. A corollary of this is that the non-cubane [4Fe-4S] cluster of DsrK can transform oxygenated sulfur species, rather than simply breaking disulfide bonds as in Hdr. Similar reactivity has previously been proposed for homologous complexes (*e.g.,* sulfur-oxidizing heterodisulfide reductase-like complex (sHdr) from sulfur-oxidizing bacteria (SOB)) ^45^ but so far direct evidence for such reactivity has been lacking. The replacement of a cluster-coordinating cysteine in Hdr with a carboxylate ligand (Asp500) in DsrK may tune the properties of the cluster, for example, through the hard-soft acid-base ‘symbiotic’ effect ^46^, making it more suited to react with a harder substrate such as perthiosulfenate. The detailed mechanism of the four-electron perthiosulfenate reduction by the non-cubane cluster will require further investigation.

**Fig. 5.**
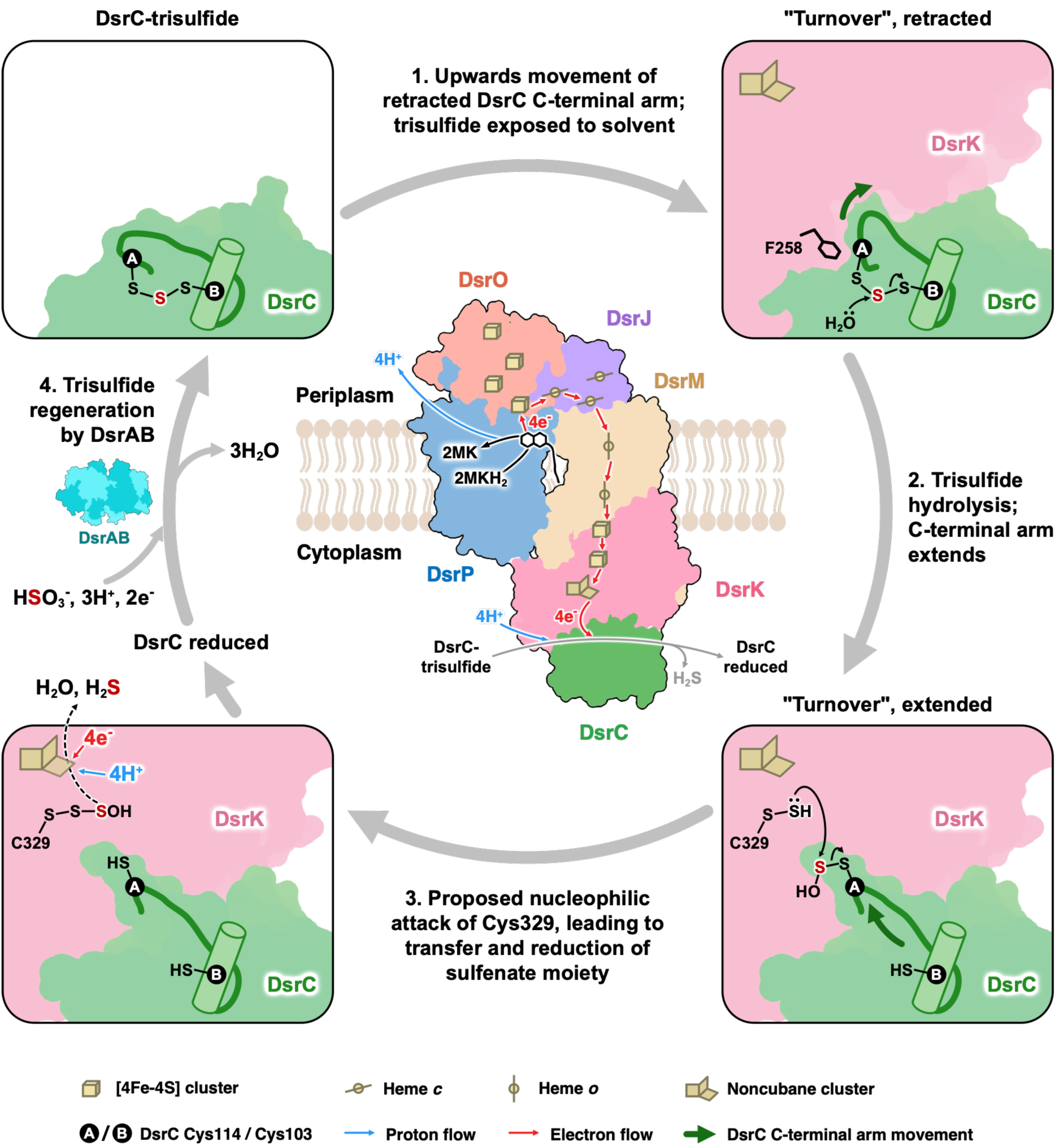
Schematic of DsrC-trisulfide reduction by DsrMKJOP. A proposed model for DsrC-trisulfide reduction, driven by the oxidation of DsrP-bound menaquinol. The overall reaction, which generates pmf through electron transfer (red arrows) to cytoplasmic DsrC and proton release (light blue arrow) into the periplasm, is illustrated in the central panel. A flat surface representation of the DsrMKJOP pentamer, based on the “turnover” sample in the retracted state, is shown. Local conformational (dark green arrows) and chemical (solid black arrows) transformations essential for DsrC-trisulfide reduction are shown in the corner panels and described alongside the gray curved arrows. Reduction of the sulfenate moiety on Cys329^DsrK^ through the non-cubane cluster is marked by a dashed black arrow in the lower left panel. The S^0^ in the trisulfide and its derivatives throughout the cycle are colored dark red. *A. fulgidus* DsrAB (PDB ID 3MMC) ^48^ is shown in flat surface representation, with DsrA colored dark turquoise and DsrB colored cyan.

Our structural and biochemical data allowed us to propose a mechanism of the reactivity in the widely-conserved complex DsrMKJOP (movie S1). This work gives new insight into how biological systems control the reactivity of sulfur and informs our understanding of sequence-structure-function relationships across a wide diversity of sulfur-metabolizing organisms.

## Methods

### Construction of *N. vulgaris* deletion strains

*Nitratidesulfovibrio vulgaris* Hildenborough (*N. vulgaris*), a genetically tractable SRM formerly known as *Desulfovibrio vulgaris* Hildenborough, was used to study the physiological role of the DsrJOP module and the DsrJ subunit. *N. vulgaris* strains lacking the *dsrJOP* (Δ*dsrJOP*) or the *dsrJ* gene (Δ*dsrJ*) were constructed by double homologous recombination. Suicide plasmids for insertion of a kanamycin resistance cassette in both strains were constructed using the Sequence- and Ligation-Independent Cloning (SLIC) method, as described before ^49^. *N. vulgaris* genomic DNA was used as a template to amplify the upstream and downstream regions of *dsrJOP* or *dsrJ* genes. The kanamycin resistance cassette was amplified from pSC27, while the pUC backbone was amplified from plasmid pMO719. One Shot^TM^ TOP10 chemically competent *Escherichia coli* cells (Invitrogen) were transformed with the assembled plasmids and plated on Luria Bertani (LB) agar plates containing 50 μg/mL kanamycin and 100 μg/mL spectinomycin.

After sequence confirmation, *N. vulgaris* cells were electroporated with the respective suicide plasmids (pMOIPAB01 or pMOIPAB02), using the following conditions: 1500 V, 250 W and 25 µF. Electroporated cells were left to recover anaerobically for 36 h at 37 °C in 1 mL Missouri (MO) medium with 1 g/L yeast extract (MOY), 60 mM sodium pyruvate, and 3 mM sodium sulfate ^50^. After the recovery period, cells were plated inside the anaerobic glove box with the same medium supplemented with 1.5% (w/v) agar, and 400 µg/mL geneticin (G418), an analogue of kanamycin that is more effective in *N. vulgaris*, as described in Keller *et al.* ^50^. The plates were incubated at 37 °C under anaerobic conditions for one week until black ellipsoid *N. vulgaris* colonies appeared. Colonies resistant to G418 but sensitive to spectinomycin were selected and grown in medium containing G418. The absence of the *dsrJOP* operon and the *dsrJ* gene in *N. vulgaris* Δ*dsrJOP* and *N. vulgaris* Δ*dsrJ* strains, respectively, was verified by genomic DNA isolation followed by PCR.

### Construction of *N. vulgaris* complemented strains

To create the pMO-*dsrJ* complemented plasmid, *dsrJ* was amplified from *N. vulgaris* genomic DNA. Following the SLIC protocol described above, this gene was cloned into pMOIP03, a plasmid containing a Strep-tag at the C-terminal of the protein. One Shot^TM^ TOP10 chemically competent *E. coli* cells (Invitrogen) were transformed with this plasmid and plated on LB agar plates containing 100 μg/mL spectinomycin. The correct plasmid construct was screened by colony PCR and later confirmed by sequencing.

Site-directed mutagenesis on the pMO-*dsrJ* complemented plasmid was performed using NZYMutagenesis kit (NZYTech) to replace a conserved cysteine residue (Cys45) with an alanine (C45A), histidine (C45H) or serine (C45S). *E. coli* NZYStar competent cells were transformed with the PCR products treated with Dpn I. After transformation, cells were plated on LB agar plates containing 100 µg/mL spectinomycin. The cloning vectors were verified by sequencing.

*N. vulgaris* Δ*dsrJ* deletion strain was complemented with pMO-*dsrJ*, pMO-*dsrJ* C45A, pMO-*dsrJ* C45H or pMO-*dsrJ* C45S by electroporation, using the following conditions: 1250 V, 250 W and 25 µF. Colonies resistant to both antibiotics (G418 and spectinomycin) were selected as described in the previous section. Complementation with *dsrJ* gene and its point mutations were confirmed by sequencing.

Descriptive lists of plasmids, strains, and primers used throughout this study are presented in tables S4-5.

### Growth studies

*N. vulgaris* WT and mutant strains were grown anaerobically at 37 °C in MOY medium (0.2 g/L yeast extract), under respiratory conditions. The medium was adjusted to contain lactate or formate as electron donor, and sulfate or sulfite as electron acceptor. Lactate/Sulfate medium contained 30 mM sodium lactate and 30 mM sodium sulfate; Lactate/Sulfite medium contained 15 mM sodium lactate and 10 mM sodium sulfite; Formate/Sulfate medium contained 50 mM sodium formate, 20 mM sodium acetate and 30 mM sodium sulfate; and Formate/Sulfite medium contained 50 mM sodium formate, 20 mM sodium acetate and 10 mM sodium sulfite. All media were inoculated with 2% (v/v) of fresh precultured cells grown in MOY medium (0.5 g/L yeast extract) containing 60 mM sodium pyruvate and 3 mM sodium sulfate. Antibiotics G418 at 400 µg/mL and spectinomycin at 100 µg/mL were added to the media according to the strain. The growth of the cultures was monitored by determining the optical density at 600 nm (OD_600_) at various time points, and triplicate biological experiments were performed for each condition.

### Protein Purification

DsrMKJOP was natively purified from *Archaeoglobus fulgidus* (*A. fulgidus*) cells. Frozen cells were resuspended in lysis buffer (50 mM potassium phosphate (KPi), pH 7.0, 10% (v/v) glycerol, and cOmplete™ protease inhibitor cocktail (Roche)), in the presence of DNase (Sigma-Aldrich). Cells were lysed in a APV Model 2000 Homogenizer at 80 MPa. Unlysed cells were removed by centrifugation at 7930 × *g* for 20 min at 4 °C. Membrane fraction was prepared by ultracentrifugation at 138000 × *g* for 2 h at 4 °C, followed by overnight solubilization in lysis buffer containing 2% (w/v) n-dodecylβ-D-maltoside (DDM, Glycon Biochemicals GmbH). The solubilized membrane proteins were separated by ultracentrifugation (138000 × *g* for 2 h at 4 °C), and the membrane pellet was used for a second solubilization for 5 h with gentle stirring at 4 °C. A third ultracentrifugation step was performed before loading the membrane fraction onto a Q-Sepharose high-performance column (2.6 × 10.0 cm, GE Healthcare) equilibrated with buffer A (50 mM KPi pH 7.0, 10% (v/v) glycerol, 0.1% (w/v) DDM, and a cOmplete™ protease inhibitor tablet/L). A stepwise gradient of increasing concentrations of NaCl (from 0 to 1 M NaCl) was applied. The fractions eluted at 0.45 M and 0.50 M NaCl were pooled, concentrated, and the ionic strength of the solution was lowered by dilution with buffer A and ultrafiltration (50-kDa cutoff, Amicon, Millipore). These fractions were then applied into a Resource Q column (1.6 × 3.0 cm, GE Healthcare) equilibrated with buffer A. A stepwise gradient of increasing concentrations of NaCl (from 0 to 1 M NaCl) was performed, and the fractions eluted at 0.30 M and 0.35 M NaCl were pooled, concentrated, and the ionic strength lowered by dilution and concentration steps as described above.

All chromatographic steps were performed at 4 °C, aerobically, and monitored by Ultraviolet– visible (UV–Vis) spectroscopy. The purity of DsrMKJOP was assessed by a 10% Tricine-sodium dodecyl sulfate polyacrylamide gel electrophoresis (Tricine-SDS-PAGE) ^51^, followed by Coomassie Blue staining and heme staining (fig. S1) ^52^. Protein concentration was determined using the previously determined absorption coefficient (ε) of 132.4 mM^−1^.cm^−1^ at 555 nm in the reduced state ^11^.

To prepare the DsrC-trisulfide, DsrAB was natively isolated from the soluble fraction of *A. fulgidus*, and DsrC from the same organism was heterologously expressed in *E. coli* and purified by affinity chromatography. DsrC-trisulfide was enzymatically produced *in vitro* by reaction of DTT-reduced DsrC with DsrAB and sulfite. Detailed purification protocols, as well as procedures for DsrC-trisulfide production and analysis, are described in earlier studies ^10^.

### Mass spectrometry (MS) analysis

Intact MS was performed to analyze DsrC in isolation and upon incubation with the DsrMKJOP complex. The stability of the DsrC-trisulfide was assessed by storing the sample under anaerobic conditions at 4 °C for four weeks. Each week, 30 µg of sample was analyzed by MS to monitor the integrity of the trisulfide. For the interaction assays, 10 µM DsrC-trisulfide was incubated with 100 nM DsrMKJOP and 500 µM menadiol in 50 mM KPi buffer (pH 7.0) for 30 min at 60 °C. The assay was performed inside a Coy anaerobic chamber (98% N_2_, 2% H_2_). Stoichiometric amounts of DsrC-trisulfide and DsrMKJOP (10 µM each) were incubated with 0.5 or 5 mM menadiol (see Fig. S32 legend) at 60 °C following the same procedure. For this assay, different time points were tested (1, 10, and 30 min), and the effect of the absence or presence of menadiol was also assessed. At the end of each experiment, samples were immediately flash-frozen in liquid N_2_ and stored at −80 °C until MS analysis.

Samples were thawed and subjected to an ultrafiltration clean-up procedure using 10 kDa cutoff filtration units with water and 0.1% formic acid. The data was acquired in positive TOF-MS mode using an X500B Q-TOF with the twin-spray ion source (Sciex) mass spectrometer coupled to an ExIonLC. Proteins were separated by RP using a Acquity UPLC Protein BEH C4, 300 Å 1.7 µm 2.1×150 mm (from Waters), at 60 °C, flow rate of 0.2 mL/min and using water and 0.1% formic acid (buffer A) and acetonitrile and 0.1% formic acid (buffer B) as mobile phase. The LC gradient is as follows: 0-10 min from 15% to 90% B, 10-11 min at 90% B, 11-12 min from 90% to 15% B, 12-15 min at 15% B. The MS was set to TOF-MS Intact protein mode with an ion accumulation of 1 sec, 40 bins to sum, and the m/z range from 600-3000. The ESI parameters were as follows: ion source gas 1 at 60, ion source gas 2 at 40, curtain gas at 30, ionization source voltage floating at 5500 V, temperature at 500 °C, declustering potential at 110 V, and collision energy at 15 V. Data were acquired and processed using the Sciex OS software 2.3 at the Mass Spectrometry Unit (UniMS), ITQB/iBET, Oeiras, Portugal.

### Methoxy-polyethylene glycol maleimide (MalPEG) gel-shift assays

To analyze the redox state of DsrC cysteines, a MalPEG gel-shift assay was performed. DsrC-trisulfide (10 μg in 25 mM KPi pH 7.0) was incubated with 1 mM MalPEG (MW 5,000 g/mol; Fluka) at 37 °C for 15 minutes under aerobic conditions. The reaction was quenched by adding SDS loading buffer (62.5 mM Tris-HCl pH 6.8, 5% (w/v) SDS, 5% (v/v) glycerol, 0.025% (w/v) bromophenol blue). Samples were analyzed by 10% Tricine-SDS-PAGE without reducing agent and boiling, followed by Coomassie Blue staining. As a control, untreated DsrC-trisulfide (without MalPEG) and reduced DsrC were also analyzed by SDS-PAGE.

### Specimen preparation for cryo-electron microscopy (cryo-EM)

DsrMKJOP and DsrC were exchanged into a new buffer (20 mM HEPES pH 7.0, 20 mM NaCl, and 0.017% (w/v) DDM), and cryo-EM specimens were prepared under four different biochemical conditions, *i.e.*, as-isolated DsrMKJOP (“as-isolated” sample), DsrMKJOP thoroughly incubated with DsrC-trisulfide (“DsrC-trisulfide” sample), DsrMKJOP incubated with menadiol (“reduced” sample), DsrMKJOP incubated with menadiol and DsrC-trisulfide (“turnover” sample). For all four conditions, C-Flat holey carbon grids (1.2/1.3, 400 mesh; Protochips) were glow-discharged using a PELCO easiGlow system (90 s, 15 mA, 0.38 mBar air) immediately prior to use. For “as-isolated” and “DsrC-trisulfide” samples, 1.5 mM fluorinated fos-choline-8 (0.5x critical micelle concentration; Anatrace Products LLC) was added to each sample immediately before grid application. A 3 μL aliquot of each condition was applied to an individual grid and blotted for 6-10 seconds with filter paper (595; GE HealthCare), and plunge-frozen in liquid ethane using a Vitrobot Mark IV (Thermo Fisher Scientific) at 4 °C and 90% humidity. Additional differences for each sample preparation are described below.

<u>“As-isolated” sample</u>: specimens were prepared under aerobic conditions. The final concentration of the DsrMKJOP pentamer was 22.9 μM.

<u>“DsrC-trisulfide” sample</u>: specimens were prepared in an anaerobic chamber (Coy Laboratories) containing 3-5% H_2_ in N_2_. Grids were glow-discharged immediately before being transferred into the anaerobic chamber. Final concentrations of the DsrMKJOP pentamer and DsrC-trisulfide were 14 μM and 176 μM, respectively.

<u>“Reduced” sample</u>: specimens were prepared in an anaerobic chamber (Coy Laboratories) containing 3-5% H_2_ in N_2_. DsrMKJOP was mixed with menadiol and incubated at 60 °C for one hour. Insoluble menadiol was then removed by filtration using Costar Spin-X centrifuge tube filters (0.22 μm pore size; Corning). The filtrate was subsequently plunge-frozen, with final concentrations of 11 μM for the DsrMKJOP pentamer and up to 500 μM for menadiol. Samples of 3 μL DsrMKJOP, taken before and after menadiol incubation, were used for UV– Vis spectroscopy measurements (fig. S20).

<u>“Turnover” sample</u>: during the preparation of the “reduced” sample, the filtered mixture of DsrMKJOP and menadiol was further incubated with DsrC-trisulfide at 60 °C, and the mixture was plunge-frozen approximately 30 seconds after the addition of DsrC-trisulfide. The final concentrations were 11 μM for DsrMKJOP pentamer, up to 500 μM for menadiol, and 197 μM for DsrC-trisulfide.

### Cryo-EM imaging

Specimens were imaged by cryogenic transmission electron microscopy, using automated fast acquisition implemented via aberration-free image shift (AFIS) within EPU (Thermo Fisher Scientific) (table S1). Additional differences in the imaging process for each sample are described below.

<u>“As-isolated” sample</u>: data were collected on a Krios G2 operated at 300 kV, equipped with a BioQuantum imaging filter (Gatan) and K3 direct electron detector (Gatan), at a calibrated pixel size of 0.831 Å. Two datasets totaling 14,774 movies were recorded in TIFF format, with a total electron dose of 60 e^-^/Å^2^ distributed over 61 fractions and a defocus range of −0.8 μm to −2.5 μm.

<u>“DsrC-trisulfide”, “reduced”, and “turnover” samples</u>: specimens were imaged using a Krios G4 operated at 300 kV and equipped with a cold field emission gun and a Selectris X imaging filter. For “DsrC-trisulfide” sample, data were collected using a Falcon 4 detector, yielding 13,250 movies in EER format at a pixel size of 0.73 Å. “Reduced” sample was acquired with a Falcon 4 detector and resulted in 9,148 movies, also in EER format. “Turnover” sample used a Falcon 4i detector under otherwise identical settings, generating 16,205 EER-format movies. In all three cases, the total electron dose was 60 e⁻/Å², and the defocus ranged from −0.8 to −2.5 μm.

### Cryo-EM data processing

The critical steps of cryo-EM data processing are illustrated in figures S3–S6. Initially, beam-induced motion correction and dose weighting were performed, using a wrapper for the original MotionCor2 implementation ^53^ for the “as-isolated” sample, and RELION’s own implementation of the MotionCor2 algorithm ^54^ for the other three samples. Subsequently, contrast transfer function (CTF) parameters were estimated from the motion-corrected micrographs using CTFFIND 4.1.13 ^55^.

<u>“As-isolated” sample</u>: two datasets were collected and processed separately up to the completion of Bayesian polishing ^56^. Cryo-EM data processing was carried out using RELION 3.1 ^57,58^ and cryoSPARC 3.3 ^59^. Initial particle picking was performed using the blob picker, followed by 2D classification, both within CryoSPARC live. Coordinates from selected particle classes were then exported via UCSF pyem ^60^ and imported into RELION for further processing, including 2D classification, *de novo* 3D initial model generation, and 3D classification. Selected particle classes were used to train convolutional neural networks in TOPAZ ^61^, which was then used for particle picking. TOPAZ-picked particles were extracted in RELION at an initial pixel size of 1.7529 Å and subjected to successive rounds of 2D and 3D classification to remove junk particles. Particles within well-resolved classes were re-extracted at a pixel size of 1.2465 Å and used for 3D auto-refinement with *C_1_* symmetry. Focused 3D classifications were then performed to isolate particles contributing to the complete DsrMKJOP decamer. These selected particles were refined in two successive 3D auto-refinement steps, first with *C_1_* symmetry and then with *C_2_* symmetry. Subsequently, non-uniform refinement with *C_2_* symmetry was conducted in cryoSPARC, incorporating per-particle defocus refinement and per-exposure-group CTF refinement on the fly. The resulting particles were polished and re-extracted in RELION at a final pixel size of 1.00165 Å. Particles from both datasets were then combined in RELION and subjected to another round of non-uniform refinement with *C_2_* symmetry, followed by heterogeneous refinement to further improve resolution. A final non-uniform refinement with *C_2_* symmetry yielded a reconstruction at 2.2-Å resolution (table S2). The final map was further improved through density modification using *PHENIX* ^62^.

<u>“DsrC-trisulfide” sample</u>: cryo-EM data processing was carried out using RELION 4.0 ^54^ and cryoSPARC 3.3. The dataset was processed following a workflow similar to that used for the “as-isolated” sample, up to the stage of particle picking with TOPAZ. At this point, particles were extracted at a pixel size of 1.5398 Å, and then imported into cryoSPARC for further processing, including 2D classification, *ab-initio* reconstruction, and heterogeneous refinement. A subset of 65,708 particles corresponding to the decameric DsrMKJOP complex was selected and extracted in RELION at a pixel size of 0.9476 Å. Non-uniform refinement was subsequently performed in cryoSPARC with *C_2_* symmetry applied, followed by Bayesian polishing in RELION, during which particles were re-extracted at a pixel size of 0.73 Å. A final round of non-uniform refinement with *C_2_* symmetry yielded a map at 1.97-Å resolution (table S2). The final map was locally filtered in cryoSPARC according to the estimated local resolution.

<u>“Reduced” sample</u>: data processing was carried out using RELION 4.0 and cryoSPARC 4.3. Initial particle picking was performed using the blob picker in cryoSPARC live, followed by 2D classification, *ab-initio* reconstruction, and heterogeneous refinement within cryoSPARC. Well-resolved particle classes were identified, and coordinates from these selected classes were exported to RELION for particle extraction. The extracted particles were then used as training data for TOPAZ. Particles subsequently identified by TOPAZ were extracted in RELION at a pixel size of 1.1406 Å and imported into cryoSPARC for downstream analysis, including 2D classification, *ab-initio* reconstruction, and heterogeneous refinement. A subset of particles corresponding to pentameric DsrMKJOP complex was selected and subjected to non-uniform refinement with *C_1_* symmetry applied, yielding a reconstruction at 2.78-Å resolution. These refined particles were then polished in RELION, re-extracted at a pixel size of 0.9955 Å, and used in a subsequent round of non-uniform refinement in cryoSPARC with *C_1_* symmetry, resulting in a consensus map at 2.29-Å resolution. To further investigate structural heterogeneity, 3D variability analysis (3DVA) was performed and visualized in cluster mode, revealing two particle sub-populations. One sub-population, comprising 65,514 particles, yielded a reconstruction at 2.10-Å resolution following non-uniform refinement (table S2). The final map was locally filtered in cryoSPARC according to the estimated local resolution.

<u>“Turnover” sample</u>: data processing was performed using RELION 4.0 and cryoSPARC 4.3. The dataset was processed following a workflow similar to that used for the “reduced” sample, up to the stage of particle picking with TOPAZ. At this point, particles were extracted at a pixel size of 1.6425 Å and imported into cryoSPARC for further processing, including 2D classification, *ab-initio* reconstruction, and heterogeneous refinement. Subsequently, masked 3D classification was carried out in RELION to select particles contributing to the best-resolved decameric DsrMKJOP complex, yielding a subset of 58,508 particles. These particles were subjected to homogeneous refinement with *C_2_* symmetry applied, followed by re-extraction at a pixel size of 0.8959 Å. Non-uniform refinement with *C_2_* symmetry, Bayesian polishing, and a subsequent round of non-uniform refinement produced a map at 1.98-Å resolution. To further investigate conformational heterogeneity in the vicinity of DsrC, particles were symmetry-expanded (*C_2_*) and subjected to particle subtraction in cryoSPARC to remove regions outside of DsrK and DsrC. Local refinement was then performed, focusing on the DsrK-DsrC region. Inspection of the resulting density revealed conformational flexibility in the region corresponding to the C-terminal arm of DsrC. Based on this observation, models of DsrK and DsrC were built, with the DsrC C-terminal arm in two alternative conformations (fig. S27B). These models were used to generate molmaps in ChimeraX at a resolution of 6 Å. To account for the likely partial occupancy of DsrC, an additional molmap was generated using DsrK alone, also at 6-Å resolution. These three molmaps, lowpass-filtered to 6 Å, served as reference volumes for focused 3D classification in cryoSPARC. During 3D classification, each reference volume was first masked around DsrC using a focus mask (fig. S27C), with external voxels replaced by those from the consensus reconstruction. This was followed by masking around both DsrK and DsrC using a solvent mask (fig. S27C), with voxels outside this region set to zero. One 3D class (43,375 particles) yielded map density for DsrC in the extended state, while another class (40,856 particles) revealed the retracted conformation (fig. S6). Both classes underwent local refinement using the same pre-classification reference. The refined particle subsets were then reverted to the original image stacks and subjected to homogeneous reconstruction in cryoSPARC to generate the respective focused maps (figs. S6 and S27C). These subsets were also locally refined using a mask encompassing the entire decameric complex, resulting in two consensus maps of DsrMKJOP bound to DsrC in extended and retracted states, respectively. For each conformational state, a composite map was generated using phenix.combine_focused_maps ^63^, by integrating the corresponding focused map with the consensus reconstruction from each 3D class.

### Cryo-EM model building and refinement

Initial models were generated by AlphaFold2 ^64,65^ and rigid-body fitted into the corresponding cryo-EM maps. Subsequent manual model tuning and refinement were carried out in Coot ^66^. Cofactors were manually inserted into each model using Coot, while ligan restraints were generated using eLBOW and ReadySet within *PHENIX* ^63^. Real-space refinement was carried out iteratively in *PHENIX* ^67^, alternating with manual adjustments in Coot. During real-space refinement, the strategies minimization_global and local_grid_search were applied. For the models from the “as-isolated” and “DsrC-trisulfide” samples, non-crystallographic symmetry constraints were applied. In the final refinement round, b-factors for the complete models were refined using Servalcat ^68,69^. Maps and models were visualized in figures and in a movie using ChimeraX ^47^.

### Sequence conservation analysis

We selected a diverse set of Dsr protein sequences from SRM with fully sequenced genomes, representing distinct phylogenetic groups, based on a previous large-scale phylogenetic study ^15^. This yielded 61 DsrC, 56 DsrP, 57 DsrO, 80 DsrM, and 52 DsrK sequences. To confidently assign DsrM sequences to either the minimal DsrMK complex or the larger DsrMKJOP complex, we further restricted our analysis to genomes that encode exclusively one of the two configurations. This resulted in 29 DsrM sequences from *dsrMK* operons, classified as DsrM-1, and 49 DsrM sequences from *dsrMKJOP* operons, classified as DsrM-2, the latter all belonging to the DsrM-2a subclass defined later. For DsrK, only those encoded within the *dsrMKJOP* gene cluster were analyzed.

For sequence analysis within SOB, following previous studies ^15,70^, we selected only fully sequenced genomes that lack the sHdr system, an alternative sulfur-oxidizing pathway, to minimize functional redundancy. This dataset comprises two distinct evolutionary lineages of SOB, *Chlorobiota* and *Pseudomonadota*, with 10 and 13 genomes from each phylum, respectively.

For sequence analysis of cable bacteria, we selected all seven closed genomes currently available ^71–75^. We also analyzed 16 genomes of *Gamma-* and *Betaproteobacteria* closely related to cable bacteria, previously reported in a genomic study ^29^, with a focus on DsrO_h_P_h_ and the associated tetraheme cytochrome *c*.

Sequences of each subunit were aligned using Clustal Omega with default parameters ^76^, and the resulting alignments were inspected and analyzed in Jalview (version 2.11.4.1). For structurally important residues, the residue conservation images were generated using the WebLogo 3 web interface (https://weblogo.threeplusone.com) ^77^. To compare sequence patterns between DsrM-1 and DsrM-2a in SRM, their sequences were initially aligned together and then separated for conservation analysis.

### AF3 structure predictions

To investigate structural differences between DsrM-1 and DsrM-2 beyond available experimental structures, predictions were carried out using the AF3 server (https://alphafoldserver.com) ^78^. Each prediction run used one random seed and yielded five structural models. We report the predicted template modeling (pTM) score, a confidence metric provided by AF3 that reflects the accuracy of the overall structure ^79,80^.

As an initial validation step, we assessed whether AF3 could approximate known experimental structures with high accuracy. To this end, we predicted the structures of the *A. fulgidus* DsrMKJOP pentamer and the *E. coli* NarGHI complex, including their associated heme cofactors. The resulting predicted structures were then compared to their respective experimentally determined counterparts. Specifically, the predicted DsrMKJOP models were aligned to our cryo-EM structure of “as-isolated” DsrMKJOP, and the predicted NarGHI models were aligned to an x-ray structure of NarGHI (PDB ID 1Y4Z ^27^) (fig. S16A). Alignments were performed using the Matchmaker tool in ChimeraX, and structural similarity was quantified by calculating the root-mean-square deviation (RMSD) in Å over all residue pairs, as reported by Matchmaker.

Following this validation step, we analyzed SRM to predict structures of DsrM-1 and DsrM-2 proteins that had not previously been structurally characterized; DsrM-2 corresponds to the DsrM-2a subclass defined later. Four phylogenetically distant representatives were selected from organisms encoding either only DsrMK or the full DsrMKJOP complex (figs. S16-17), drawn from the same sequence pool used for sequence conservation analysis. For DsrMK systems, the *dsrMK* gene cluster was used as the prediction input, while for DsrMKJOP systems, the entire *dsrMKJOP* gene cluster was used.

For structure predictions within SOB, four phylogenetically distant representatives were selected from the sequence pool used for sequence conservation analysis, including two from *Chlorobiota* and two from *Pseudomonadota* (fig. S18A-D). For cable bacteria, one representative strain with a fully sequenced and closed genome was selected from each known genus, *Candidatus Electronema* and *Candidatus Electrothrix*, for structure prediction (fig. S18I-J).

Throughout the prediction process, heme *b* was used as the DsrM-associated cofactor due to limitations in the ligand types supported by the AF3 server. To assess structural variability within each prediction run, the five predicted models were aligned pairwise using Matchmaker, and RMSD values in Å over all residue pairs were reported.

## Data availability

Cryo-EM structures of the DsrMKJOP are available from the PDB under accession codes 9RWH (“as-isolated” sample), 9RWJ (“DsrC-trisulfide” sample), 9RWN (“reduced” sample), 9RWK (“turnover” sample, retracted state), and 9RWL (“turnover” sample, extended state). Maps and half-maps are available from the Electron Microscopy Data Bank (EMDB) under accession codes 54329 to 54337. Atomic coordinates from the AlphaFold 3 predictions are archived in Zenodo ^81^. The sequence alignment files and other data used for the construction of figures are also archived in Zenodo ^82^.

## Acknowledgements

The Mechanistic Structural Biology group gratefully acknowledges the Central Electron Microscopy Facility at the Max Planck Institute of Biophysics for providing access to cryo-EM instrumentation and expert technical support. We thank Rita Zimmermann for her skilled assistance with anaerobic experiments; Yonca Ural-Blimke and Werner Kühlbrandt for their contributions to the initial sample screening; and José Guadalupe Rosas Jiménez and Rolf Thauer for helpful discussions. The Bacterial Energy Metabolism group thanks the research facilities at ITQB NOVA for their support, and in particular João Carita for bacterial growth and Ricardo Gomes (UniMS) for MS data acquisition and analysis.

## Author contributions

RMB, ACCB, and AIP purified the proteins. MDY prepared cryo-EM samples, collected and analysed cryo-EM data. RMB and ACCB carried out the mutagenesis and growth experiments. RMB performed the mass spec. studies. IACP and BJM acquired funding and supervised the experiments. MDY, RMB, IACP and BJM wrote the manuscript. All authors discussed the results and commented on the manuscript.

