## Supplementary Information for "Structural basis of sulfide production in dissimilatory sulfur metabolism"

| Bonnie J. Murphy | Inês A. C. Pereira |
|---|---|
| | |
| +49 69 6303 3210 | +351 214469327 |
| ORCID 0000-0001-6341-9368 | ORCID 0000-0003-3283-4520 |

† These authors contributed equally to this work.

‡ Present address: Ecole Polytechnique Fédérale de Lausanne (EPFL), Laboratory of Molecular Nanodynamics, CH-1015 Lausanne, Switzerland

§ Present address: Escola de Psicologia e Ciências da Vida, Universidade Lusófona de Humanidades e Tecnologias, Campo Grande 376, 1749-024 Lisboa, Portugal

**Supplementary information – Table of contents:**

Supplementary Discussion

Figs. S1 to S35

Tables S1 to S5

Movies S1

### Supplementary Discussion

#### Sequence, structural, and evolutionary diversification of DsrM and associated complexes

Sequence and structure comparisons within the DsrM family reveal functionally important divergence linked to both membrane-complex architecture and sulfur metabolism. These comparisons are elaborated here to provide mechanistic and evolutionary context for the DsrM classification introduced in the main text. We begin by analyzing DsrM diversification in SRM before extending the comparison to SOB and cable bacteria (see Supplementary Figure S35).

The minimal transmembrane complex required for sulfite reduction is DsrMK, a complex that is related to the methanogenic membrane-bound heterodisulfide reductase HdrDE and is suggested by large-scale phylogenetic analysis to have an archaeal origin. Later in evolutionary time, additional subunits J, O, and P were incorporated into the complex [1]. As a result of this diversification, extant SRM encode DsrMK or DsrMKJOP, and a few encode both types of complexes [1]. Here we refer to DsrM proteins associated with DsrMK complexes as DsrM-1, and those associated with DsrMKJ(OP) complexes as DsrM-2.

Our structural analysis of *A. fulgidus* DsrM-2, including comparison with *E. coli* NarI, a well-characterized DsrM homolog, reveals a fundamental divergence in quinol-site architecture. In our DsrM-2 structure, we identified a loop insertion near the region equivalent to the binding pocket for the competitive inhibitor pentachlorophenol (PCP)- in NarI, which is widely believed to be the quinol binding-site for this family of proteins. For convenience, we refer to this region as the "PCP site" in DsrM. This insertion interrupts the transmembrane helix TMH3 and produces a short helical segment, which occludes the "PCP site" in *A. fulgidus* DsrM-2 (fig. S14A-B). The inserted region contains a proline and a glycine, known to destabilize helices, which likely cause the TMH3 discontinuity (fig. S15A). In addition to its effect on the "PCP site", the loop insertion is also necessary to create the interaction interface for DsrJ docking. The discontinuity of TMH3 prevents a steric clash of DsrM with DsrJ (fig. S14C). Three arginines, Arg118, Arg121 and Arg198, located near the DsrM-DsrJ interface and the "PCP site", are strictly conserved in DsrM-2 (fig. S15B) and play a role in stabilizing DsrJ binding. Arg118 forms electrostatic interactions with Glu138 on the short helical segment (fig. S15A), stabilizing the loop structure. Arg121 and Arg198 interact electrostatically with a DsrJ heme *c* propionate, while Arg118 and Arg198 also contact separate propionates of the periplasmic heme *o* in DsrM (fig. S15A). These charge-charge interactions likely stabilize DsrJ docking and may mitigate charge repulsion between propionates of the heme *o* and heme *c*. Two of the three arginines, Arg118 and Arg121, occlude the "PCP site" in DsrM-2 (fig. S13B).

These features appear to reflect a general sequence pattern in the divergence between DsrM-1 and DsrM-2. Alignment of 78 phylogenetically diverse DsrM sequences (see methods), representative of a larger dataset [1], reveals that the loop insertion is a consistent feature of DsrM-2 and is absent in DsrM-1 (fig. S15C). Following the *A. fulgidus* DsrM-2 numbering, Arg198 is strictly conserved in DsrM-2, but not in DsrM-1 (fig. S15B). Arg118 is replaced with a conserved glycine in DsrM-1, while Arg121 is replaced by glycine or other small residues in DsrM-1 (fig. S15B). The much smaller glycines likely render the "PCP site" accessible to quinol in DsrM-1. Glu138, which forms an electrostatic interaction with Arg118, is conserved as aspartate in DsrM-2 but not in DsrM-1 (fig. S15C).

We asked whether these sequence differences mirror a structural and functional difference between DsrM-1 and DsrM-2, as observed between *A. fulgidus* DsrM-2 and *E. coli* NarI (DsrM-1-like). We examined AF3-predicted structures of four phylogenetically distant representatives of each group, which consistently showed an occluded "PCP site" in DsrM-2 proteins and an accessible "PCP site" in DsrM-1 (figs. S16F-I, and S17E-H). Notably, the elongated hydrophobic cavity, occupied by a menaquinone-7 (MK-7) in our DsrM-2 structure, appears conserved across predicted DsrM-1 and DsrM-2 structures (figs. S16F-I, and S17E-H). However, due to the lack of surrounding charged residues, this site is unlikely to host a substrate quinone, which would require proton transfer for turnover.

We also evaluated DsrM sequences in organisms beyond classical SRM, including sulfur oxidizers using the reverse Dsr pathway. The structural features previously described for DsrM-2 in reductive systems are also conserved in the sulfur-oxidizing *Chlorobiota*, and cable bacteria DsrM (here named DsrM-2a) (fig. S17, 18A-B, I-J). *Chlorobiota* are believed to have acquired the *dsr* genes by lateral gene transfer from SRM [1,2]. In contrast, DsrM-2 proteins from sulfur-oxidizing *Pseudomonadota* (here named DsrM-2b), whose *dsr* genes are phylogenetically more distantly related to SRM, lack the TMH3 loop interruption and instead harbor a conserved proline within TMH3. AlphaFold structural models of DsrM-2b proteins confidently predict that this proline introduces a kink within TMH3, also leading to occlusion of the "PCP site" (fig. S18C-D). Both DsrM-2a and DsrM-2b subclasses share the conserved arginine (or lysine) residues (figs. S15B, S18M) that contribute to "PCP-site" occlusion and stabilization of DsrM-DsrJ interactions, which provides further evidence that "PCP site" occlusion occurs in both DsrM-2a and DsrM-2b proteins.

Taken together, our analysis suggests that evolutionary remodeling of DsrM-2 led to both occlusion of the "PCP site" and creation of an interaction interface for DsrJ. We have shown that, in DsrMKJOP from *A. fulgidus*, DsrP harbors a separate quinol-binding site where menadiol binds, acting as electron donor for DsrC-trisulfide reduction (Fig. 2H). Accordingly, our analysis suggests that DsrMK complexes (containing DsrM-1) couple DsrC-trisulfide reduction to quinol oxidation at the "PCP site", whereas DsrMKJOP complexes (containing DsrM-2) link sulfur metabolism to quinol chemistry at DsrP rather than at DsrM.

This architectural shift raises the question of what physiological advantage is conferred by replacing one quinol-binding site ("PCP site") with another (DsrP site), given that both sites appear to exchange protons with the periplasm. One possibility is that DsrP couples quinol oxidation to vectorial proton translocation, as discussed in the main text. Although our structural data do not reveal an obvious well-connected transmembrane proton pathway in *A. fulgidus* DsrP, they also do not exclude the possibility of such a pathway, and this question requires further experimental investigation.

In cable bacteria, the Dsr complex appears to be composed only of DsrMKJ subunits [3], and the sequence signature is of a DsrM-2a protein, corresponding to an occluded "PCP site". Although *dsrOP*-like genes are present in cable bacteria genomes (named *$dsrO_hP_h$*), they are encoded at a locus separate from *dsrMKJ*, and co-localize with the gene for an additional tetraheme cytochrome *c*, and have been suggested to be part of a different membrane complex (named $DsrO_hP_h$Cyt) with unknown physiological function [3]. Structural modeling of the DsrMKJ and $DsrO_hP_h$Cyt complexes with AF3 gives putative complexes that appear reasonable and show similar architecture to the corresponding subcomplexes in our structure (fig. S19A). However, superposing these subcomplexes together as in the *A. fulgidus* DsrMKJOP architecture results in extensive steric clashes (fig. S19B). Predictions of a $DsrMKJO_hP_h$Cyt complex were highly variable between different species and did not appear reasonable in terms of protein-protein interface stability or cofactor placement. Compared with *A. fulgidus* DsrOP, these $DsrO_hP_h$Cyt complexes contain additional α-helical elements in $DsrO_h$ and $DsrP_h$ that sterically clash with DsrJ and DsrM (fig. S19C), precluding formation of a canonical $DsrMKJO_hP_h$ assembly. This divergence in complex architecture is mirrored at the level of DsrJ heme coordination: whereas *A. fulgidus* DsrJ harbors a *c*-type heme coordinated by a histidine from DsrO (fig. S19D), in cable bacteria the equivalent heme is structurally predicted to be coordinated only by DsrJ due to an additional histidine residue in its sequence (fig. S19E). These coordination modes are mutually exclusive: DsrJ of non-cable bacteria encodes an arginine in the position of this additional histidine (fig. S19D, G), while cable bacteria $DsrO_h$ lacks the histidine (fig. S19F, G). Overall, these structural observations fully support the recent proposal that the DsrMKJ and $DsrO_hP_hCyt$ proteins function independently in cable bacteria [3]. Consistent with this, AF3-predicted structures show that $DsrO_hP_hCyt$ from cable bacteria closely resembles homologous complexes from *gamma-* and *betaproteobacteria* that lack DsrMKJ and DsrAB entirely. The predicted DsrMKJ complex of cable bacteria with a blocked quinol-binding site in DsrM-2a, suggests that activity at DsrK is not associated with electron exchange with the quinol/quinone pool, therefore, a separate (currently unknown) partner would be required for activity. Further studies will be required to elucidate the role of this unique Dsr architecture.

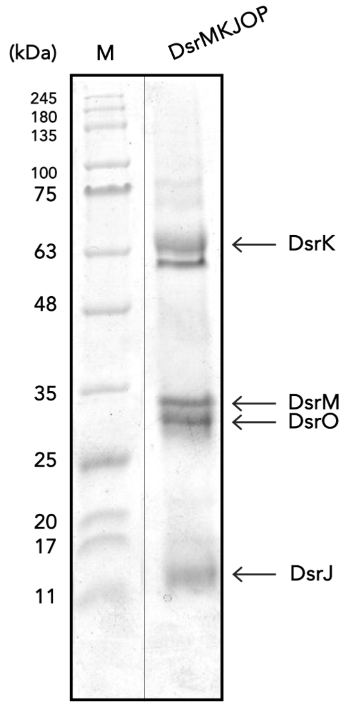


**Fig. S1. Tricine-SDS-PAGE of the "as-isolated" *A. fulgidus* DsrMKJOP.** The purified *A. fulgidus* DsrMKJOP complex (25 µg) was run in a 10% gel, stained with Coomassie blue, and then heme-stained. All subunits were identified by MS except DsrP, likely undetected due to its highly hydrophobic nature, which may cause protein aggregation and prevent migration into the gel.

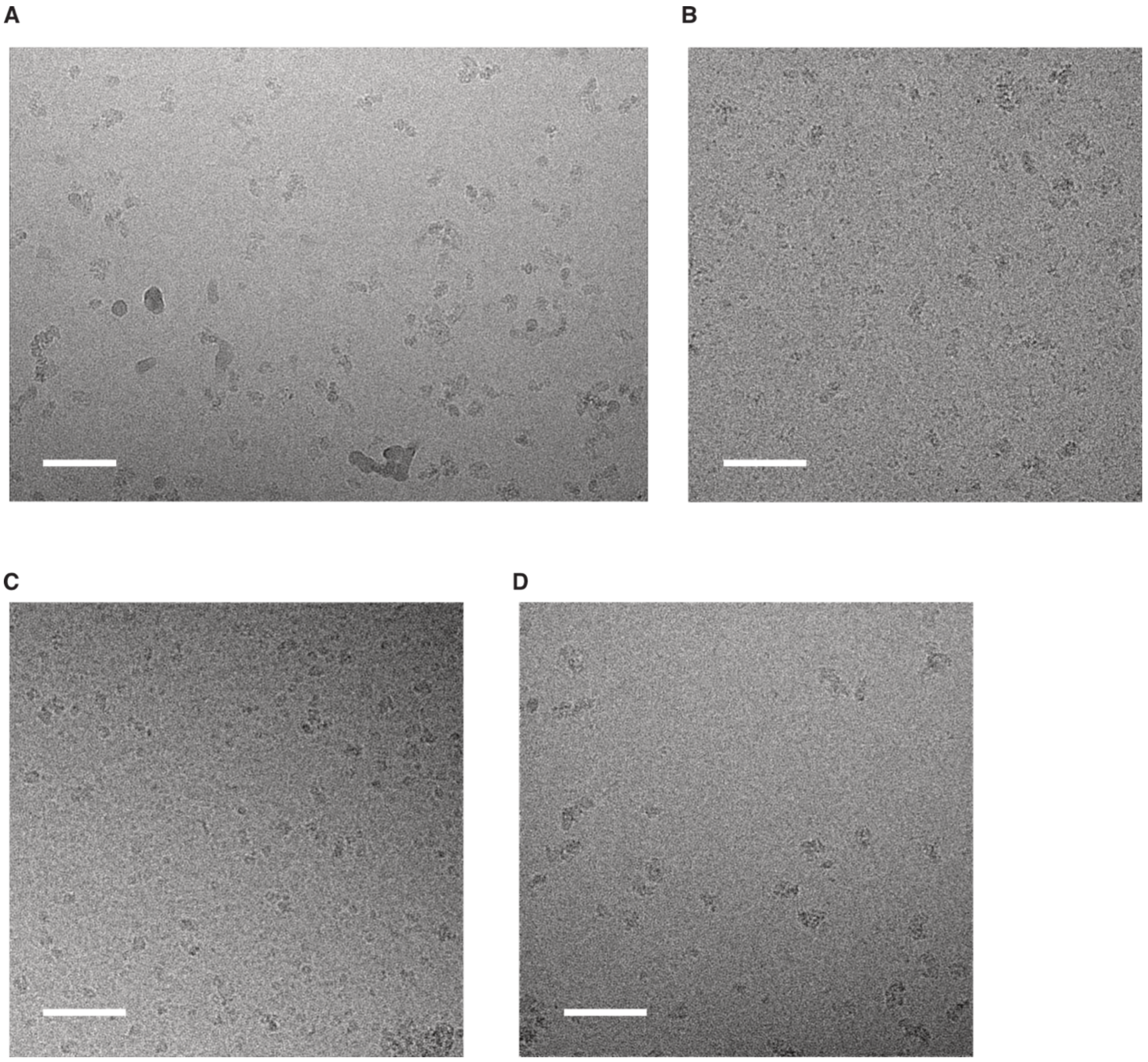

**Fig. S2. Representative micrographs of DsrMKJOP samples under different conditions**. **(A)** “As-isolated” sample, **(B)** “DsrC-trisulfide” sample, **(C)** “reduced” sample, and **(D)** “turnover” sample. All micrographs were lowpass-filtered to 20 Å and displayed at 0 σ contrast. Scale bars represent 500 Å.

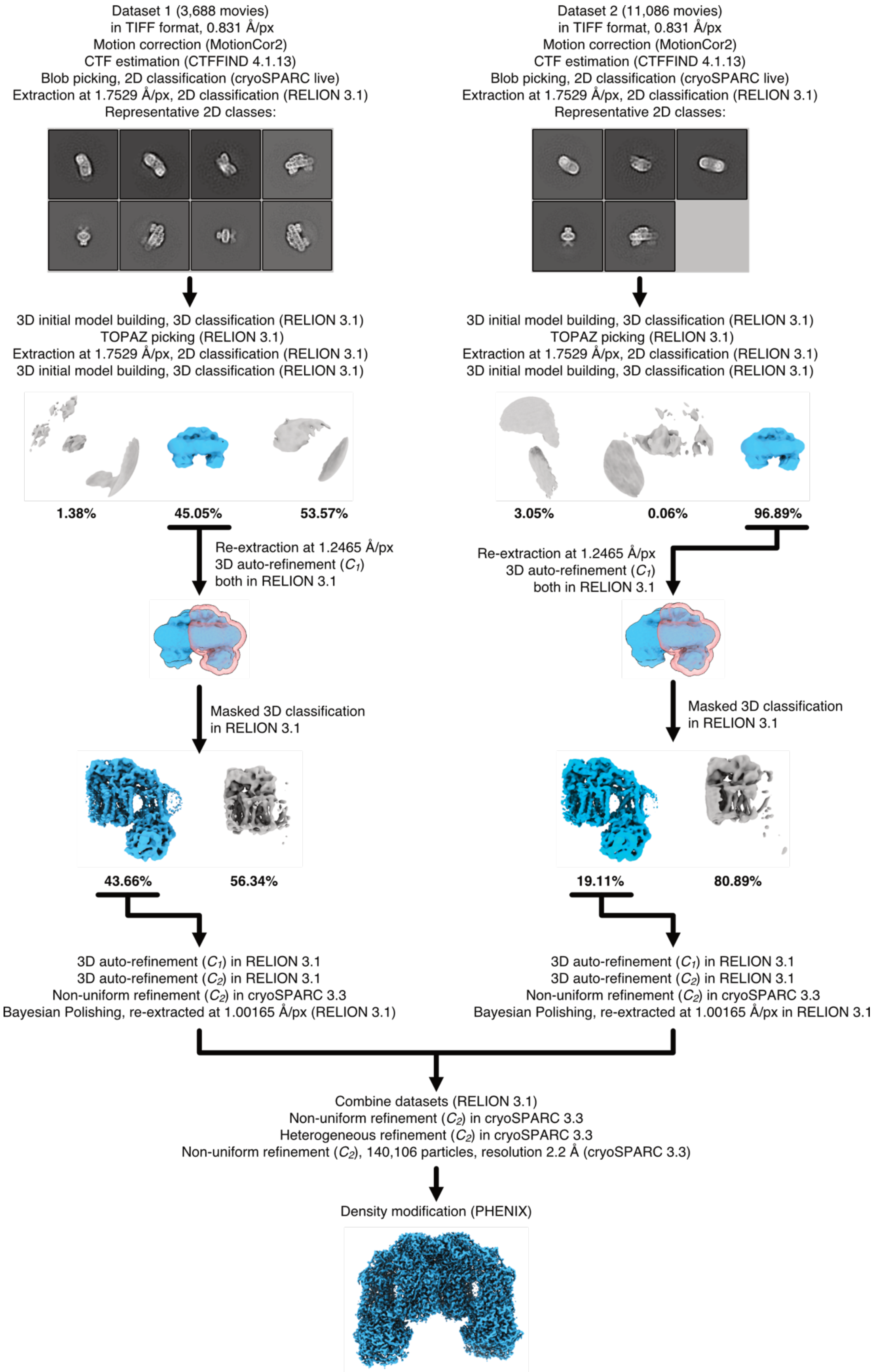


**Fig. S3. Cryo-EM processing workflow for the "as-isolated" sample.** The software packages used for each processing step are indicated. The reported resolution was determined using the gold-standard FSC at the 0.143 threshold following refinement in cryoSPARC. For each 3D classification step, the proportions of particles contributing to each class are shown. Map densities from particles selected for further processing are shown in blue, while those from discarded particles are shown in gray.

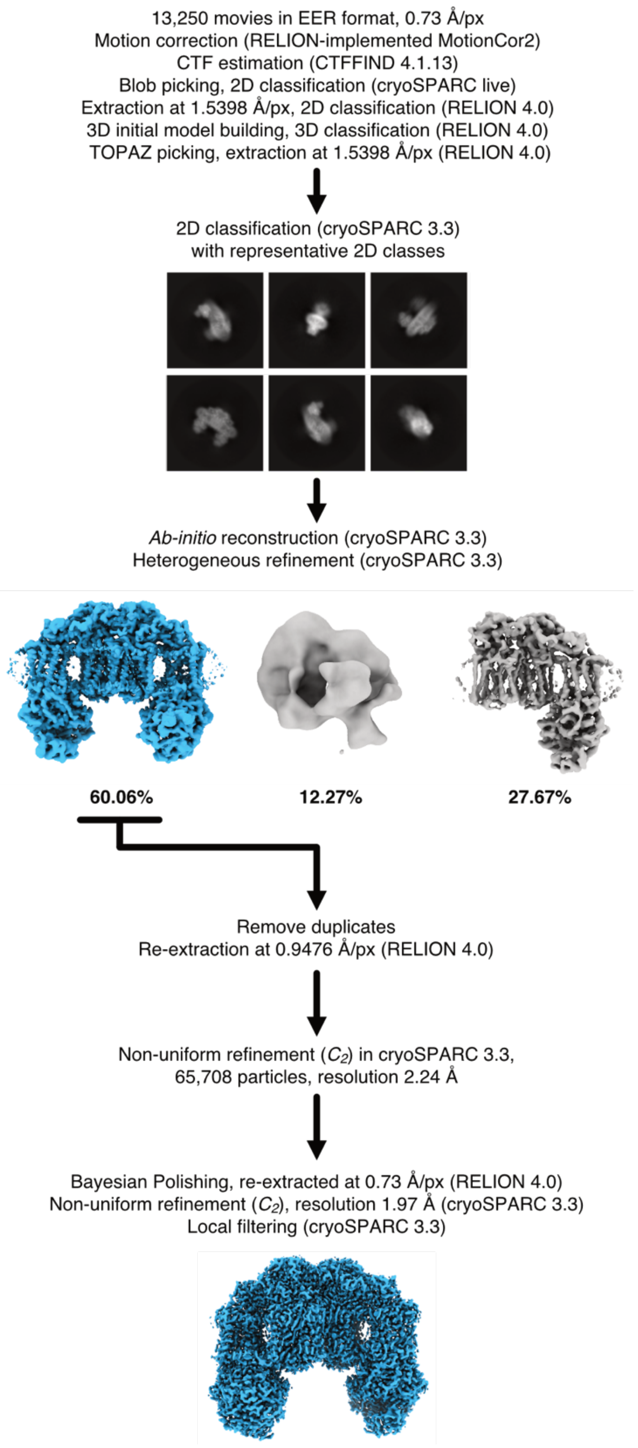


**Fig. S4. Cryo-EM processing workflow for the "DsrC-trisulfide" sample.** The software packages used for each processing step are indicated. The reported resolutions were determined using the gold-standard FSC at the 0.143 threshold following refinement in cryoSPARC. For the heterogeneous refinement step, the proportions of particles contributing to each class are shown. Map densities from particles selected for further processing are shown in blue, while those from discarded particles are shown in gray.

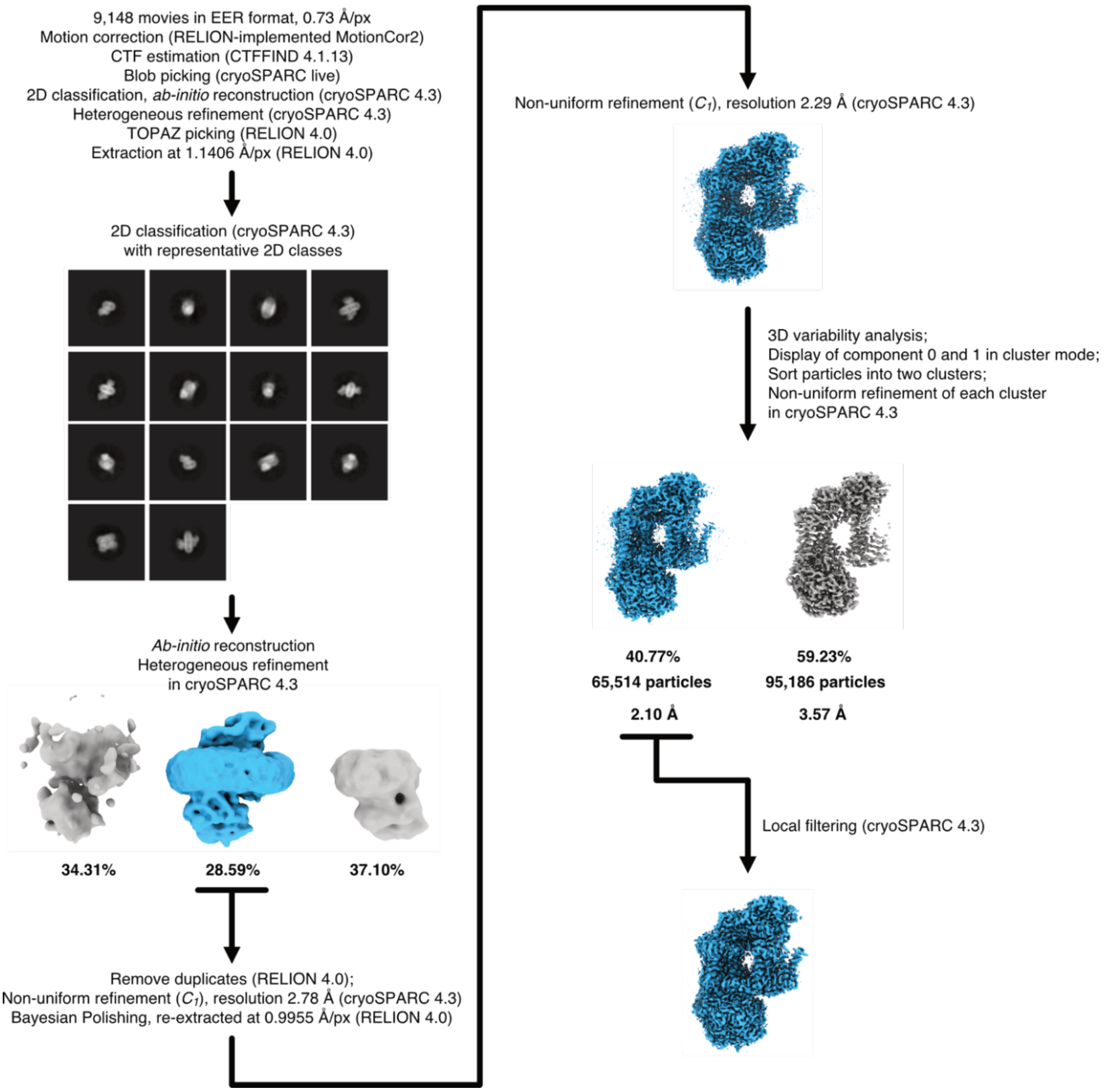


**Fig. S5. Cryo-EM processing workflow for the "reduced" sample.** The software packages used for each processing step are indicated. The reported resolutions were determined using the gold-standard FSC at the 0.143 threshold following refinement in cryoSPARC. For each heterogeneous refinement and 3DVA step, the proportions of particles contributing to each class or cluster are shown. Map densities from particles selected for further processing are shown in blue, while those from discarded particles are shown in gray.

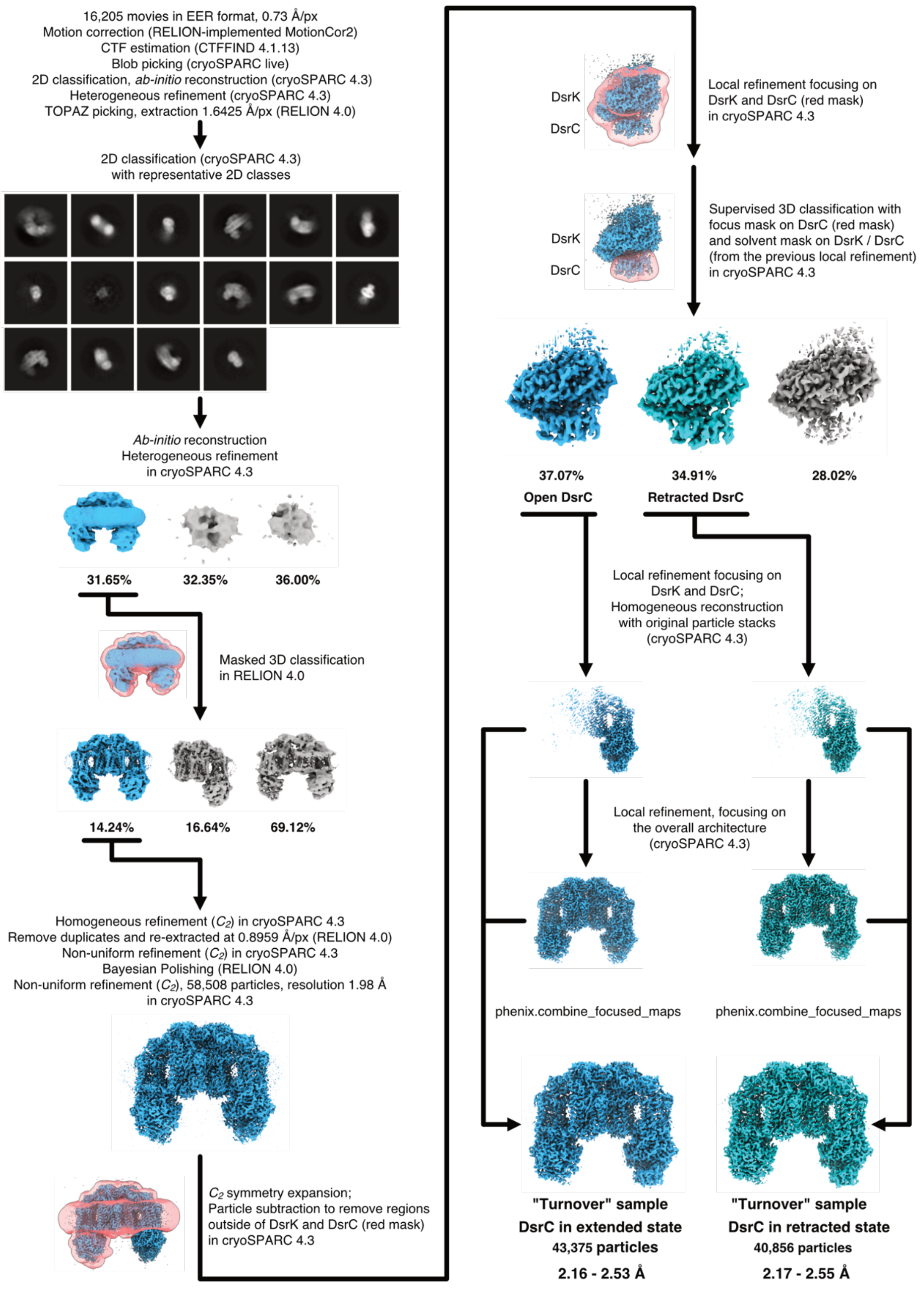


**Fig. S6. Cryo-EM processing workflow for the "turnover" sample.** The software packages used for each processing step are indicated. For each 3D classification and heterogeneous refinement step, the proportions of particles contributing to each class are shown. Map densities from particles selected for further processing are shown in blue, while those from discarded particles are shown in gray. The supervised 3D classification used to separate different conformational states of DsrC is further illustrated in fig. S27. Following this step, maps corresponding to the extended state are colored blue, while those corresponding to the retracted state are colored cyan. The reported resolutions were determined using the gold-standard FSC at the 0.143 threshold following refinement in cryoSPARC. Resolutions of the composite maps, which reflect the range of resolutions from their respective component maps, are also reported, along with the number of particles contributing to each state.

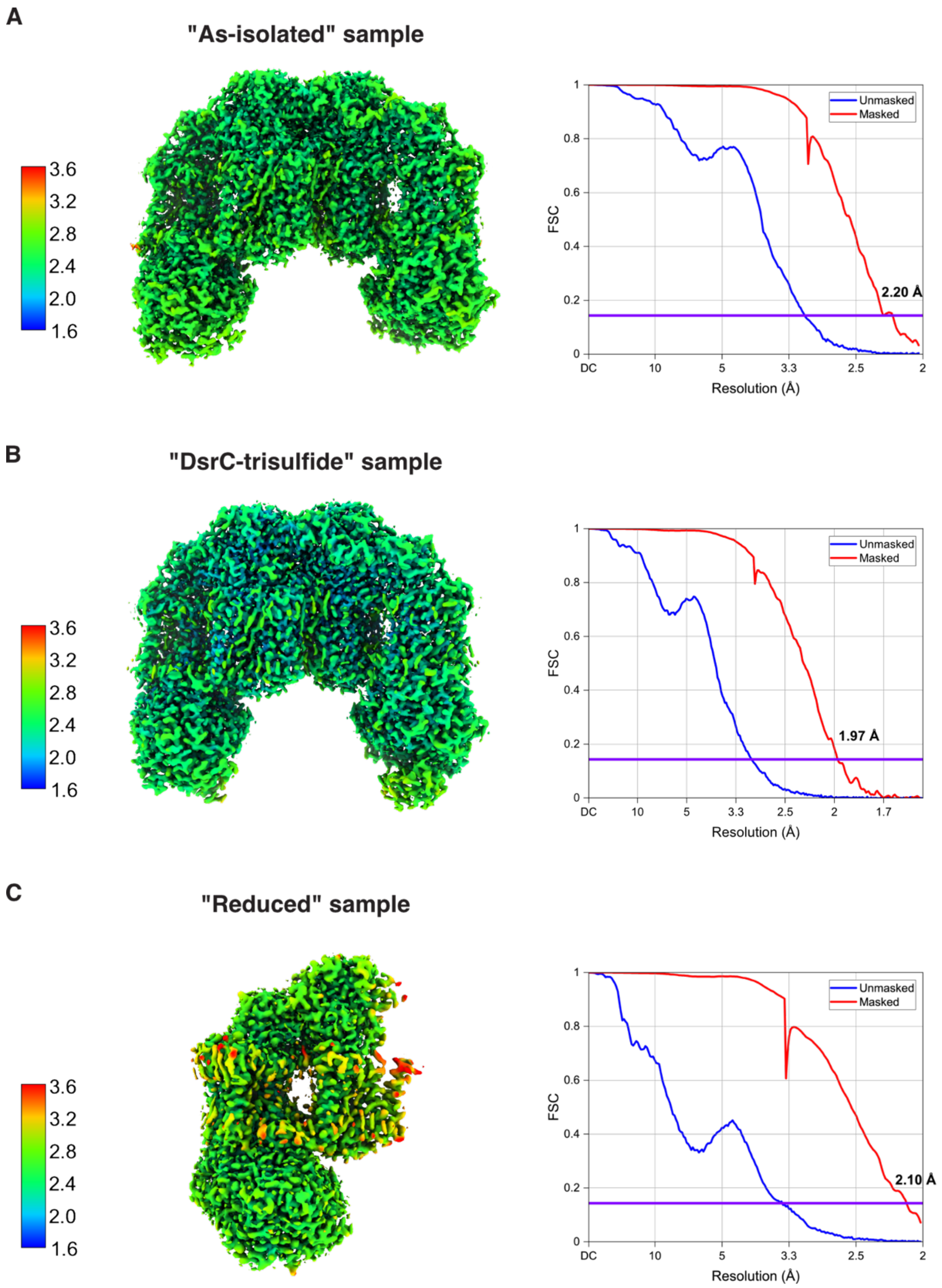


**Fig. S7. Local resolution estimates and FSC curves.** Cryo-EM maps of DsrMKJOP from the "as-isolated" **(A)**, "DsrC-trisulfide" **(B)**, and "reduced" **(C)** samples were colored according to local resolution, estimated using cryoSPARC. FSC curves between independent half-maps are plotted for each condition. Reported resolutions were determined using the gold-standard FSC criterion at the 0.143 threshold, indicated by purple horizontal lines.

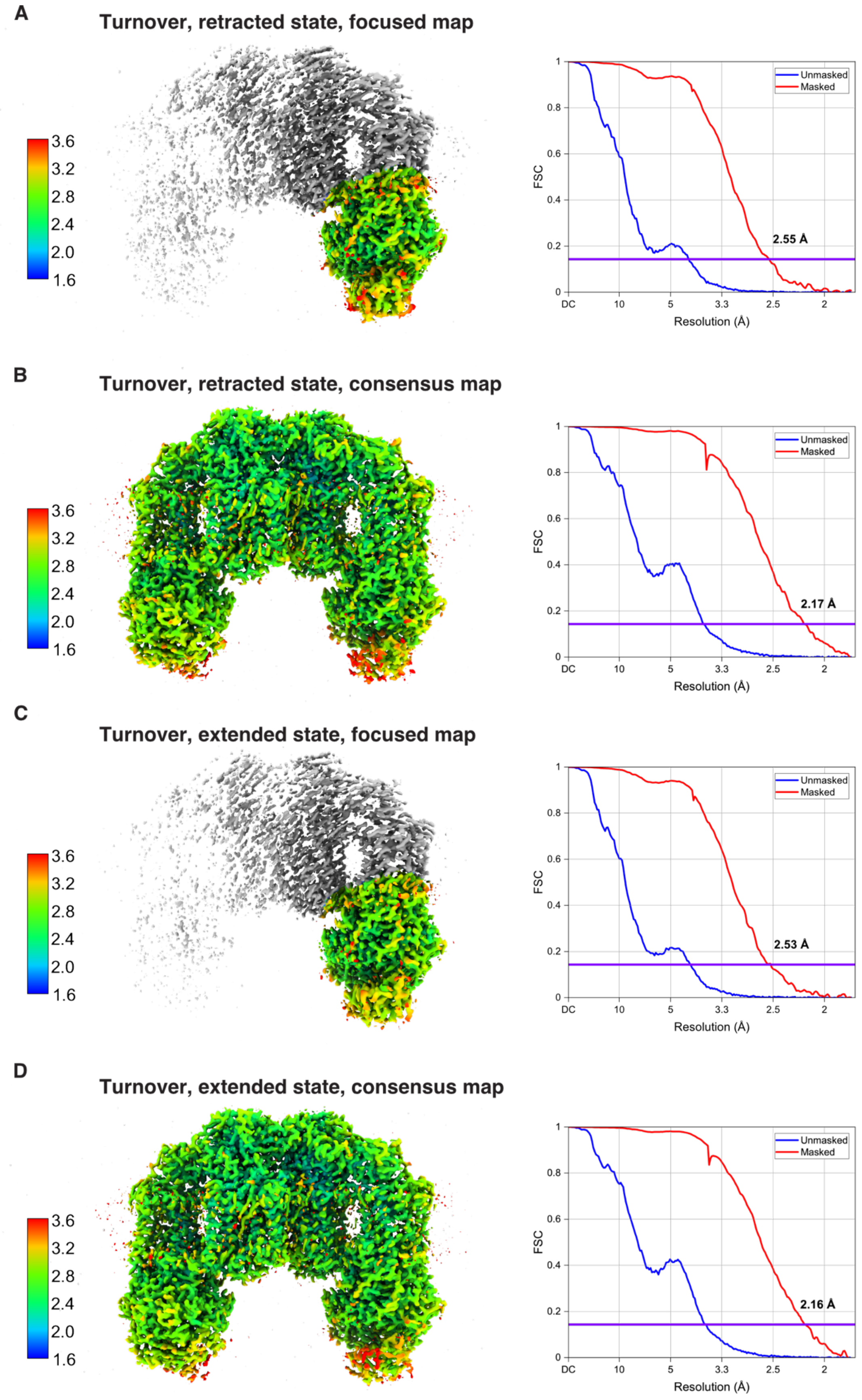


**Fig. S8. Local resolution estimates and FSC curves.** Cryo-EM maps from the "turnover" sample were colored by local resolution, estimated using cryoSPARC. FSC curves between independent half-maps are shown for: the focused map in the “retracted” state **(A)**, the consensus map in the “retracted” state **(B)**, the focused map in the “extended” state **(C)**, and the consensus map in the “extended” state **(D)**. Reported resolutions were determined using the gold-standard FSC criterion at the 0.143 threshold, indicated by purple horizontal lines. Local resolution in (A) and (C) was estimated only within the regions subjected to local refinement.

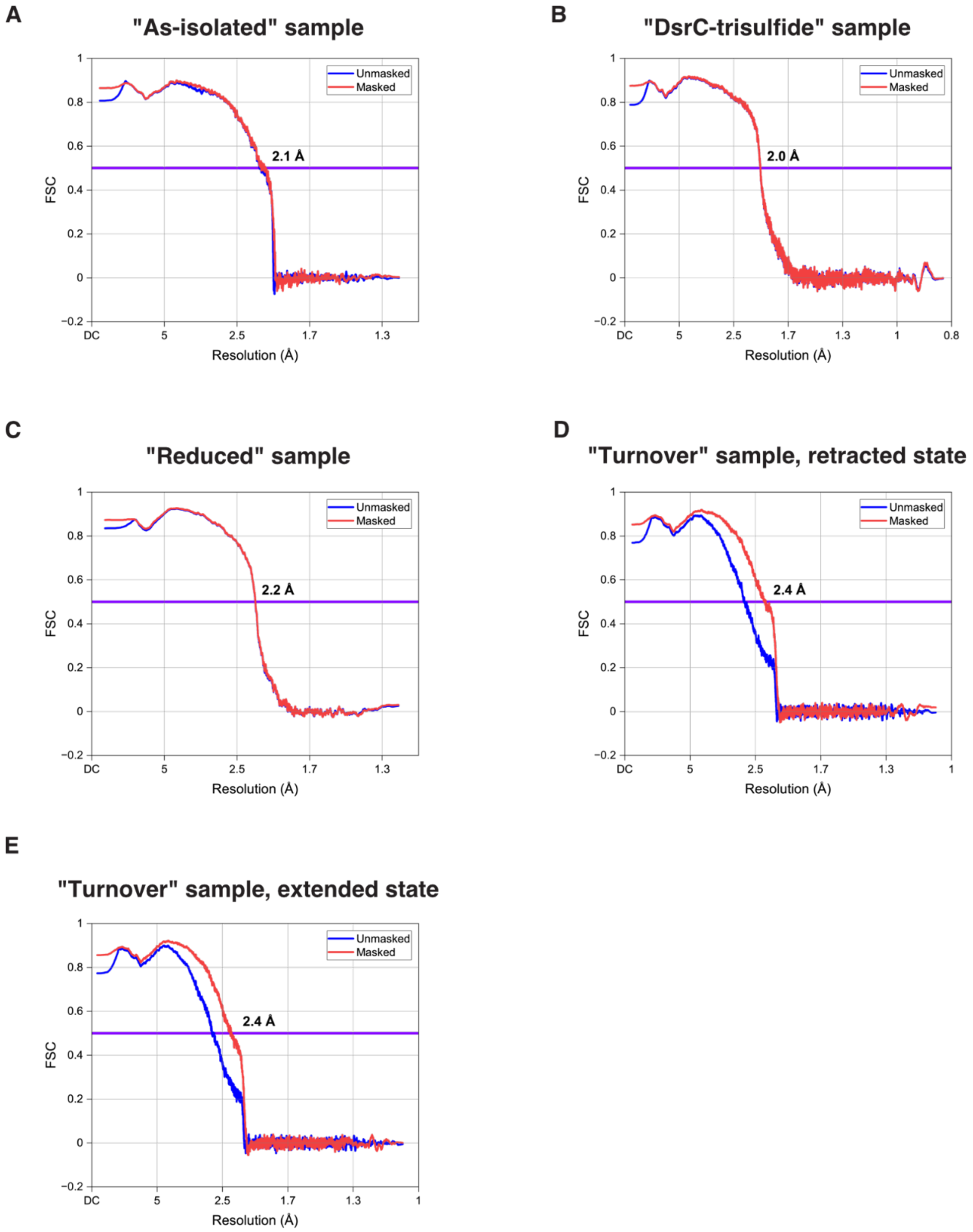


**Fig. S9. Model-map FSC curves.** FSC between refined atomic models and their corresponding cryo-EM maps are shown. Reported resolutions were determined at the 0.5 FSC threshold, indicated by purple horizontal lines, as part of the model validation procedure in *PHENIX*.

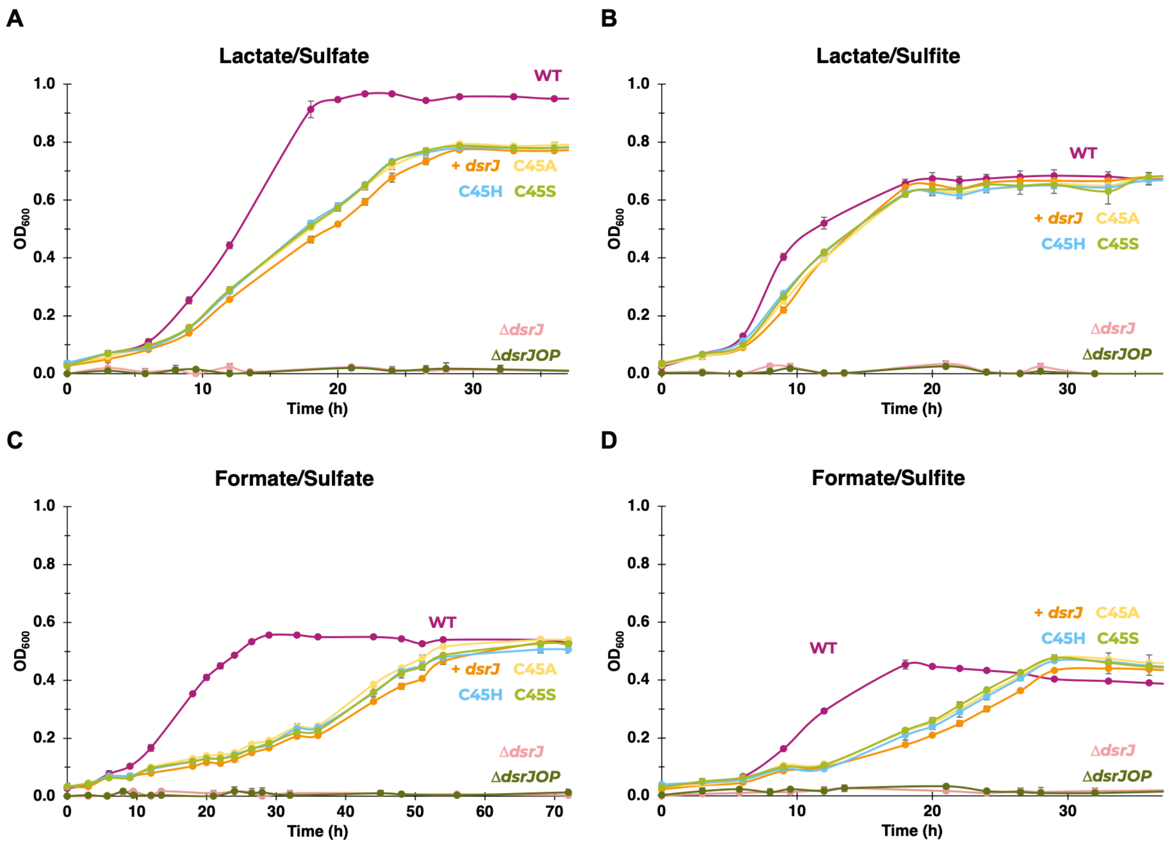


**Fig. S10. Growth curves of *N. vulgaris* WT and mutant strains under respiratory conditions.** *N. vulgaris* WT (magenta); *N. vulgaris* Δ*dsrJOP* (dark green) and Δ*dsrJ* (pink) deletion strains; and *N. vulgaris* Δ*dsrJ* + pMO-*dsrJ* complemented strain (+ *dsrJ*, orange) along with *dsrJ* variant strains: C45A (yellow), C45H (blue), and C45S (green) were grown in MOY medium (0.2 g/L yeast extract) containing **(A)** 30 mM lactate/30 mM sulfate, **(B)** 15 mM lactate/10 mM sulfite, **(C)** 50 mM formate/30 mM sulfate, and **(D)** 50 mM formate/10 mM sulfite. Data points are mean ± SD, n = 3 independent biological experiments.

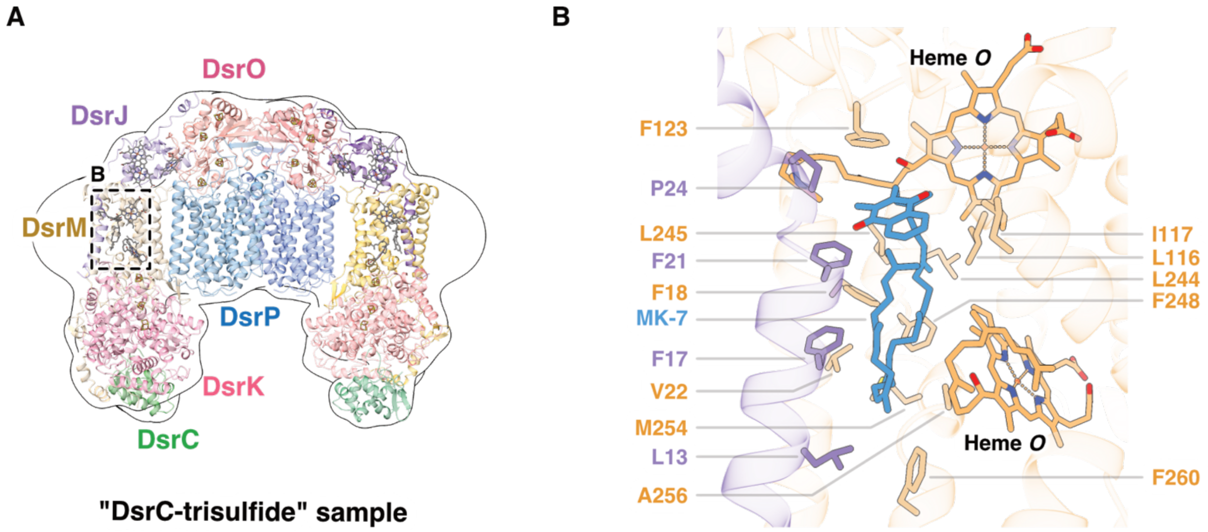


**Fig. S11. Copurified MK-7 is surrounded by nonpolar residues from DsrM and DsrJ. (A)** An overview of the DsrMKJOP complex from the "DsrC-trisulfide" sample, shown with the same color scheme as in Fig. 1. The cryo-EM map was Gaussian-filtered to 4 Å and is displayed as a transparent surface contoured at 5.7 σ. **(B)** A close-up view of the MK-7-binding pocket, shown with the same color scheme as in Fig. 2B. Hydrophobic residues from DsrM and DsrJ surrounding MK-7 are highlighted. Two nearby heme *o* cofactors within DsrM are also shown.

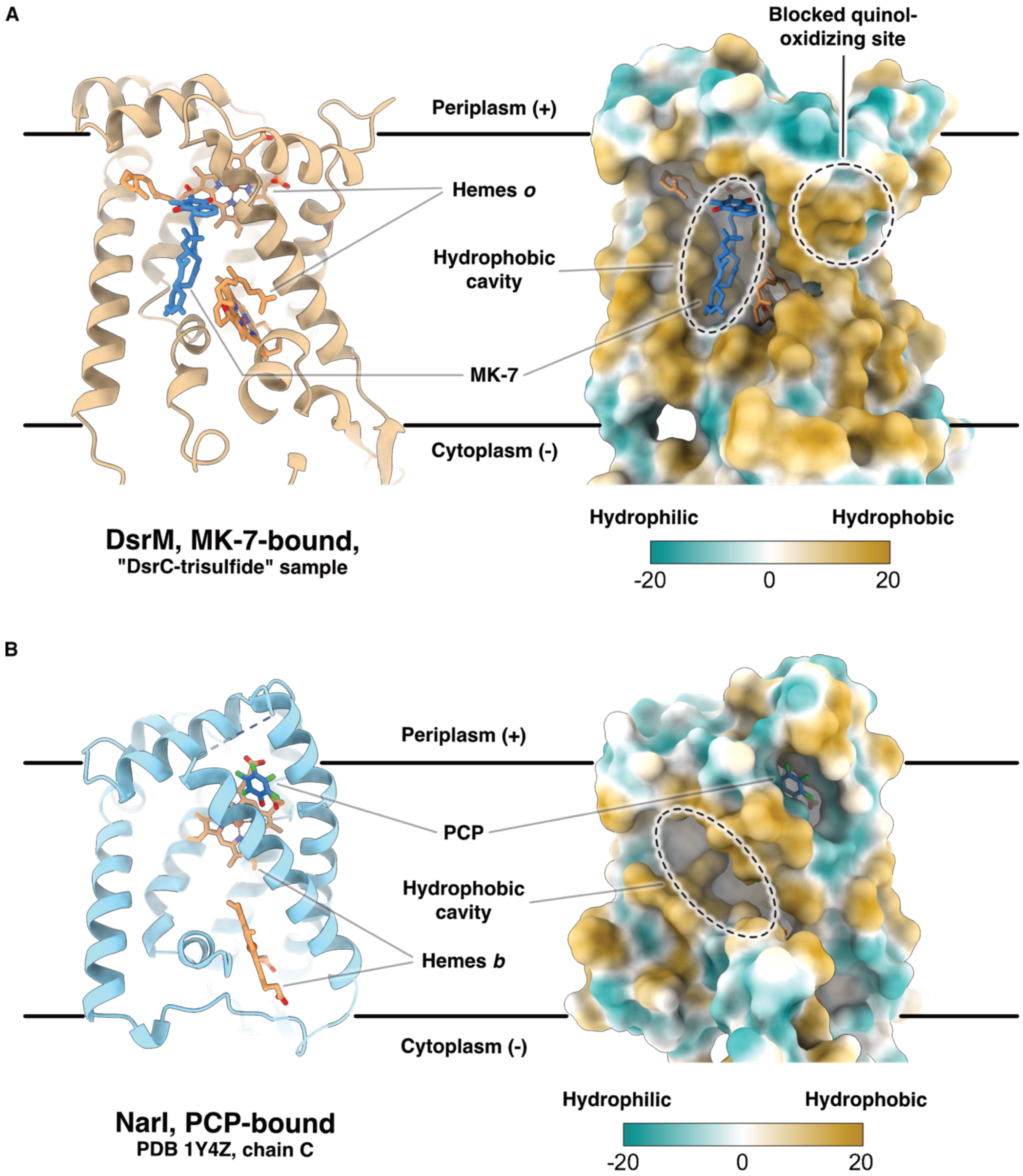


**Fig. S12. MK-7 and PCP occupy distinct binding pockets in DsrM and NarI, respectively.** **(A)** MK-7 binds within a hydrophobic cavity of DsrM ("DsrC-trisulfide" sample, chain m; tan cartoon), located between two hemes *o*. **(B)** In contrast, PCP binds to NarI (PDB ID 1Y4Z [4], chain C; light blue cartoon) in a separate pocket on the periplasmic side of the periplasmic heme *b*, which is likely the quinol-oxidizing site. The equivalent binding site in DsrM is blocked, whereas the hydrophobic cavity that accommodates MK-7 in DsrM remains unoccupied in NarI. MK-7, PCP, and heme cofactors are depicted as sticks and colored as follows: oxygen - red; nitrogen - blue; iron - dark orange; chlorine - green; and carbon - orange in hemes and steel blue in MK-7 and PCP, respectively. Molecular surfaces of DsrM and NarI were generated and colored in ChimeraX according to molecular lipophilicity potential, with a color gradient ranging from dark cyan (most hydrophilic) to dark goldenrod (most hydrophobic).

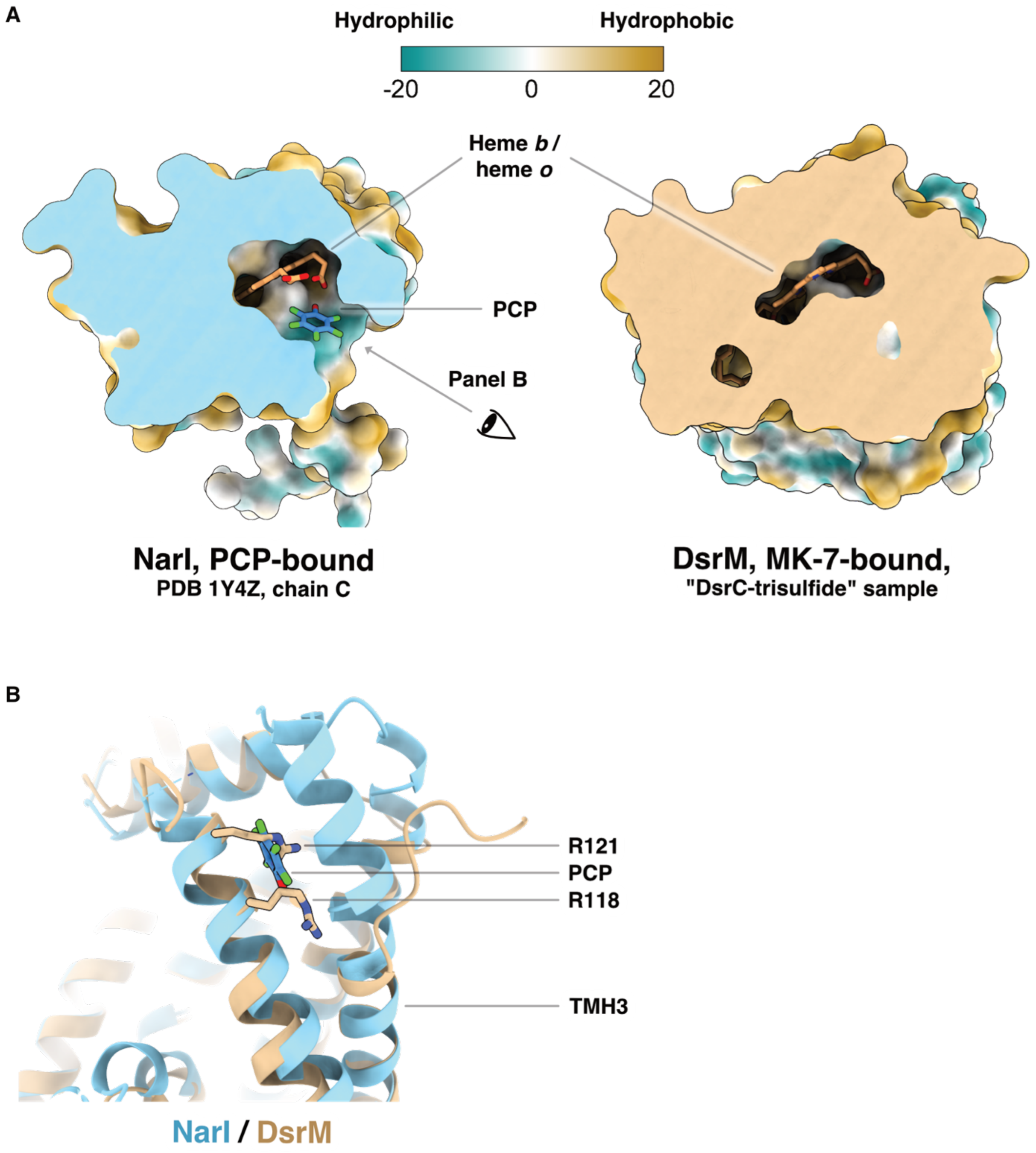


**Fig. S13. Two arginine residues block the quinol-oxidizing site in DsrM. (A)** Sliced views of the same NarI and DsrM structures shown in fig. S12 reveal that the site in DsrM corresponding to the PCP-binding pocket in NarI, which is likely the quinol-oxidizing site, is blocked. **(B)** Superposition of DsrM (tan cartoon) and NarI (light blue cartoon) shows that Arg118 and Arg121 in DsrM clash with PCP. In NarI, the equivalent positions to these two arginines are glycine residues, highlighting a key structural difference in this region. Hydrophobicity surfaces in (A) were generated as described in fig. S12.

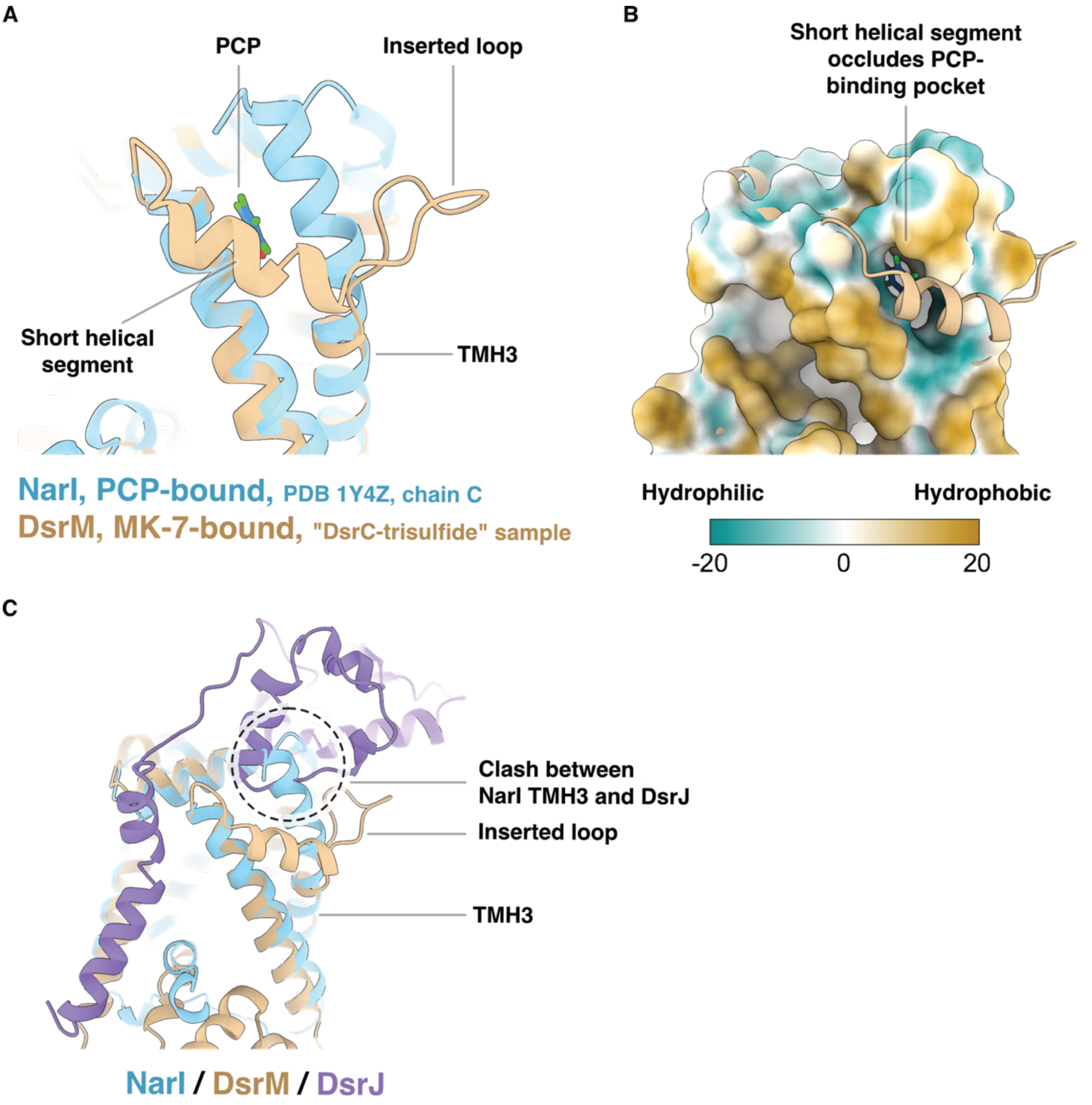


**Fig. S14. The short helical segment of TMH3 occludes the quinol-oxidizing site in DsrM.** **(A)** The same superposition of DsrM and NarI as in fig. S13B, now with the short helical segment of TMH3 visualized, reveals that this segment lies in front of PCP in NarI and directly occludes the PCP-binding pocket **(B)**. **(C)** This short helical segment is formed by a loop insertion in DsrM, which may help avoid a potential steric clash with DsrJ, as indicated by the dashed circle. The hydrophobicity surface in (B) was generated as described in fig. S12.

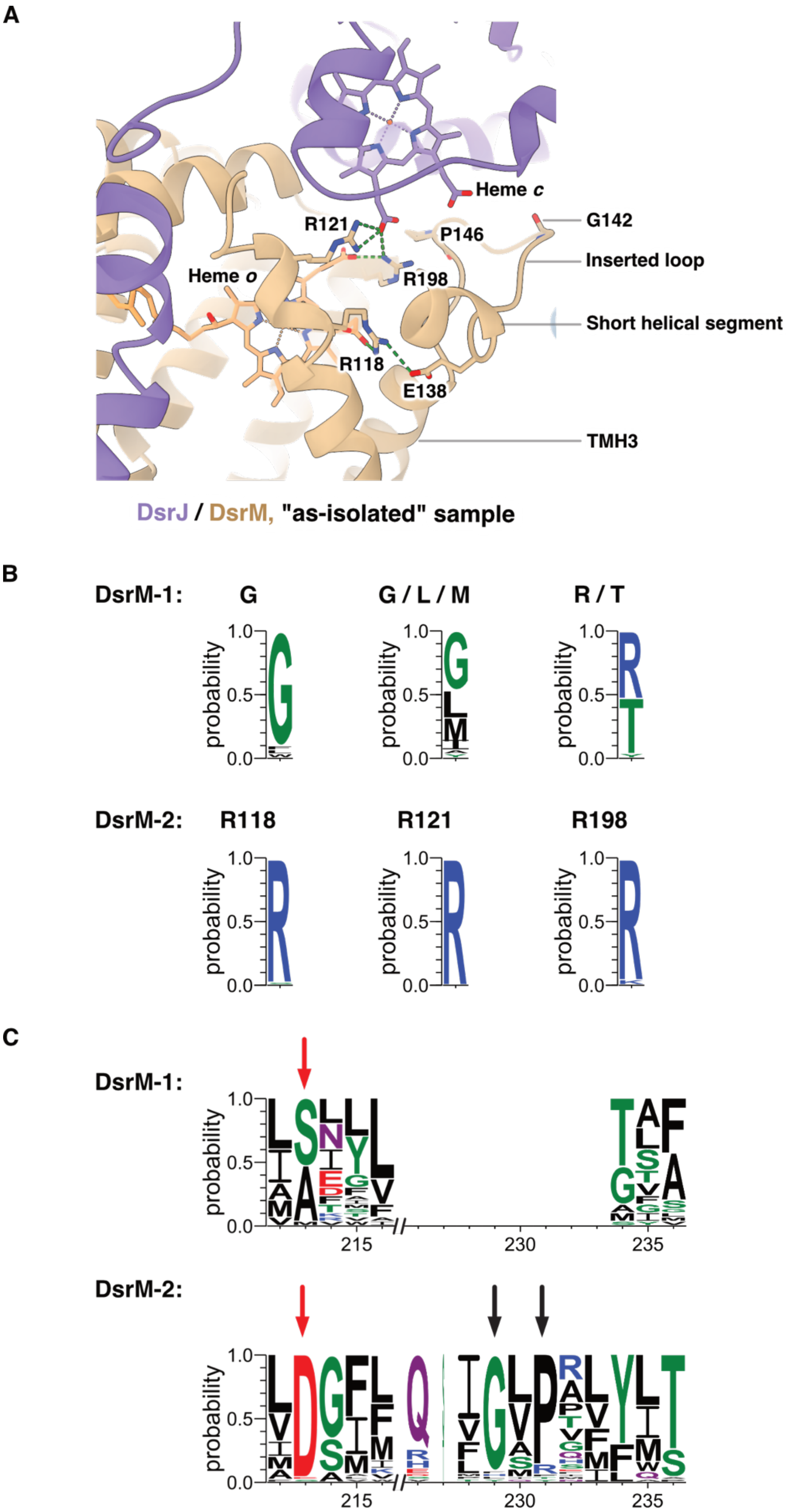


**Fig. S15. Structural adaptations in DsrM-2 facilitate interaction with DsrJ. (A)** In *A. fulgidus* DsrM, Arg118, Arg121, Glu138, and Arg198 form electrostatic interactions (green dashed lines) that stabilize both the short helical segment and the heme *c* in DsrJ located closest to DsrM. Gly142 and Pro146 disrupt the helical structure and promote loop formation in the inserted region. **(B)** Sequence conservation analysis shows that the three arginine residues in *A. fulgidus* DsrM are conserved in DsrM-2, whereas in DsrM-1, they are either substituted with glycine or show lower conservation. **(C)** In the sequence conservation plot, residues 212 to 236 in the consensus sequence (corresponding to residues 137 to 151 in *A. fulgidus* DsrM) reveal the absence of the loop insertion in DsrM-1, indicated by a gap. The residue corresponding to Glu138 (vertical red arrows) is conserved as aspartate in DsrM-2, preserving the negative charge, but appears as serine or alanine in DsrM-1. The conserved glycine and proline in DsrM-2 are marked by two black vertical arrows. Note that Gly142 in *A. fulgidus* DsrM does not align with the conserved glycine, but its presence within the loop region likely still contributes to the local destabilization of the helix. The color scheme in (A) matches that in Fig. 1. The consensus sequences in (B) and (C) are colored according to the chemical property of the residues. We note for clarity that the DsrM-2 proteins shown here fall within the DsrM-2a subclass defined later in the Supplementary Text.

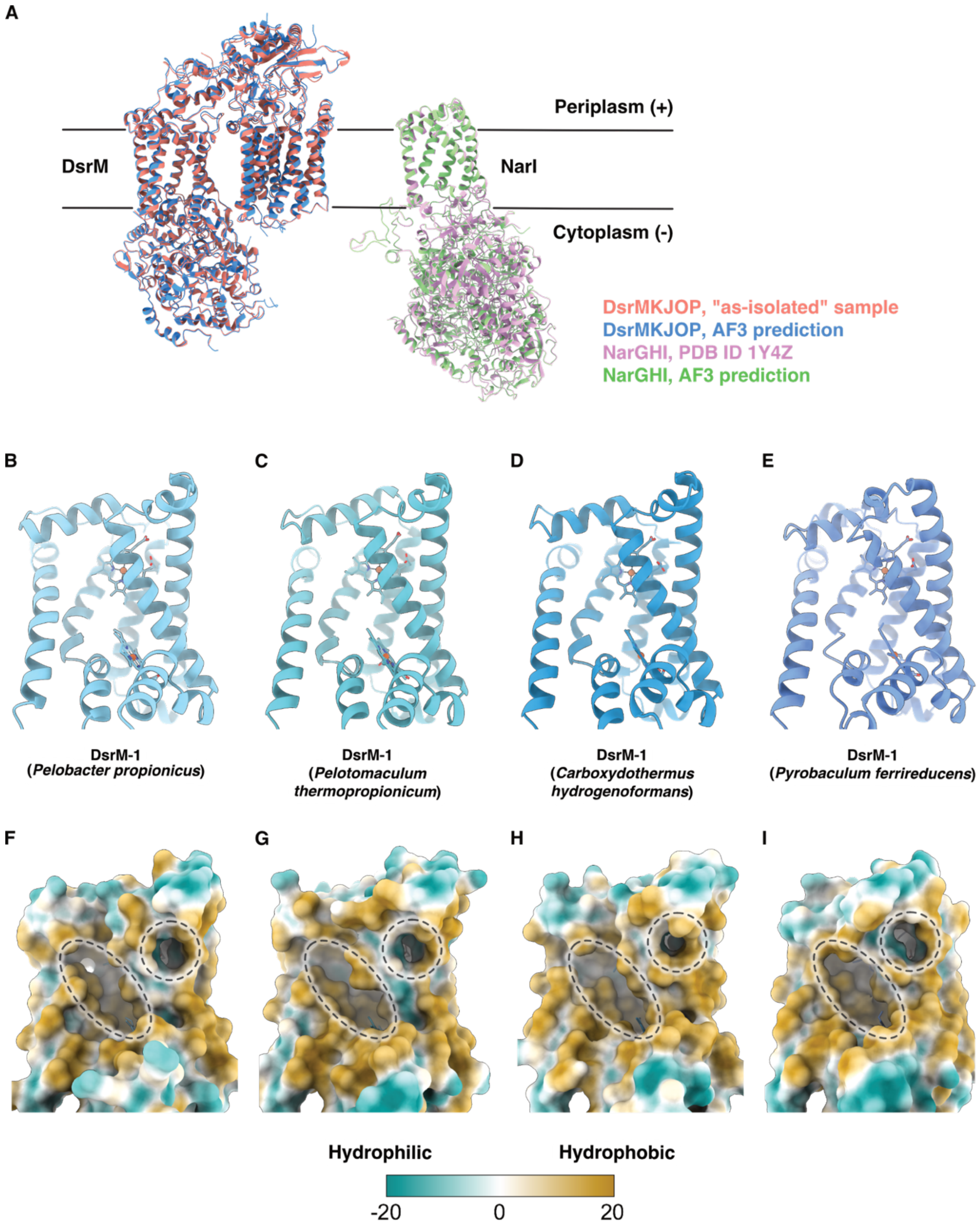


**Fig. S16. AF3-predicted structures reveal an accessible quinol-oxidizing site in DsrMK.** **(A)** In an initial validation step, AF3 predictions of DsrMKJOP and NarGHI were superimposed onto their respective experimental structures (DsrMKJOP: "as-isolated" sample; NarGHI: PDB ID 1Y4Z [4]), revealing close structural similarity. This resemblance is quantified by RMSDs, calculated between each of the five output models per prediction and the corresponding experimental structure, ranging from 0.391 Å to 0.425 Å for DsrMKJOP and from 0.192 Å to 0.208 Å for NarGHI. The pTM scores for these predictions remain constant across five models, with values of 0.87 for DsrMKJOP and 0.95 for NarGHI, indicating confident and accurate predictions. For simplicity, only the DsrMKJOP pentamer was included in the prediction and RMSD calculation. The first-ranked model from each prediction is shown. Following the validation step, AF3 was used to predict the DsrMK structures from four SRM organisms whose DsrM subunits are phylogenetically distant representatives of the DsrM-1 group, which includes proteins that belong to Dsr complexes that lack additional J, O, and P subunits. Cartoon and surface representations are shown for *Pelobacter propionicus* **(B, F)**, *Pelotomaculum thermopropionicum* **(C, G)**, *Carboxydothermus hydrogenoformans* **(D, H)**, and *Pyrobaculum ferrireducens* **(E, I)**. All four predicted DsrM homologs exhibit accessible quinol-oxidizing sites, highlighted by dashed circles, and also display elongated hydrophobic cavities (dashed ellipses) at positions equivalent to those in *A. fulgidus* DsrM-2, where a co-purified MK-7 is bound. Among the five output models per prediction, pairwise RMSDs range from 0.116 Å to 2.492 Å, and pTM scores from 0.86 to 0.89, supporting confident and high-quality predictions. The first-ranked model from each prediction is shown. DsrM cartoon models are shown in shades of blue, from light in (B) to dark in (E). Hydrophobicity surfaces of DsrM in (F-I) were generated as described in fig. S12. See methods for further details.

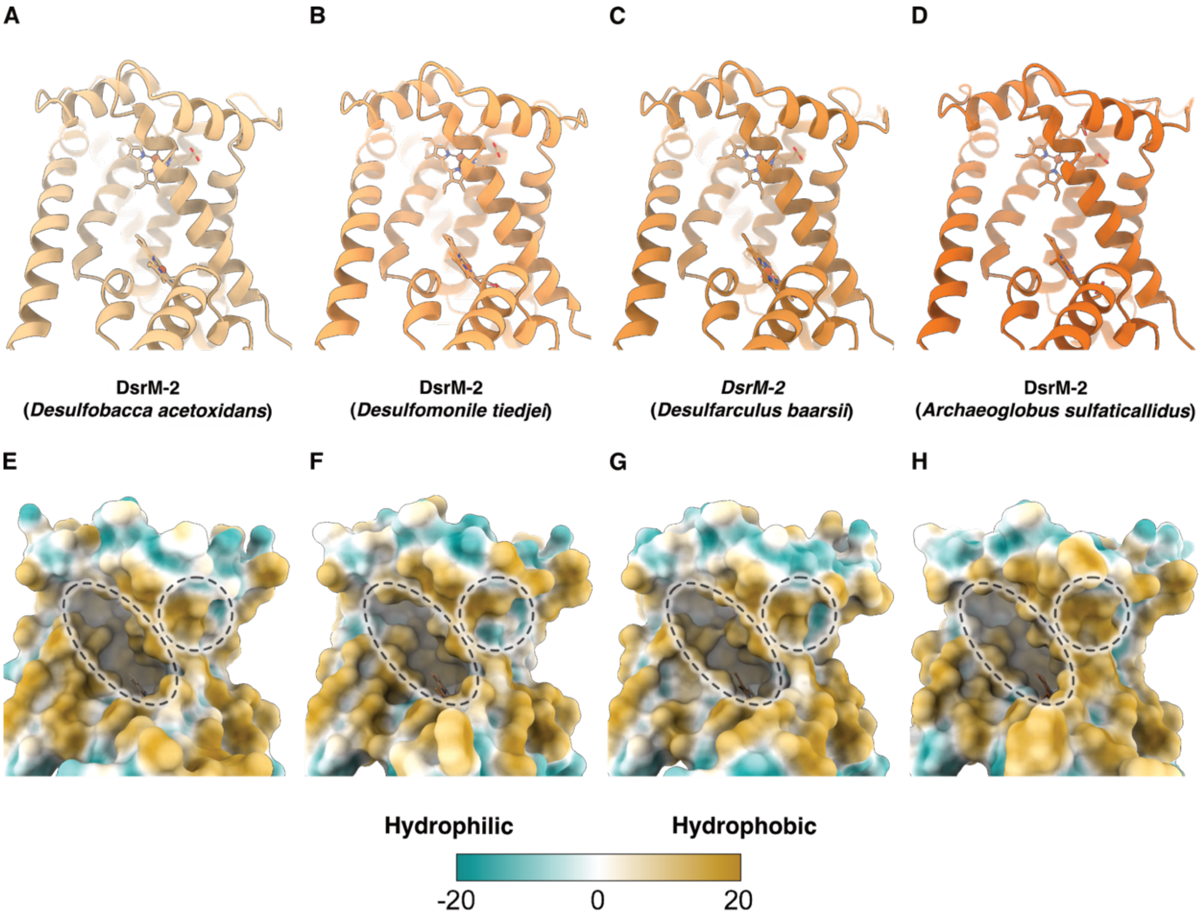


**Fig. S17. AF3-predicted structures reveal occluded quinol-oxidizing site in DsrMKJOP.** AF3 was used to predict the DsrMKJOP structures from four SRM organisms whose DsrM subunits are phylogenetically distant representatives of the DsrM-2 group, which includes proteins that belong to Dsr complexes that include all five subunits: M, K, J, O, and P. Cartoon and surface representations are shown for *Desulfobacca acetoxidans* **(A, E)**, *Desulfomonile tiedjei* **(B, F)**, *Desulfarculus baarsii* **(C, G)**, *Archaeoglobus sulfaticallidus* **(D, H)**. All four predicted DsrM homologs exhibit occluded quinol-oxidizing sites, highlighted by dashed circles, and also display elongated hydrophobic cavities (dashed ellipses) at positions equivalent to those in *A. fulgidus* DsrM-2, where a co-purified MK-7 is bound. Among the five output models generated per AF3 prediction, pairwise RMSDs range from 0.138 Å to 2.499 Å, and pTM scores range from 0.85 to 0.88, indicating confident, high-quality predictions. The first-ranked model from each prediction is shown. DsrM cartoon models are shown in shades of orange, from light in (A) to dark in (D). Hydrophobicity surfaces in (E-H) were generated as described in fig. S12. See methods for further details. Note that the DsrM-2 proteins shown here fall within the DsrM-2a subclass defined later in the Supplementary Text.

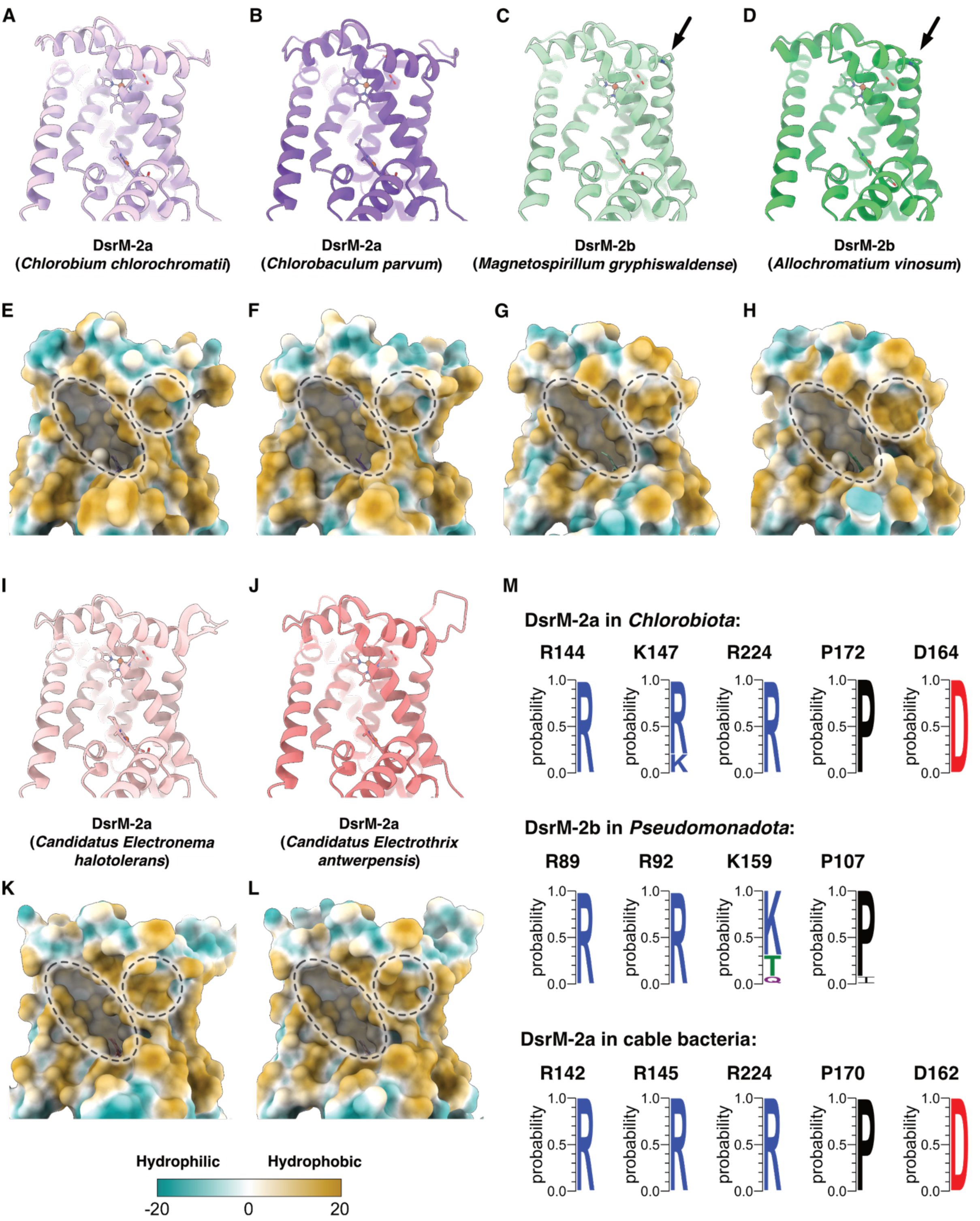


**Fig. S18. AF3-predicted structures reveal occluded quinol-oxidizing sites in DsrM from SOB and cable bacteria.** AF3 was used to predict the structures of the DsrMKJOP complexes from two sulfur-oxidizing *Chlorobiota* and two sulfur-oxidizing *Pseudomonadota*, as well as the DsrMKJ complexes from two cable bacteria species. Cartoon and surface representations are shown for *Chlorobium chlorochromatii* **(A, E)**, *Chlorobaculum parvum* **(B, F)**, *Magnetospirillum gryphiswaldense* **(C, G)**, *Allochromatium vinosum* **(D, H)**, *Candidatus Electronema halotolerans* **(I, K)**, and *Candidatus Electrothrix antwerpensis* **(J, L)**. All six predicted structures of DsrM homologs display occluded quinone-oxidizing sites (dashed circles) and elongated hydrophobic cavities (dashed ellipses) at positions corresponding to the MK-7-binding site in *A. fulgidus* DsrM. Among the five output models per prediction, pairwise RMSDs range from 0.113 Å to 3.253 Å, and pTM scores range from 0.84 to 0.88, indicating confident, high-quality predictions. The top-ranked model from each prediction is shown. DsrM cartoon representations are colored in distinct shades of purple, green, and red for *Chlorobiota*, *Pseudomonadota*, and cable bacteria, respectively. Hydrophobic surface representations (E-H, K-L) were generated as described in fig. S12. The proline residue in *Pseudomonadota* DsrM that introduces a kink in TMH3 is indicated by black arrows (C, D). See Methods for details. **(M)** Sequence conservation analysis reveals that Arg118, Arg121, and Arg198 in *A. fulgidus* DsrM are conserved as positively charged residues across SOB and cable bacteria. A conserved proline is observed in the inserted loop of *Chlorobiota* and cable bacteria DsrM, whereas DsrM of *Pseudomonadota* conserves a proline that introduces a transmembrane helix kink. Glu138 of *A. fulgidus* DsrM is conserved as an aspartate in *Chlorobiota* and cable bacteria but is absent in *Pseudomonadota*, together with the associated short helical segment. Residue numbering in the WebLogos follows that of *Chlorobium chlorochromatii* DsrM for *Chlorobiota*, *Allochromatium vinosum* DsrM for *Pseudomonadota*, and *Candidatus Electrothrix antwerpensis* DsrM for cable bacteria. The consensus residues in (M) are colored according to the chemical property of the residues.

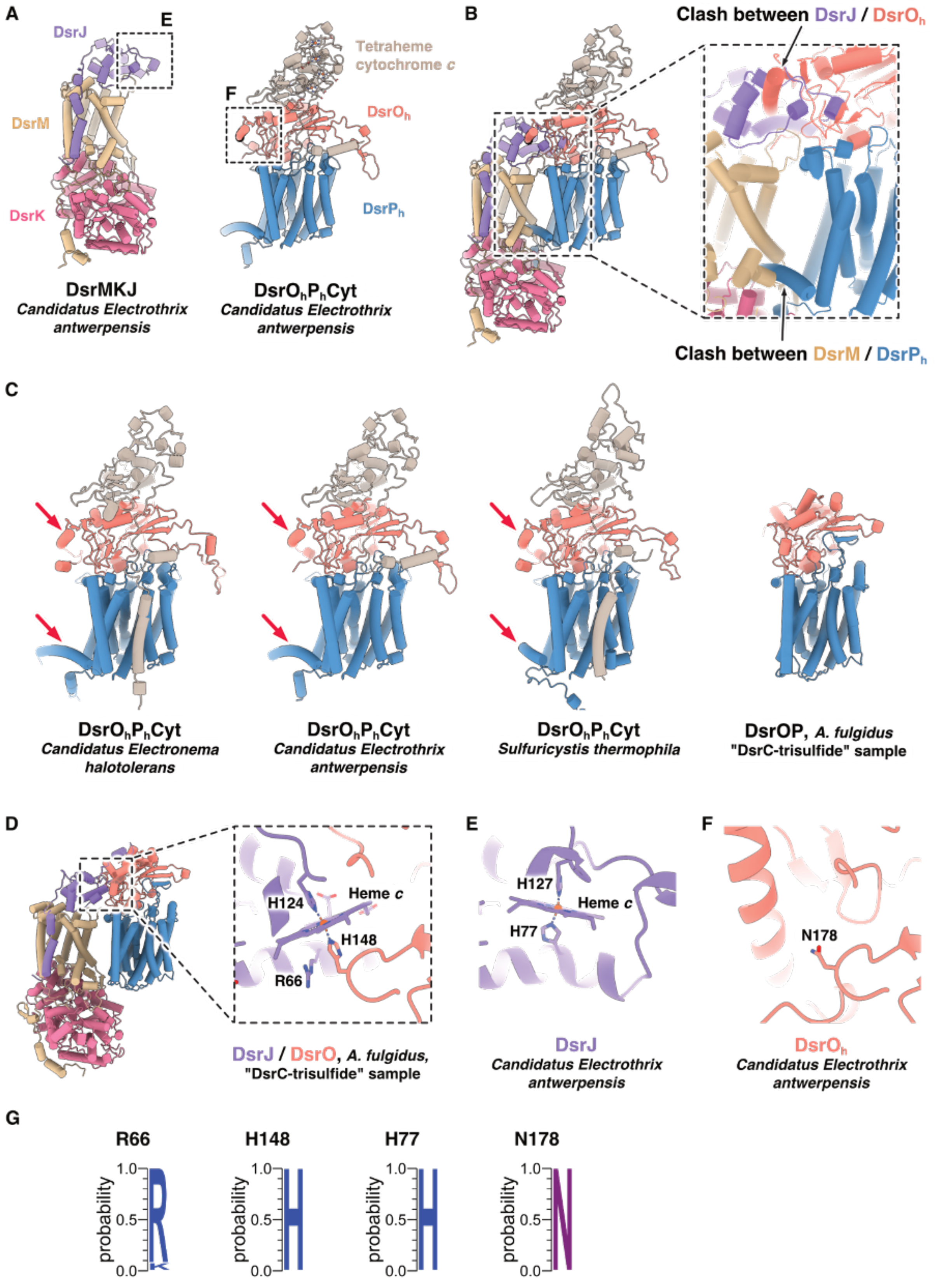


**Fig. S19. DsrMKJ and DsrO$_h$P$_h$Cyt likely form separate complexes in cable bacteria. (A)** AF3-predicted structures of DsrMKJ complex and DsrOPCyt complex from *Candidatus Electrothrix antwerpensis*. **(B)** Superposition of predicted DsrMKJ and DsrO$_h$P$_h$Cyt subcomplexes from *Candidatus Electrothrix antwerpensis* onto DsrM and DsrP subunits within our structure of DsrMKJOP from *A. fulgidus* reveals multiple steric clashes (black arrows). For clarity, *A. fulgidus* DsrMKJOP complex is not shown. **(C)** AF3-predicted structures of $DsrO_hP_hCyt$ from cable bacteria compared with homologous complexes from *Gamma-* and *Betaproteobacteria*, which lack DsrMKJ, represented here by *Sulfuricystis thermophila*, and DsrOP from *A. fulgidus*. Additional alpha helices in DsrO and DsrP (red arrows) would sterically interfere with DsrJ and DsrM, as illustrated in (B). **(D)** In *A. fulgidus*, a *c*-type heme at the DsrO-DsrJ interface is jointly coordinated by $His124^{DsrJ}$ and $His148^{DsrO}$. **(E-F)** In cable bacteria, the equivalent heme *c* is instead coordinated by two histidine residues from DsrJ, while $His148^{DsrO}$ is replaced by asparagine, precluding participation of DsrO in heme coordination. The histidine ligand $His77^{DsrJ}$ in cable bacteria corresponds to $Arg66^{DsrJ}$ in *A. fulgidus*. **(G)** Sequence conservation analysis indicates that $Arg66^{DsrJ}$ and $His148^{DsrO}$ in *A. fulgidus* are conserved among DsrM-2a-contaning SRM as well as sulfur-oxidizing *Chlorobiota* and *Pseudomonadota*, consistent with the presence of a DsrJ-DsrO interface in those lineages. In contrast, $His77^{DsrJ}$ is conserved exclusively among cable bacteria, while $Asn178^{DsrO}$ is conserved among cable bacteria and closely related *Gamma-* and *Betaproteobacteria*, lineages in which a DsrJ-DsrO interface is likely absent. Together, these observations support an evolutionary rewiring in which DsrMKJ and $DsrO_hP_hCyt$ function as separate membrane complexes in cable bacteria. The consensus residues in (G) are colored according to chemical properties. Among the five output models per prediction of $DsrO_hP_hCyt$, pairwise RMSDs range from 0.194 Å to 0.761 Å, and pTM scores range from 0.80 to 0.82. The top-ranked model from each prediction is shown.

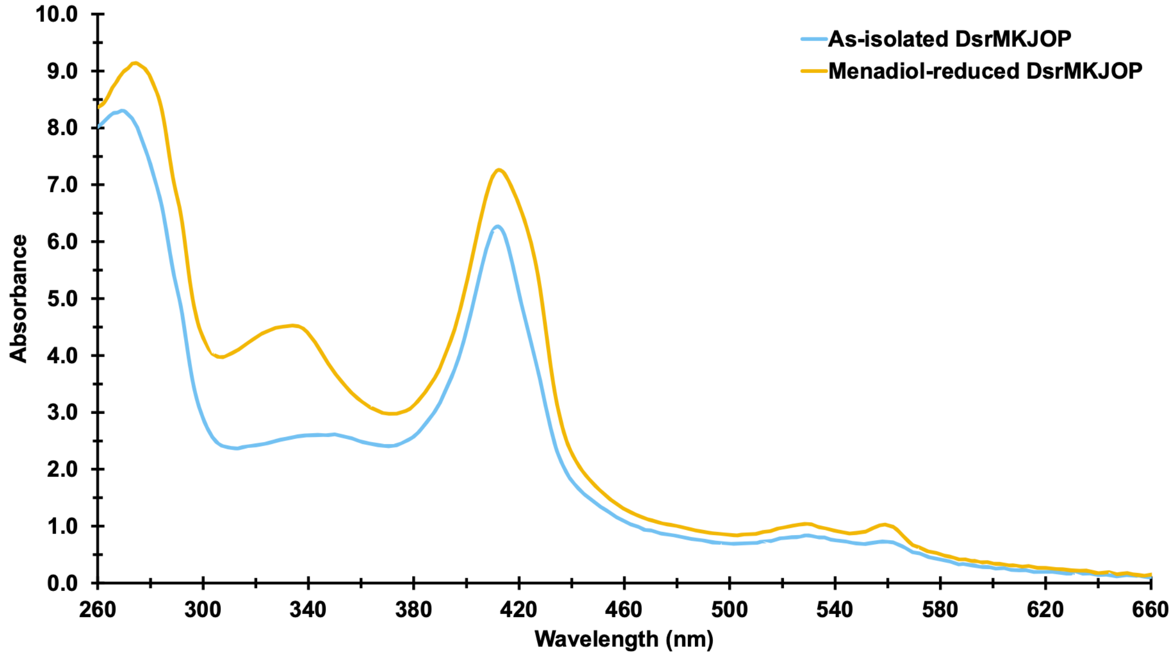


**Fig. S20. UV–Vis spectroscopy confirms DsrMKJOP reduction by menadiol.** UV–Vis spectra of "as-isolated" (blue) and menadiol-reduced (orange) DsrMKJOP are shown. The observed absorption maxima at 412 nm (Soret peak), 529 nm (β band), and 557 nm (α band) are characteristic of heme-containing proteins. Upon reduction, the α and β bands increase in intensity, and the Soret peak shifts toward 420 nm. The menadiol-reduced sample shows partial reduction of hemes, as previously reported [5].

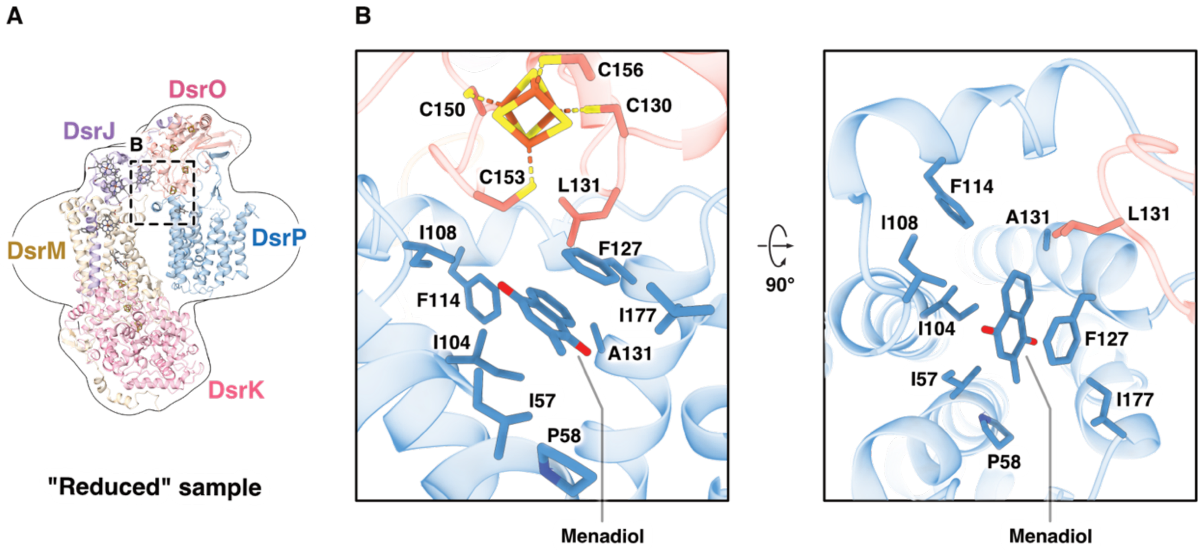


**Fig. S21. Menadiol is surrounded by a hydrophobic environment in DsrP. (A)** An overview of the pentameric DsrMKJOP from the "reduced" sample, shown using the same color scheme as in Fig. 1. The cryo-EM map of the "reduced" DsrMKJOP complex was Gaussian-filtered to 4 Å and is displayed as a transparent surface contoured at 8.1 σ. **(B)** Two close-up views of the menadiol-binding pocket, highlighting hydrophobic residues from DsrO and DsrP that surround the bound menadiol. One [4Fe-4S] cluster from DsrO, positioned within an efficient electron-transfer distance, is also shown along with its coordinating cysteine residues. All visualized cofactors, residues, and the menadiol are depicted as sticks and colored as follows: oxygen - red; nitrogen - blue; sulfur - yellow; iron – orange; and carbon - gray in (A) and steel blue in (B).

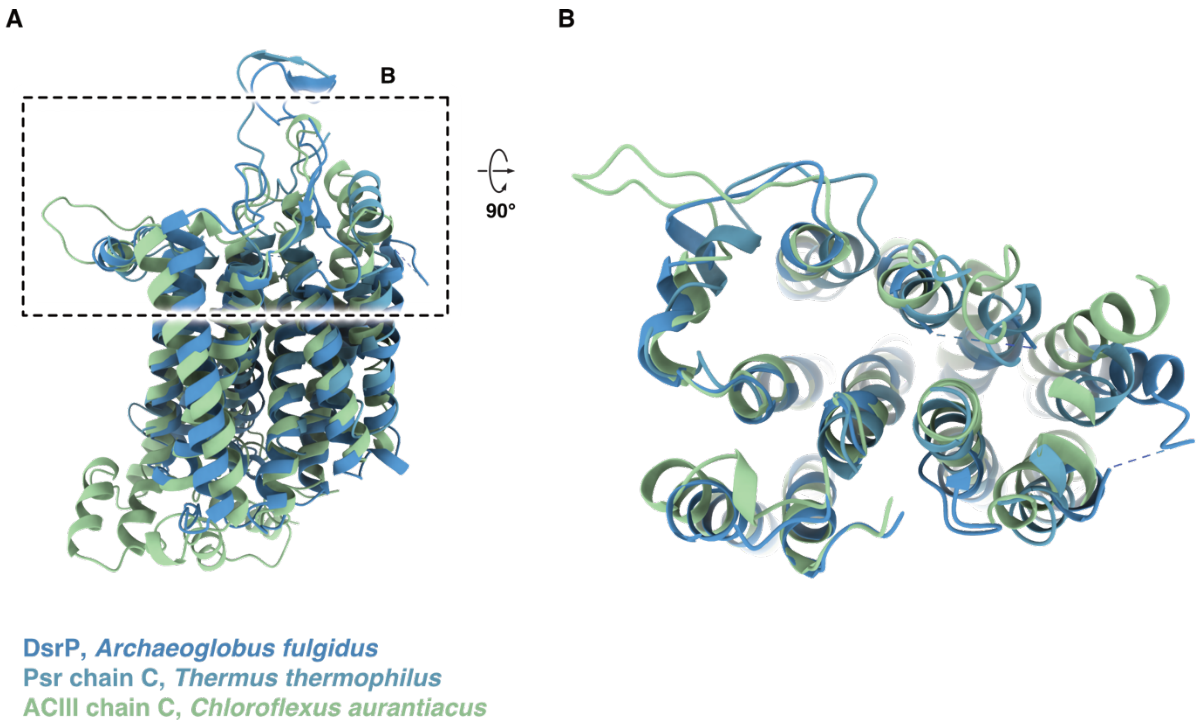


**Fig. S22. DsrP exhibits a structural fold similar to that of other NrfD family members.** The structure of *A. fulgidus* DsrP from the "reduced" sample is aligned with representative structures of NrfD family members: polysulfide reductase (Psr; *Thermus thermophilus*, PDB ID 2VPW [6], chain C), and alternative complex III (ACIII) from *Chloroflexus aurantiacus* (PDB ID 8X2J [7], chain C). Aligned structures are colored according to the legend and shown from two different viewing angles **(A, B)**.

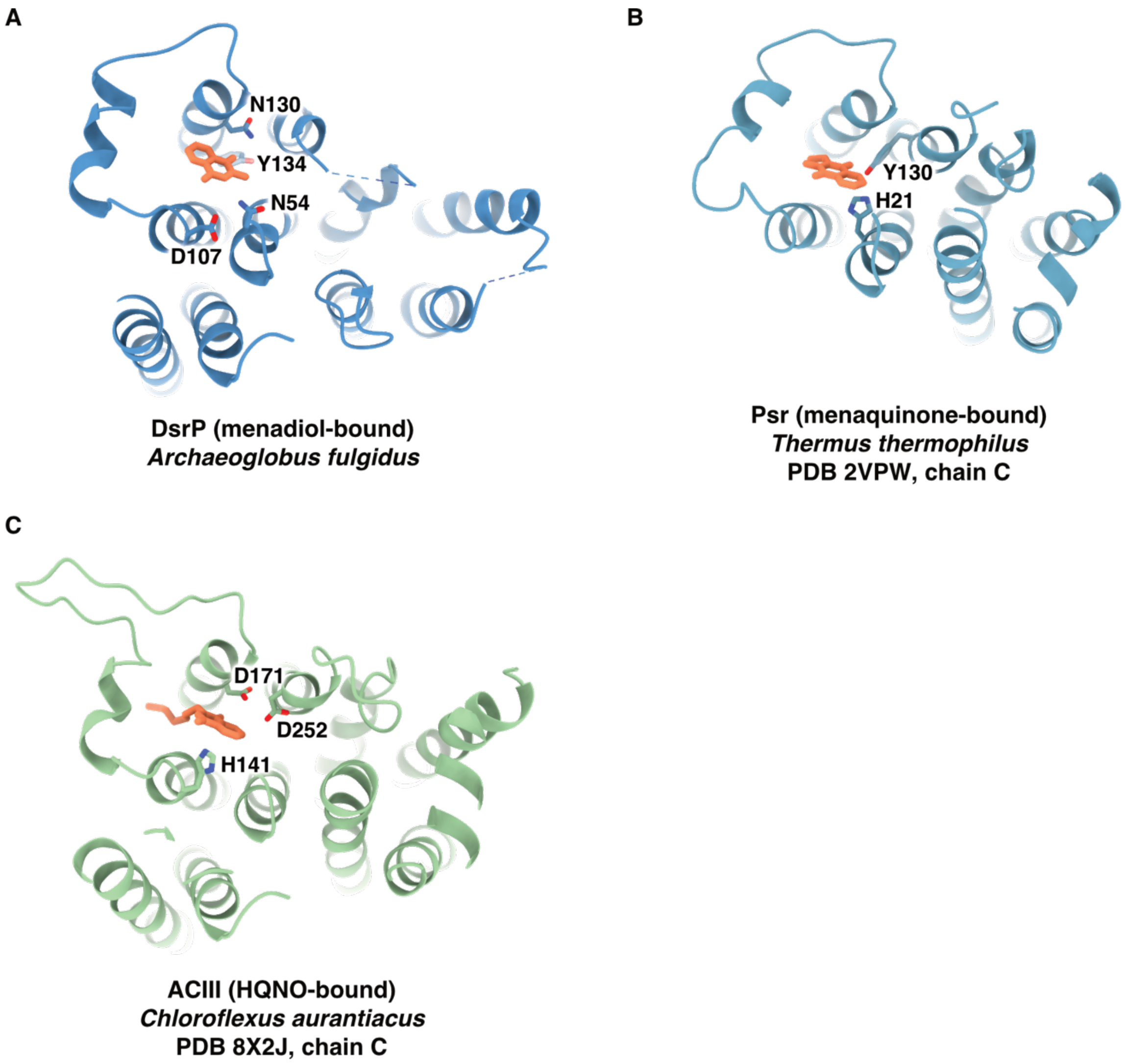


**Fig. S23. Quinol-binding pocket in DsrP aligns with those of other NrfD family members.** Structures shown in fig. S22 are displayed here individually in separate panels. Residues likely involved in quinol binding are depicted as sticks and colored according to atom type: oxygen - red; nitrogen - blue; and carbon matching the color of each corresponding protein chain. Menadiol, menaquinone, and 2-heptyl-4-hydroxyquinoline-N-oxide (HQNO) are shown as orange sticks.

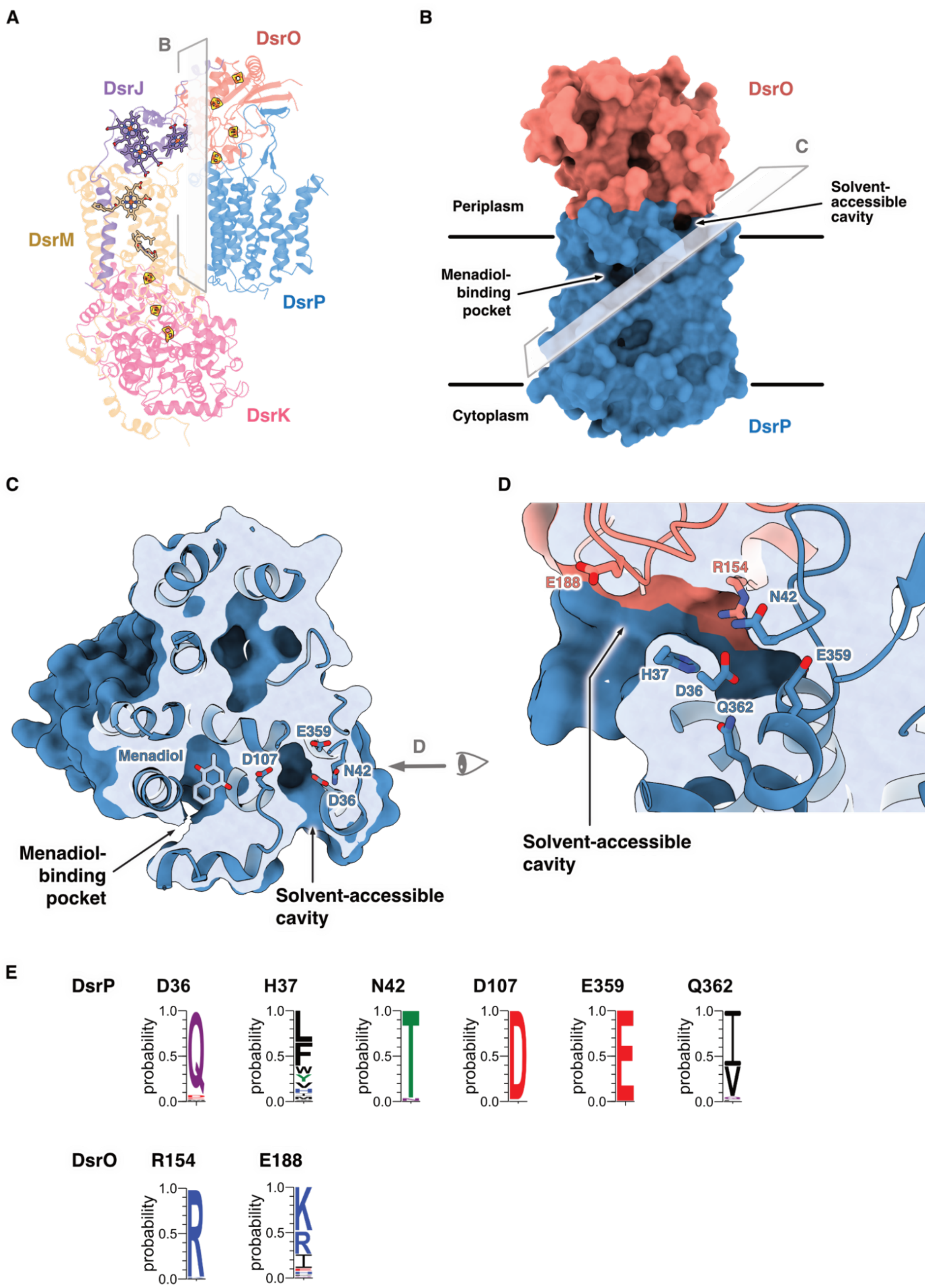


**Fig. S24. A solvent-accessible cavity at the DsrO:DsrP interface supports proton transfer to the periplasm. (A)** An overview of the pentameric DsrMKJOP complex from the "reduced" sample, shown in cartoon representation. Redox cofactors are displayed as sticks. **(B)** Surface view of the DsrOP subcomplex highlighting the spatial relationship between the menadiol-binding pocket and the solvent-accessible cavity. The membrane separating the periplasm from the cytoplasm is indicated schematically by two black horizontal lines. **(C-D)** Two sliced views of DsrOP showing the connection between the menadiol-binding pocket and the solvent-accessible cavity via Asp107$^{DsrP}$. Polar residues lining the cavity, potentially involved in proton transfer, are shown as sticks. **(E)** Sequence conservation analysis indicates that the physicochemical properties (*e.g.*, hydrophilicity) of most residues lining the cavity are conserved, particularly the Asp107$^{DsrP}$, Glu359$^{DsrP}$, and Arg154$^{DsrO}$. The consensus residues in (E) are colored according to the chemical property of the residues. The color scheme in (A-D) matches that in Fig. 1. In panels (B-D), solvent-excluded surfaces of the DsrOP subcomplex were generated in ChimeraX, with surface colors corresponding to each model chain. All visualized cofactors, residues, and the menadiol are depicted as sticks and colored as follows: oxygen - red; nitrogen - blue; sulfur - yellow; iron - orange; and carbon matching the color of each respective protein chain.

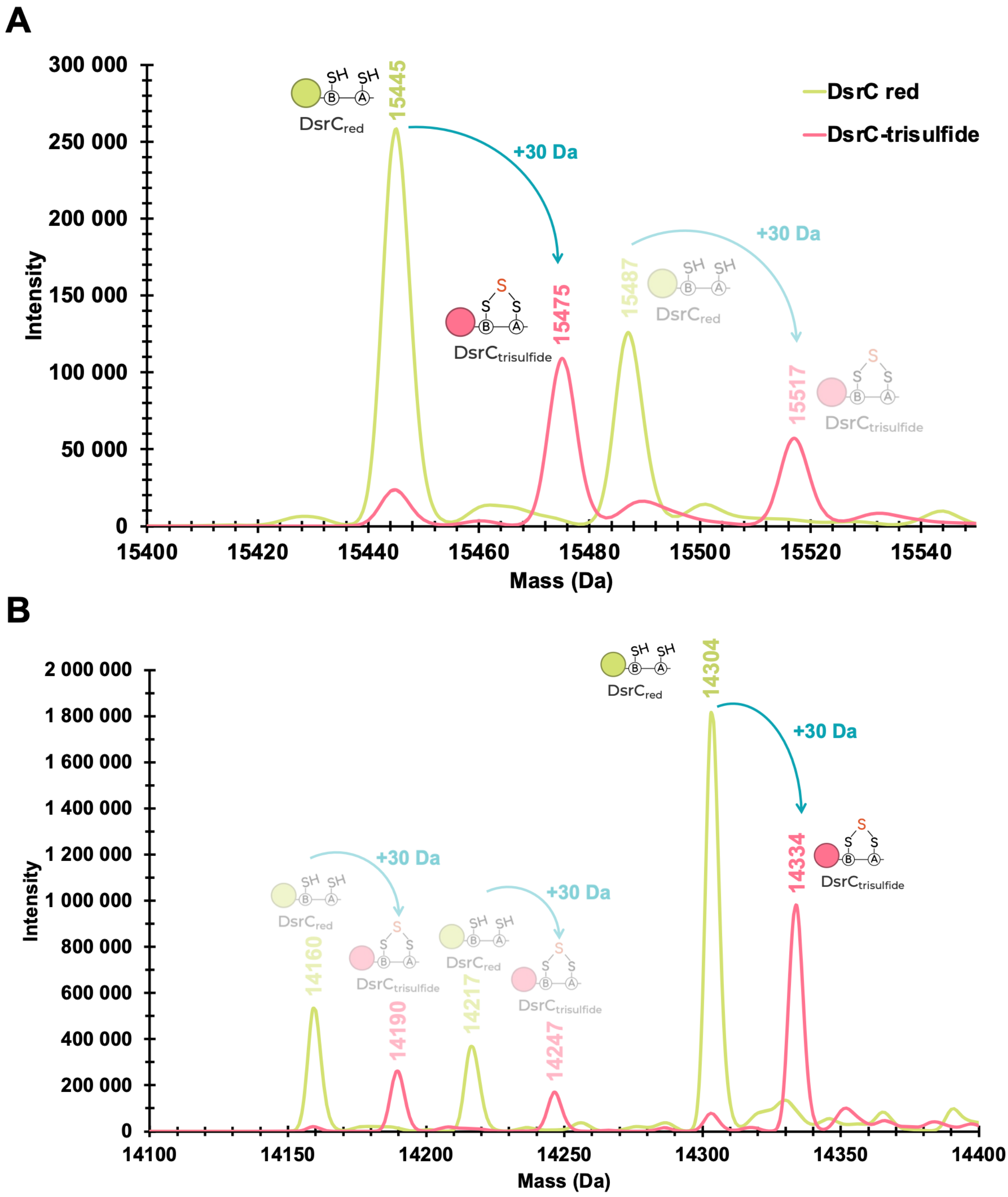


**Fig. S25. Intact MS analysis confirms DsrC-trisulfide formation.** Two similar but independent experiments are shown (**A** and **B**). Upon reaction of reduced DsrC (green) with DsrAB and sulfite, the mass of DsrC shifts by +30 Da, corresponding to the DsrC-trisulfide (pink). In **(B),** the mass of DsrC species is lower due to partial proteolysis of the N-terminal His-tag before the enzymatic reaction. This was prevented in **(A)** by the presence of protease inhibitors. Minor peaks from additional DsrC species are shown with a transparent filter.

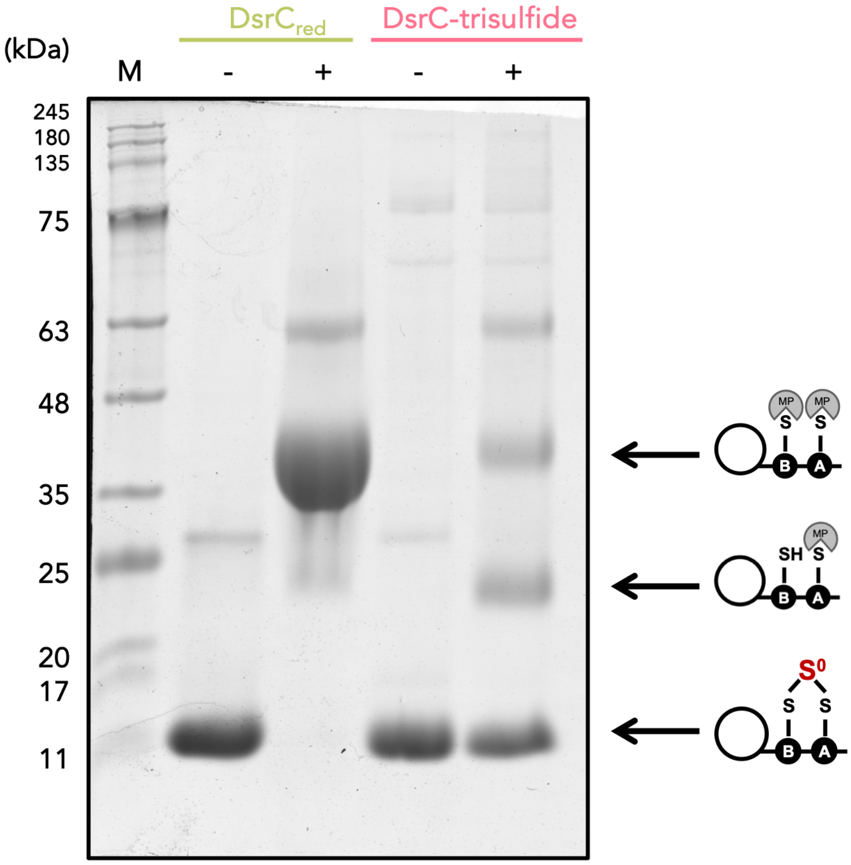


**Fig. S26. MalPEG gel-shift assay confirms DsrC-trisulfide formation.** Tricine-SDS-PAGE of DsrC (10 µg), in the absence (-) or presence (+) of 1 mM MalPEG. MalPEG-labeled DsrC in its reduced state (green) shows a shift of ~10 kDa per reduced cysteine. After reaction with DsrAB and sulfite, the resulting DsrC-trisulfide (pink) no longer reacts with MalPEG. The remaining bands observed in this lane correspond to reduced DsrC that did not react with DsrAB. Text labels for different DsrC species are colored consistently with fig. S25.

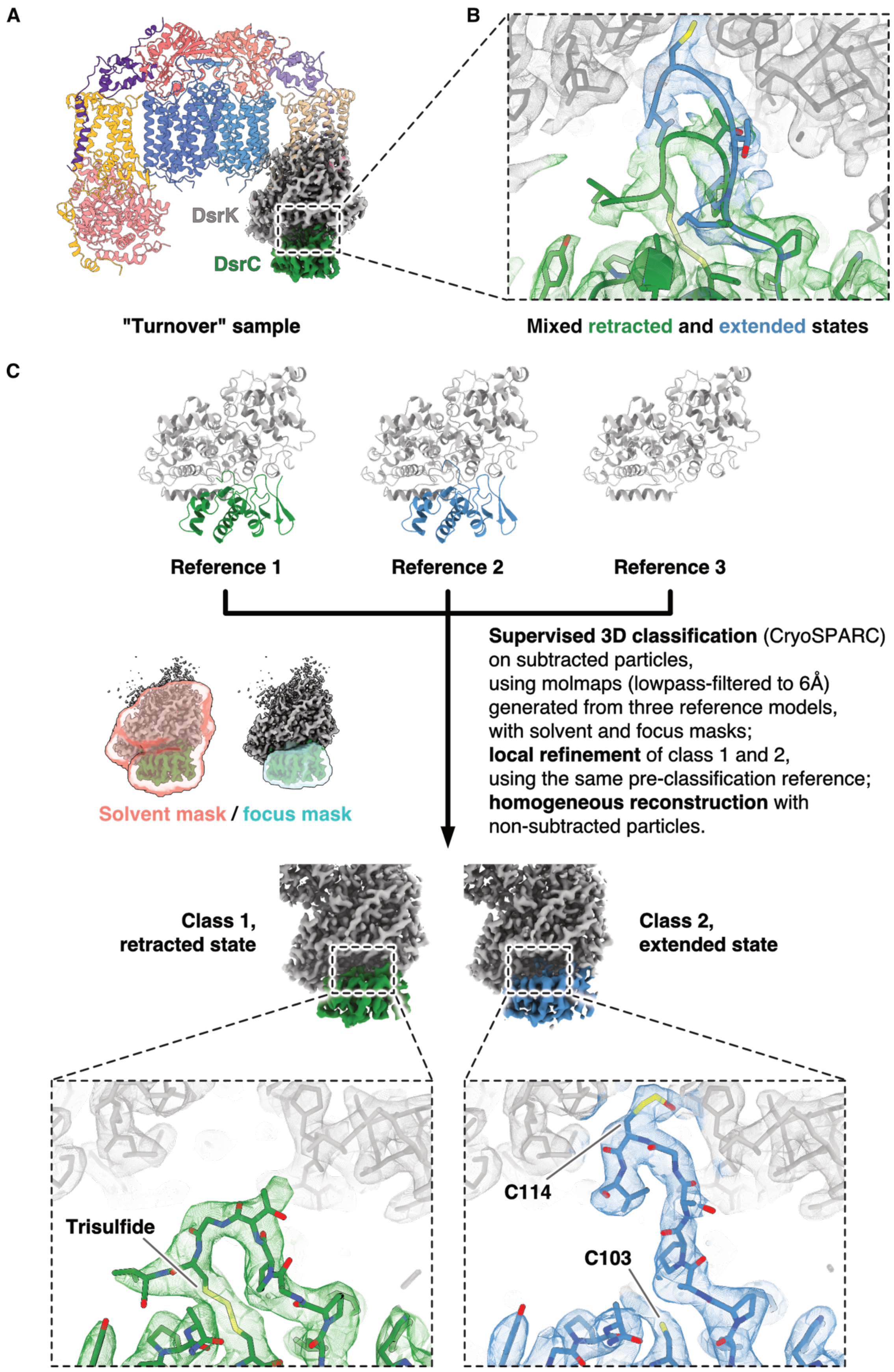


**Fig. S27. Separation of retracted and extended DsrC states by supervised 3D classification. (A)** An overview of the DsrMKJOP complex from the "turnover" sample, shown in cartoon representation using the same color scheme as in Fig. 1. The locally refined map, focused on DsrK and DsrC and obtained just prior to supervised classification, is superimposed onto the structural model. **(B)** This locally refined map, contoured at 5.25 σ, reveals that the C-terminal arm of DsrC adopts both retracted and extended conformations. **(C)** Following supervised 3D classification and local refinement, particles corresponding to these two conformations were separated, as shown in the bottom panels of (C), with both maps contoured at 8 σ. Class 3, derived from reference model 3 in which DsrC is absent, was not subjected to further refinement. Models and corresponding map densities for DsrK are displayed in gray. DsrC in the retracted state is colored green, while DsrC in the extended state is colored steel blue. The solvent mask and focus mask used during supervised classification are depicted as salmon and cyan surfaces, respectively.

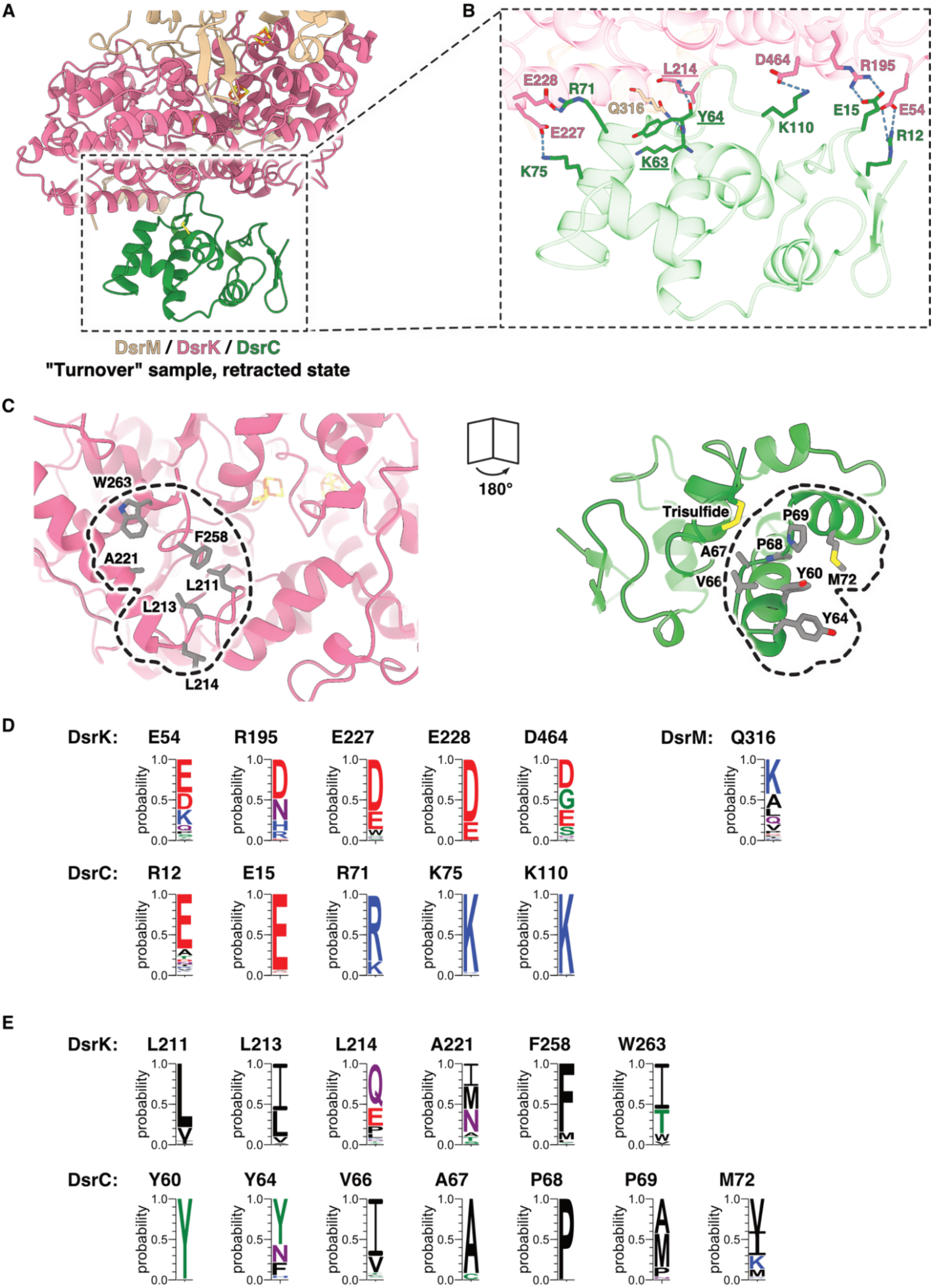


**Fig. S28. DsrC docking is primarily mediated by electrostatic attraction and hydrophobic effect. (A)** An overview of the DsrMKJOP complex bound to DsrC in the retracted state. DsrC forms extensive interactions with DsrK and a minor contact with DsrM. **(B)** A close-up view of the interactions formed by DsrC, with blue dashed lines indicating primarily electrostatic attractions and a few hydrogen bonds. **(C)** Side-by-side view of the exposed interface, highlighting residues contributing to the hydrophobic contacts. The color scheme in (A-C) matches that used in Fig. 1. Underlined residues participate in interactions via main-chain atoms. Sequence conservation analysis shows that for most of the residues involved in electrostatic attractions and hydrogen bonding **(D)**, as well as hydrophobic contacts **(E)**, the physicochemical properties (*e.g.,* hydrophobicity) are conserved. The consensus residues in (D) and (E) are colored according to the chemical property of the residues.

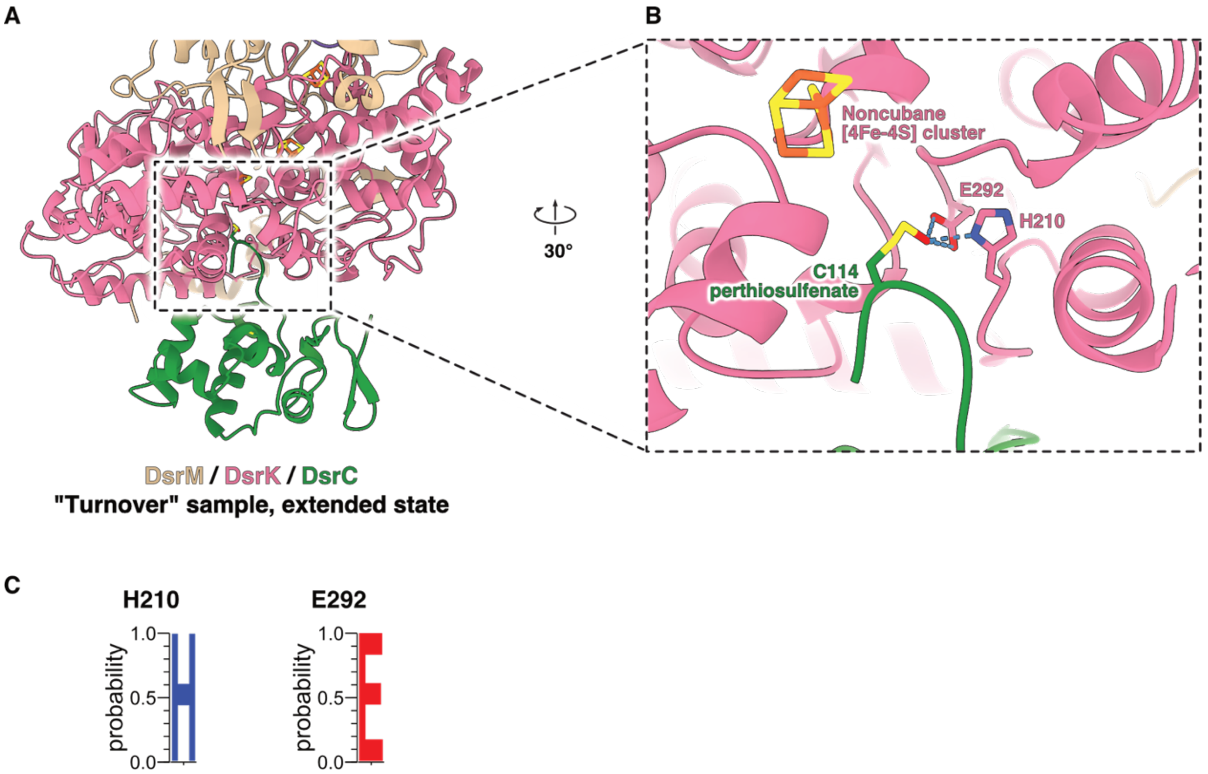


**Fig. S29. Cys114-perthiosulfenate in DsrC forms hydrogen bonds with DsrK. (A)** An overview of the DsrMKJOP complex bound to DsrC in the extended state. **(B)** A close-up view reveals hydrogen bonds between the Cys114-perthiosulfenate and His210$^{DsrK}$ and Glu292$^{DsrK}$, indicated by blue dashed lines. The color scheme in (A-B) matches that used in Fig. 1. **(C)** Sequence conservation analysis shows that His210$^{DsrK}$ and Glu292$^{DsrK}$ are well conserved. The consensus residues in (C) are colored according to the chemical property of the residues.

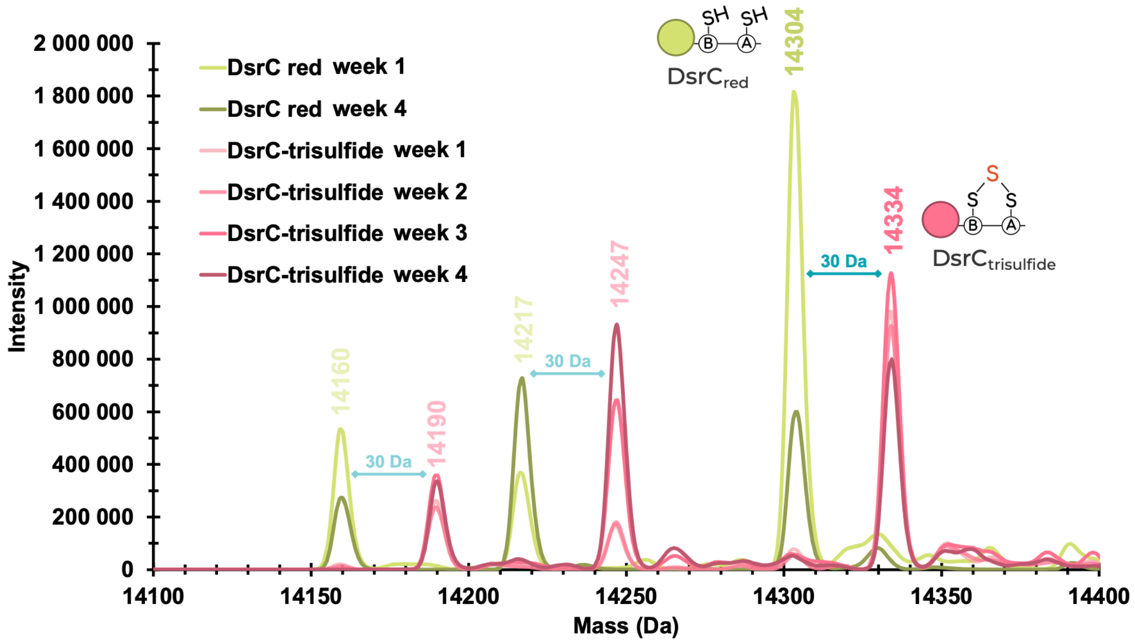


**Fig. S30. Intact MS analysis reveals DsrC-trisulfide stability.** Reduced DsrC (green) and DsrC-trisulfide (pink) were incubated at 4 °C under anaerobic conditions for four weeks. During this period, the relative intensities of the major peaks at 14304 Da ($DsrC_{red}$) and 14334 Da (DsrC-trisulfide) decreased, due to proteolytic degradation at the N-terminus, while the mass difference of +30 Da between both was maintained throughout the experiment, demonstrating the stability of the trisulfide bond under anaerobic conditions. Minor peaks from additional DsrC species are shown with a transparent filter.

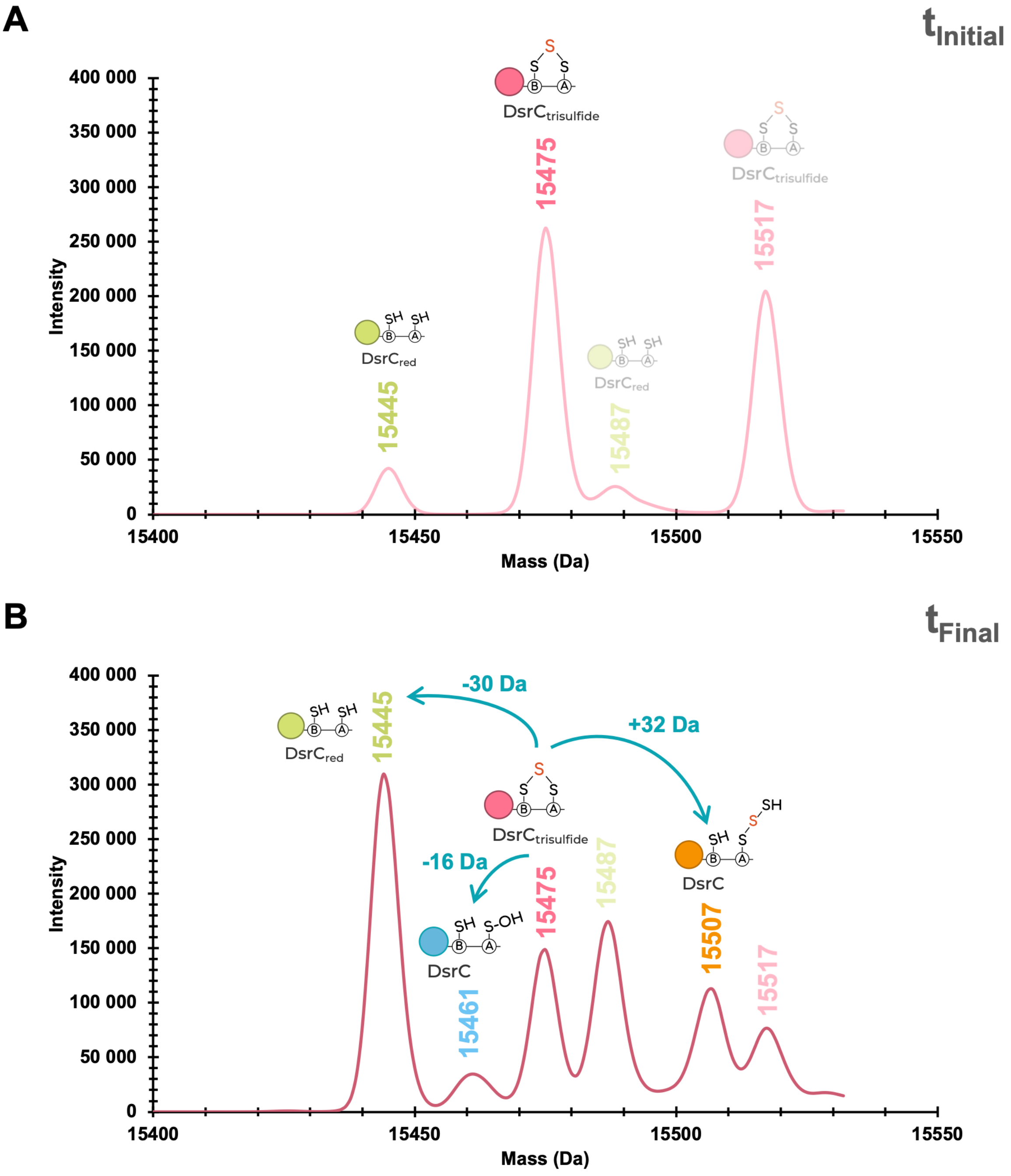


**Fig. S31. Intact MS analysis of DsrC-trisulfide reaction with catalytic DsrMKJOP and excess menadiol.** DsrC-trisulfide (10 µM) before **(A)** and after **(B)** reacting with a catalytic amount of DsrMKJOP (100 nM) and excess menadiol (500 µM) for 30 min at 60 °C. At the end of the reaction **(B)**, the intensity of DsrC-trisulfide peaks (15475 Da and 15517 Da, pink) decreases, while the peaks corresponding to reduced DsrC (15445 Da and 15487 Da, green) increase. Additional peaks at 15461 Da (blue) and 15507 Da (orange) are detected, which match the expected mass of a sulfenate state of one cysteine in reduced DsrC and a DsrC species with an additional sulfur atom versus to the trisulfide form, respectively. Minor peaks from additional DsrC species are shown with a transparent filter.

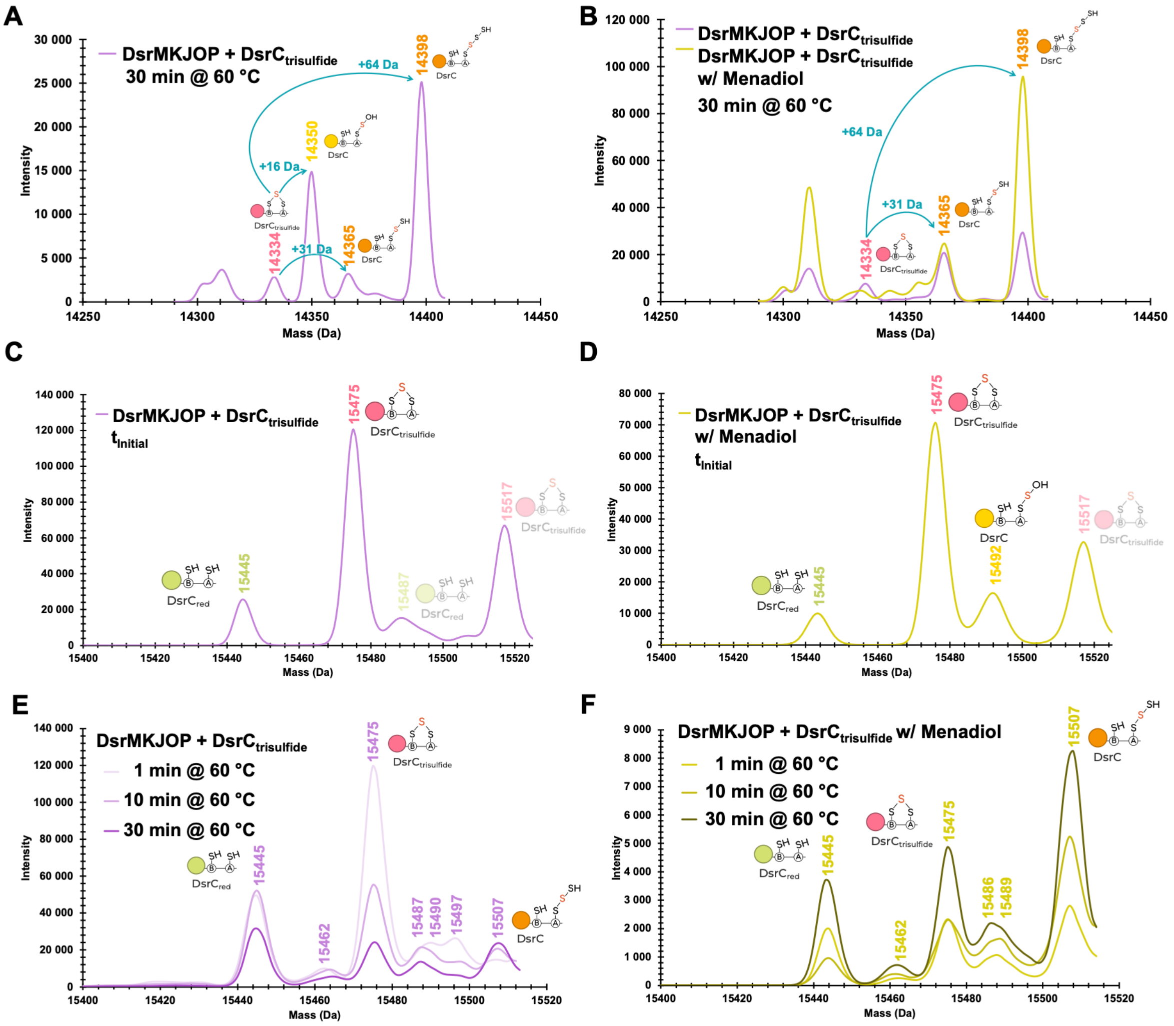


**G**

| | $DsrC_{red}$ | $DsrC_{trisulfide}$ | DsrC |
|---|---|---|---|
| 1 min @ 60 °C | 0.20 | 0.44 | 0.03 |
| 10 min @ 60 °C | 0.29 | 0.30 | 0.08 |
| 30 min @ 60 °C | 0.29 | 0.20 | 0.20 |
| | Relative abundance | | |

**H**

| | $DsrC_{red}$ | $DsrC_{trisulfide}$ | DsrC |
|---|---|---|---|
| 1 min @ 60 °C | 0.28 | 0.22 | 0.22 |
| 10 min @ 60 °C | 0.18 | 0.19 | 0.27 |
| 30 min @ 60 °C | 0.16 | 0.19 | 0.32 |
| | Relative abundance | | |

**Fig. S32. Intact MS analysis of the stoichiometric reaction of DsrC-trisulfide with DsrMKJOP.** DsrC-trisulfide (10 µM) was incubated with equimolar DsrMKJOP complex in seven separate experiments, either in the absence (purple traces, left) or presence (green traces, right) of excess menadiol. Major peaks are annotated and color-coded as follows: reduced DsrC (green), DsrC-trisulfide (pink), perthiosulfenate-DsrC (Cys-S–SOH, yellow), and persulfidated DsrC (Cys-S–S–SH, orange). **(A)** and **(B)** correspond to experiments using a DsrC sample that suffered N-terminal proteolytic cleavage (lower mass). After a 30-minute incubation at 60 °C in the absence of menadiol (A), a peak with +16 Da versus the DsrC-trisulfide was detected, consistent with a perthiosulfenate intermediate (yellow). This experiment was repeated in (B), where this intermediate was not observed, possibly due to its transient nature. In both experiments, incubation with the DsrMKJOP complex promoted the incorporation of additional sulfur atoms in DsrC (+32 Da, +64 Da, orange peaks). In the presence of menadiol (500 µM) (B), similar results were observed. In experiments **(C)** and **(D)**, the DsrC-trisulfide was mixed 1:1 with DsrMKJOP and frozen immediately in liquid $N_2$ before storage at -80 ºC and MS measurement. In the absence of menadiol (C), the DsrC-trisulfide remains intact, whereas in the presence of menadiol (5 mM), the +16 Da perthiosulfenate peak (yellow) was once again observed. In experiments **(E)** (no menadiol) and **(F)** (with menadiol), the reaction was monitored at different time points (1, 10, and 30 minutes). In the absence of menadiol (E), the DsrC-trisulfide is consumed over time, resulting in an increase of reduced DsrC and a DsrC species with an additional sulfur atom relative to the trisulfide form. A similar result is observed in the presence of menadiol (5 mM) (F), with the formation of the persulfidated DsrC being more prominent. The presence or absence of a perthiosulfenate species in this experiment could not be assessed due to the presence of broad peaks between 15480 and 15500 Da (already observed in the DsrC-trisulfide before reaction). The relative abundance of each DsrC species (area of each peak divided by the sum of the peak areas in each panel) in experiments (E) and (F) is shown in **(G)** and **(H)**, respectively.

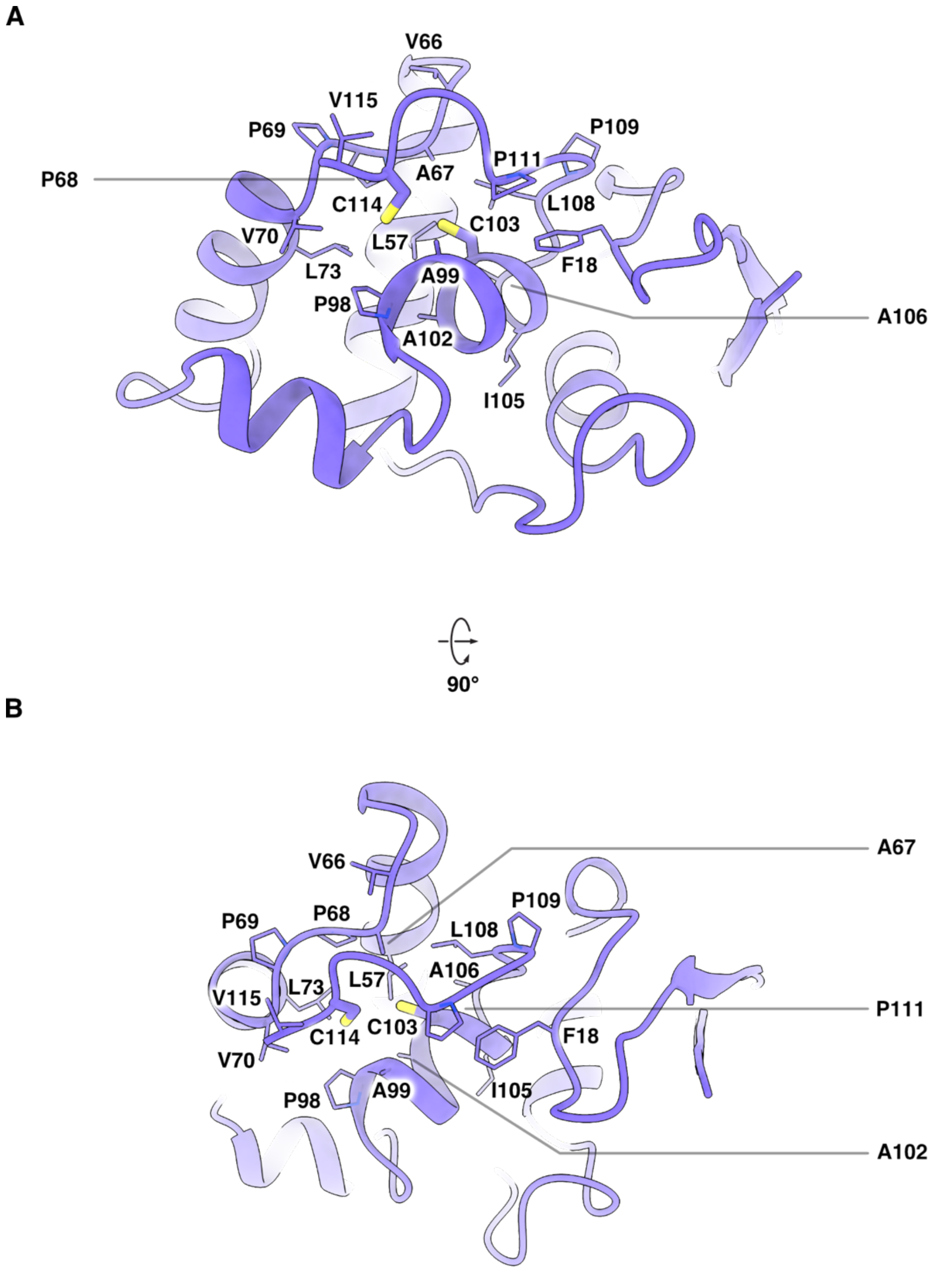


**Fig. S33. Cys103 and Cys114 are embedded in a hydrophobic environment in free DsrC. (A-B)** Two different view angles show that Cys103$^{DsrC}$ and Cys114$^{DsrC}$ are surrounded by hydrophobic residues within free DsrC (PDB ID 1SAU [8]; purple cartoon). Highlighted residues are shown in stick representation and colored as follows: carbon - purple; nitrogen - blue; and sulfur - yellow.

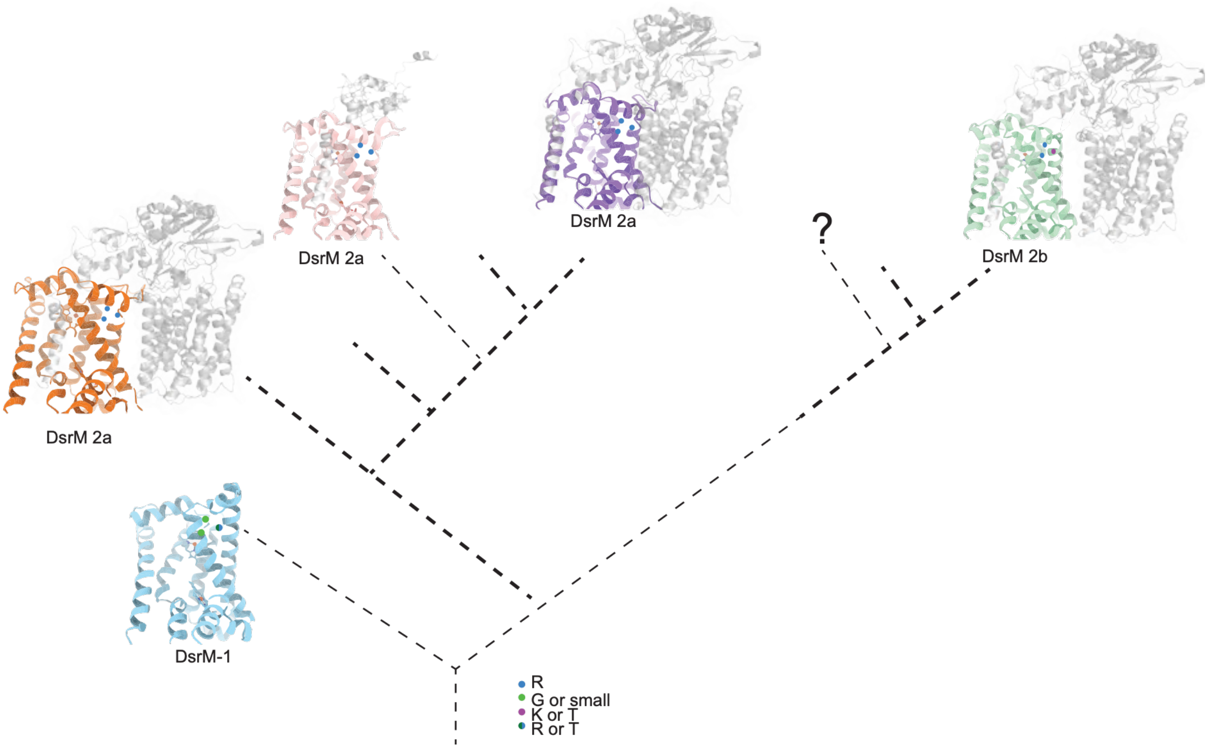


**Fig. S34. Schematic representation of the relationships between the different DsrM types.** DsrM structures are shown in light blue (DsrM-1), orange (DsrM2-a from sulfate reducers) pink (DsrM2-a from cable bacteria), purple (DsrM-2a from Chlorobi) and green (DsrM-2b from Pseudomonadota), whereas DsrJ, DsrO and DsrP are shown in grey. For simplicity, DsrK structures are omitted. Thin lines represent DsrMK or DsrMKJ complexes while thicker lines indicate DsrMKJOP complexes. The PCP site is shown with amino acids coloured according to their physicochemical properties. Empty branches represent other DsrM sequences. The PCP site appears available for quinone-binding in DsrM-1. Sequence changes in DsrM-2a complexes generate an occluded PCP site and facilitate interaction of the DsrJ, DsrO and DsrP subunits. The DsrMKJ from cable bacteria appears to have evolved from a DsrMK-2a complex that lost the J and O subunits, while in Chlorobi, a DsrM-2a complex appears to have been adapted to work in the oxidative direction (reverse Dsr pathway). An independent evolutionary event appears to have given rise to the DsrM-2b-type complexes, which have different amino acids motifs in the PCP site, as well as a different DsrM-DsrJ interface. These complexes are present in sulfur oxidizers. The question mark denotes putative DsrMK complexes basal to oxidative DsrM-2b as hypothesized in [1].

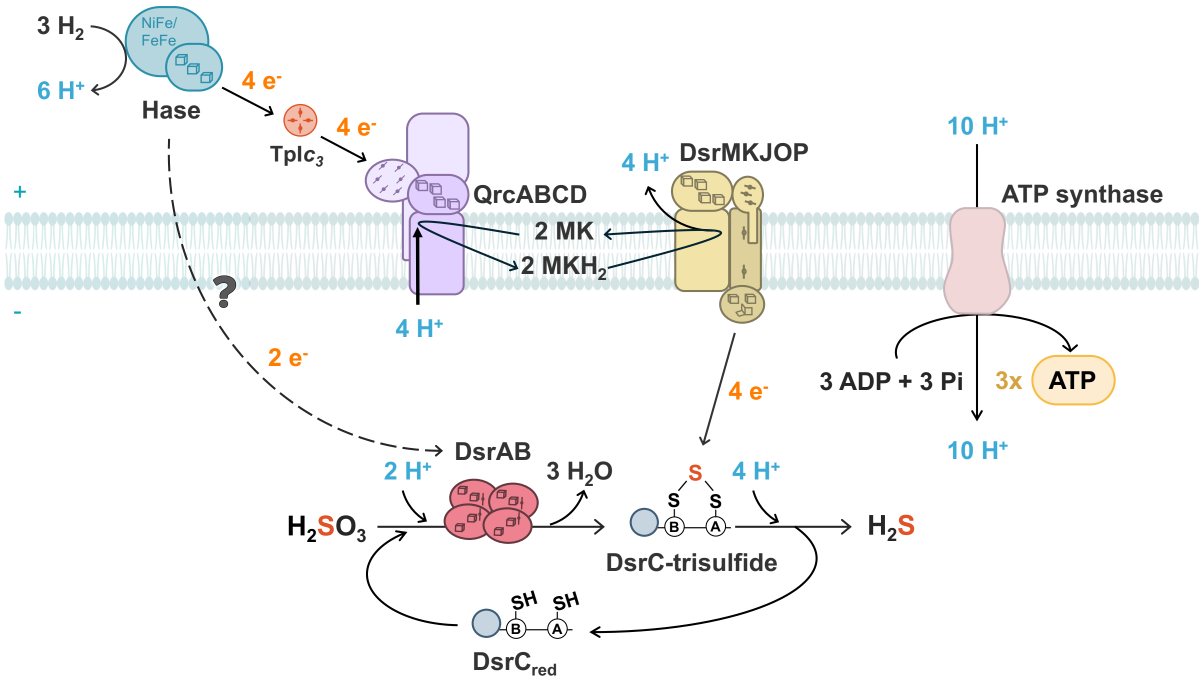


**Fig. S35. Bioenergetic model of *N. vulgaris* during $H_2$-dependent sulfite respiration.** Sulfite reduction proceeds in two steps: a two-electron reduction of sulfite to the DsrC-trisulfide intermediate, followed by a four-electron reduction of DsrC-trisulfide to sulfide. In the first step, electrons are assumed to derive from periplasmic $H_2$ oxidation, releasing two protons, and are delivered to DsrAB in the cytosol by an unknown electron carrier (dashed lines), where two protons are consumed during formation of the DsrC-trisulfide. In the second step, oxidation of two $H_2$ molecules by periplasmic hydrogenases (Hase) releases four protons and feeds four electrons into the menaquinone pool via Type I cytochrome $c_3$ (TpI$c_3$) and the QrcABCD complex, coupled to uptake of four protons from the cytoplasm [9]. Subsequent menaquinol oxidation at DsrMKJOP releases four additional protons into the periplasm, while DsrC-trisulfide reduction consumes four protons from the cytoplasm. Thus, transfer of these four electrons contributes eight protons to the pmf. Together, the two steps account for a translocation of ten protons across the membrane, sufficient to drive synthesis of approximately three ATP molecules, assuming an ATP synthase c-ring stoichiometry of 10.

**Table S1. EM statistics**

| | "As-isolated" sample | "DsrC-trisulfide" sample | "Reduced" sample | "Turnover" sample |
|---|---|---|---|---|
| **Data collection** | | | | |
| Electron microscope | Titan Krios G2 | Titan Krios G4 | Titan Krios G4 | Titan Krios G4 |
| Camera | K3 | Falcon 4 | Falcon 4 | Falcon 4i |
| Data collection software | EPU with AFIS | EPU with AFIS | EPU with AFIS | EPU with AFIS |
| Voltage | 300 kV | 300 kV | 300 kV | 300 kV |
| Nominal magnification | 105,000 x | 165,000 x | 165,000 x | 165,000 x |
| Calibrated pixel size | 0.831 Å | 0.73 Å | 0.73 Å | 0.73 Å |
| Total exposure | 60 $e^-/Å^2$ | 60 $e^-/Å^2$ | 60 $e^-/Å^2$ | 60 $e^-/Å^2$ |
| Number of frames | 61 / 61 fractions[*] | 1652 raw frames[†] | 1491 raw frames[†] | 1152 raw frames[†] |
| Defocus range | -0.8 to -2.5 μm | -0.8 to -2.5 μm | -0.8 to -2.5 μm | -0.8 to -2.5 μm |
| **Image processing** | | | | |
| Motion correction software | MotionCor2 | MotionCor2 (RELION implementation) | MotionCor2 (RELION implementation) | MotionCor2 (RELION implementation) |
| CTF estimation | CTFFIND 4.1.13 | CTFFIND 4.1.13 | CTFFIND 4.1.13 | CTFFIND 4.1.13 |
| Particle selection software | TOPAZ | TOPAZ | TOPAZ | TOPAZ |
| Exposures used | 3,688 / 11,086 (TIFF format) [‡] | 13,250 (EER format) | 9,148 (EER format) | 16,205 (EER format) |
| Classification software | RELION 3.1 / CryoSPARC 3.3 | RELION 4.0 / CryoSPARC 3.3 | RELION 4.0 / CryoSPARC 4.3 | RELION 4.0 / CryoSPARC 4.3 |
| Refinement software | CryoSPARC 3.3 | CryoSPARC 3.3 | CryoSPARC 4.3 | CryoSPARC 4.3 |
| **Model building** | | | | |
| Modeling software | Coot | Coot | Coot | Coot |
| Refinement software | *PHENIX* | *PHENIX* | *PHENIX* | *PHENIX* |

* Respective counts of fractions for the two datasets acquired for the “as-isolated” sample.
† Respective numbers of raw detector frames for “DsrC-trisulfide”, “reduced”, and “turnover” samples. These raw frames were grouped into 60 fractions during beam-induced motion correction.
‡ Respective counts of exposures for the two datasets acquired for the “as-isolated” sample.

### Table S2. Cryo-EM map identifiers and quality statistics

| EMDB ID | Description | Resolution in Å ($FSC_{0.143}$) | Sharpening B-factor | # of particles | Symm. |
|---|---|---|---|---|---|
| EMD-54329 | As-isolated DsrMKJOP, density modified map | 2.20 | n/a | 140,106 | $C_2$ |
| EMD-54330 | DsrMKJOP in complex with DsrC, locally filtered map (“DsrC-trisulfide” sample) | 1.97 | n/a | 65,708 | $C_2$ |
| EMD-54331 | DsrMKJOP in complex with DsrC, focused map (“turnover” sample, retracted state) | 2.55 | -38.3 | 40,856* | $C_1$ |
| EMD-54332 | DsrMKJOP in complex with DsrC, consensus map (“turnover” sample, retracted state) | 2.17 | -28.5 | 40,856* | $C_1$ |
| EMD-54333 | DsrMKJOP in complex with DsrC, composite map generated from EMD-54331 and EMD-54332 (“turnover” sample, retracted state) | Composite map | n/a | 40,856* | $C_1$ |
| EMD-54334 | DsrMKJOP in complex with DsrC, focused map (“turnover” sample, extended state) | 2.53 | -38.3 | 43,375* | $C_1$ |
| EMD-54335 | DsrMKJOP in complex with DsrC, consensus map (“turnover” sample, extended state) | 2.16 | -28.6 | 43,375* | $C_1$ |
| EMD-54336 | DsrMKJOP in complex with DsrC, composite map generated from EMD-54334 and EMD-54335 (“turnover” sample, extended state) | Composite map | n/a | 43,375* | $C_1$ |
| EMD-54337 | Menadiol-treated DsrMKJOP, locally filtered map (“reduced” sample) | 2.10 | n/a | 65,514 | $C_1$ |

*Particle counts after $C_2$ symmetry expansion.

**Table S3. Model identifiers and quality statistics***

| DsrMKJOP ("as-isolated" sample) | |
|---|---|
| PDB ID | 9RWH |
| EMDB ID | EMD-54329 |
| Residues built | 3280 |
| Ligands built | 12x SF4, 6x HEC, 4x HEO, 2x 9S8, 2x MQ7, 2x H2S |
| RMS bond lengths | 0.003 |
| RMS bond angles | 0.881 |
| Ramachandran outliers | 0.00% |
| Ramachandran favored | 97.91% |
| Rotamer outliers | 0.21% |
| Clash score | 3.61 |
| CaBLAM outliers | 0.86% |
| C-beta outliers | 0.00% |
| DsrMKJOP in complex with DsrC ("Dsr-trisulfide" sample) | |
| PDB ID | 9RWJ |
| EMDB ID | EMD-54330 |
| Residues built | 3508 |
| Ligands built | 12x SF4, 6x HEC, 4x HEO, 2x 9S8, 2x MQ7, 2x H2S |
| RMS bond lengths | 0.002 |
| RMS bond angles | 0.836 |
| Ramachandran outliers | 0.00% |
| Ramachandran favored | 98.39% |
| Rotamer outliers | 0.00% |
| Clash score | 1.70 |
| CaBLAM outliers | 0.58% |
| C-beta outliers | 0.00% |
| DsrMKJOP in complex with DsrC ("turnover" sample, retracted state) | |
| PDB ID | 9RWK |
| EMDB ID | EMD-54333 |
| Residues built | 3388 |
| Ligands built | 12x SF4, 6x HEC, 4x HEO, 2x 9S8, 2x MQ7, 2x H2S |
| RMS bond lengths | 0.003 |
| RMS bond angles | 0.873 |
| Ramachandran outliers | 0.00% |
| Ramachandran favored | 97.92% |
| Rotamer outliers | 0.52% |
| Clash score | 2.40 |
| CaBLAM outliers | 0.90% |
| C-beta outliers | 0.00% |

| | |
|---|---|
| DsrMKJOP in complex with DsrC ("turnover" sample, extended state) | |
| PDB ID | 9RWL |
| EMDB ID | EMD-54336 |
| Residues built | 3389 |
| Ligands built | 12x SF4, 6x HEC, 4x HEO, 2x 9S8, 2x MQ7, 2x H2S |
| RMS bond lengths | 0.003 |
| RMS bond angles | 0.875 |
| Ramachandran outliers | 0.00% |
| Ramachandran favored | 98.30% |
| Rotamer outliers | 0.41% |
| Clash score | 2.40 |
| CaBLAM outliers | 0.72% |
| C-beta outliers | 0.00% |
| Menadiol-treated DsrMKJOP ("reduced" sample) | |
| PDB ID | 9RWN |
| EMDB ID | EMD-54337 |
| Residues built | 1602 |
| Ligands built | 6x SF4, 3x HEC, 2x HEO, 1x 9S8, 1x MQ7, 1x H2S, 1x VK3 |
| RMS bond lengths | 0.004 |
| RMS bond angles | 0.918 |
| Ramachandran outliers | 0.00% |
| Ramachandran favored | 98.61% |
| Rotamer outliers | 0.07% |
| Clash score | 2.46 |
| CaBLAM outliers | 0.64% |
| C-beta outliers | 0.00% |

* Quality statistics were obtained from PHENIX real-space refinement.

**Table S4. List of plasmids and strains used in this study.**

| Plasmid | Relevant | Source or reference |
|---|---|---|
| **pMO719** | pCR8/GW/TOPO containing SRB replicon (pBG1); Spec® | Keller *et al.*, 2009 [10] |
| **pSC27** | *Desulfovibrio* shuttle vector; source of aph(3')-II; Km® | Rousset *et al.*, 1998 [11] |
| **pMO9075** | pCR8/GWTOPO with pBG1; Km® gene aph(3')-II promoter; multicloning site; Spec® | Keller *et al.*, 2011 [12] |
| **pMOIP03** | pMO9075 expressing *hysAB* with Strep-tag at C-terminal of *hysA*; Spec® | Marques *et al.*, 2017 [13] |
| **pMOIPAB01** | Δ*dsrJOP*; Km® | This work |
| **pMOIPAB02** | Δ*dsrJ*; Km® | This work |
| **pMO-*dsrJ*** | pMOIP03 with *hysAB* replaced by *dsrJ* from *N. vulgaris* with Strep-tag at C-terminal; Spec® | This work |
| **pMO-*dsrJ* C45A** | pMO-*dsrJ* with one point mutation Cys45Ala, Strep-tag at C-terminal; Spec® | This work |
| **pMO-*dsrJ* C45H** | pMO-*dsrJ* with one point mutation Cys45His, Strep-tag at C-terminal; Spec® | This work |
| **pMO-*dsrJ* C45S** | pMO-*dsrJ* with one point mutation Cys45Ser, Strep-tag at C-terminal; Spec® | This work |

| Strains | Genotype | Source or reference |
|---|---|---|
| ***E. coli* NZYStar** | end A1 hsd R17 (rk$^-$, mk$^+$) sup E44 thi -1 rec A1 gyr A96 rel A1 lac [F' pro A$^+$B$^+$ lac Iq ZDM15:Tn10(TcR)] | NZYTech |
| ***E. coli* One Shot$^{TM}$ TOP10** | F$^-$ mcrA Δ(mrr -hsd RMS-mcr BC) Φ80lac ZΔM15 Δlac X74 rec A1 ara D139 Δ(araleu )7697 gal U gal K rps L (StrR) end A1 nup G | Invitrogen |
| ***N. vulgaris* WT** | *Nitratidesulfovibrio vulgaris* Hildenborough ATCC 29579 | ATCC |
| ***N. vulgaris* Δ*dsrJOP*** | *N. vulgaris* WT Δ*dsrJOP* ::ΩKm | This work |
| ***N. vulgaris* Δ*dsrJ*** | *N. vulgaris* WT Δ*dsrJ* ::ΩKm | This work |
| ***N. vulgaris* Δ*dsrJ* + pMO-*dsrJ*** | *N. vulgaris* WT Δ*dsrJ* ::ΩKm + pMO-*dsrJ* | This work |
| ***N. vulgaris* Δ*dsrJ* + pMO-*dsrJ* C45A** | *N. vulgaris* WT Δ*dsrJ* ::ΩKm + pMO-*dsrJ* C45A | This work |
| ***N. vulgaris* Δ*dsrJ* + pMO-*dsrJ* C45H** | *N. vulgaris* WT Δ*dsrJ* ::ΩKm + pMO-*dsrJ* C45H | This work |
| ***N. vulgaris* Δ*dsrJ* + pMO-*dsrJ* C45S** | *N. vulgaris* WT Δ*dsrJ* ::ΩKm + pMO-*dsrJ* C45S | This work |

### Table S5. List of primers used in this study.

| # | Primer | Primer sequence (5' → 3') | Observations |
|---|---|---|---|
| **1** | *dsrJ* -upstream-forward | GCCTTTTGCTCACATTCACCGAGTTCACGGCC GACC | Construction of pMOIPAB01/02 by SLIC |
| **2** | *dsrJ* -upstream-reverse | AGCTGGCAATTCCGGGATTATGCGTCCTCCAT GCCGGGC | Construction of pMOIPAB01/02 by SLIC |
| **3** | *dsrP* -downstream-forward | GACGAGTTCTTCTGACCCCCCTTCCGACATCA AGGCC | Construction of pMOIPAB01 by SLIC |
| **4** | *dsrP* -downstream-reverse | CATTTCTGTCCTGGCTGGTGAGGGGTTCAATG TCGTCACCGC | Construction of pMOIPAB01 by SLIC |
| **5** | *dsrJ* -downstream-forward | GACGAGTTCTTCTGAATGAACAACAGCAGAA GACGATTCCTGAAGATCG | Construction of pMOIPAB02 by SLIC |
| **6** | *dsrJ* -downstream-reverse | CATTTCTGTCCTGGCTGGCCATGCAGAAGCG GCAACCG | Construction of pMOIPAB02 by SLIC |
| **7** | Kan-forward | GGAGGACGCATAATCCCGGAATTGCCAGCTG GGGC | Construction of pMOIPAB01/02 by SLIC (Kan from pSC27 forward) |
| **8** | Kan-reverse_AB01 | GATGTCGGAAGGGGGGTCAGAAGAACTCGTC AAGAAGGCGATAGAAGGCG | Construction of pMOIPAB01 by SLIC (Kan from pSC27 reverse) |
| **9** | Kan-reverse_AB02 | GCTGTTGTTCATTCAGAAGAACTCGTCAAGA AGGCGATAGAAGGCG | Construction of pMOIPAB02 by SLIC (Kan from pSC27 reverse) |
| **10** | Spec-forward | CCAGCCAGGACAGAAATGCCTCG | Construction of pMOIPAB01/02 by SLIC (Spec from pMO719 forward) |
| **11** | pUC ori-reverse | ATGTGAGCAAAAGGCCAGCAAAAGGC | Construction of pMOIPAB01/02 by SLIC (pUCori from pMO719 forward) |
| **12** | *dsrJ* -forward | TGCAGTCCCAGGAGGTACCATATGTATAACG GCAAGTACATCATCCCCGG | Construction of pMO-dsrJ by SLIC |
| **13** | *dsrJ* -reverse | CGTTTATTTTTCGAACTGCGGGTGGCTCCACT GGTTCCCCCTCGGAGCGAC | Construction of pMO-dsrJ by SLIC |
| **14** | pUC ori/Spec-forward | GGGAACTGCCAGGCATCAAATAAAACGAAA GGCTCAGTCG | Construction of pMO-dsrJ by SLIC (pUCori and Spec from pMOIP3 forward) |
| **15** | pUC ori/Spec-reverse | CTTGCCGTTATACATATGGTACCTCCTGGGAC TGCATTG | Construction of pMO-dsrJ by SLIC (pUCori and Spec from pMOIP3 reverse) |
| **16** | pBG1-forward | CAGTGGAGCCACCCGCAGTTCGAAAAATAAA CGAAAAGACCCGATCATGAAGGGG | Construction of pMO-dsrJ by SLIC (pBG1 from pMOIP3 forward) |
| **17** | pBG1-reverse | CGACTGAGCCTTTCGTTTTATTTGATGCCTGG CAGTTCCC | Construction of pMO-dsrJ by SLIC (pBG1 from pMOIP3 reverse) |
| **18** | *dsrJ* C45A-forward | GCC GGG GAG AAG GAA GCC ATC GAG CCC GTG GGC TAC | Replacement of Cys for Ala on pMO-*dsrJ* by Site-Directed Mutagenesis |
| **19** | *dsrJ* C45A-reverse | GTA GCC CAC GGG CTC GAT GGC TTC CTT CTC CCC GGC | Replacement of Cys for Ala on pMO-*dsrJ* by Site-Directed Mutagenesis |
| **20** | *dsrJ* C45H-forward | GCC GGG GAG AAG GAA CAT ATC GAG CCC GTG GGC TAC | Replacement of Cys for His on pMO-*dsrJ* by Site-Directed Mutagenesis |
| **21** | *dsrJ* C45H-reverse | GTA GCC CAC GGG CTC GAT ATG TTC CTT CTC CCC GGC | Replacement of Cys for His on pMO-*dsrJ* by Site-Directed Mutagenesis |
| **22** | *dsrJ* C45S-forward | GCC GGG GAG AAG GAA AGC ATC GAG CCC GTG GGC TAC | Replacement of Cys for Ser on pMO-*dsrJ* by Site-Directed Mutagenesis |
| **23** | *dsrJ* C45S-reverse | GTA GCC CAC GGG CTC GAT GCT TTC CTT CTC CCC GGC | Replacement of Cys for Ser on pMO-*dsrJ* by Site-Directed Mutagenesis |

**Movie S1.**

**A proposed mechanism of DsrC-trisulfide reduction by DsrMKJOP.** The reaction begins with quinol binding to DsrP and its oxidation, during which electrons are transferred through the entire DsrMKJOP complex to DsrC, and protons are released to the periplasm. The DsrC-trisulfide initially docks onto DsrK, where trisulfide hydrolysis is promoted by Phe258$^{DsrK}$. Subsequently, the C-terminal arm of DsrC extends toward the non-cubane cluster within DsrK, positioning the substrate near the cluster and priming the system for sulfur reduction. Membrane boundaries shown at the beginning of the movie were calculated using the PPM 3.0 web server [14].
